# Behavioural changes in aposematic *Heliconius melpomene* butterflies in response to their predatory bird calls

**DOI:** 10.1101/2023.10.24.563859

**Authors:** Sushant Potdar, Madhuri Dinakar, Erica L. Westerman

## Abstract

Prey-predator interactions have resulted in the evolution of many anti-predatory traits. One of them is the ability for prey to listen to predators and avoid them. Although prey anti-predatory behavioural responses to predator auditory cues are well described in a wide range of taxa, studies on whether butterflies change their behaviours in response to their predatory calls are lacking. *Heliconius* butterflies are unpalatable and form Müllerian mimicry rings as morphological defence strategies against their bird predators. Like many other butterflies in the *Nymphalidae* family, *Heliconius* butterflies possess auditory organs, which are hypothesized to have evolved to assist with predator detection. Here we test whether *Heliconius melpomene* change their behaviour in response to their predatory bird calls by observing the behaviour of male and female *H. m. plessini* exposed to calls of *Heliconius* avian predators: rufous-tailed jacamar, migratory Eastern kingbird, and resident tropical kingbird. We also exposed them to the calls of the toco toucan, a frugivorous bird as a control bird call, and an amplified greenhouse background noise as a noise control. We found that individuals changed their behaviour in response to jacamar calls only. Males increased their walking and fluttering behaviour, while females did not change their behaviour during the playback of the jacamar call. Intersexual behaviours like courtship, copulation, and abdomen lifting did not change in response to bird calls. Our findings suggest that despite having primary predatory defences like toxicity and being in a mimicry ring, *H. m. plessini* butterflies changed their behaviour in response to predator calls. Furthermore, this response was predator specific, as *H. m. plesseni* did not respond to either the Eastern kingbird or the tropic kingbird calls. This suggests that *Heliconius* butterflies may be able to differentiate predatory calls, and potentially the birds associated with those calls.

**Highlights:**

1. Many prey animals change their behaviour in response to their predator’s calls.
2. Whether butterflies alter behaviour in response to bird predator calls is unknown.
3. We show that *Heliconius melpomene* change behaviour in response to jacamar calls.
4. Males increased walking and fluttering, but did not alter courting behaviour.
5. *H. melpomene* did not respond to predatory Eastern kingbird or tropical kingbird calls.

## Introduction

Predation is a ubiquitous interspecific interaction in almost all ecosystems and can be a strong evolutionary force for the emergence and selection of prey anti-predatory strategies that increase survival (Lind & Cresswell, 2005). Anti-predatory strategies are widespread in prey animals and can be morphological or behavioural. Morphological strategies include aposematism, chemical toxicity, and crypsis (Rojas et al., 2019; Vallin et al., 2006), while behavioural anti-predatory strategies include active evasion of predatory attacks, and behaviours that decrease detection (Palmer & Packer, 2021). Anti-predatory strategies can also be a combination of both morphological and behavioural strategies such as the deimatic displays in mountain katydid *Acripeza reticulata* and swallowtail butterflies (Olofsson et al., 2012; Umbers & Mappes, 2015).

The most common anti-predatory strategies are behavioural responses to detection and active attacks by predators. These behavioural responses help prey escape predation, either in the absence of morphological defences, or as a combination with morphological defences, and can be highly variable across species, within species, and between sexes (Apfelbach et al., 2005; Lind & Cresswell, 2005). Some species, such as desert isopods (*Hemilepistus reaumuri*), freeze and retreat inside their burrows upon smelling their predator’s scent (Zaguri & Hawlena, 2020); while others, such as male tree lizards (*Urosaurus ornatus*) actively escape by fleeing after detecting their predators (Thaker et al., 2009). Anti-predatory behaviours can also differ within species in response to different predators, as illustrated by red squirrels (*Tamiasciurus hudsonicus*), which have different alarm calls for avian predators and ground predators (Greene & Meagher, 1998). Anti-predatory behavioural responses can also be sex-specific, either due to the inherent sex-specific differences in physiology and behaviour, or due to the increased vulnerability of predation during intraspecific sexual behaviours (Curlis et al., 2016; Edomwande & Barbosa, 2020; Lea & Blumstein, 2011; Sitvarin & Rypstra, 2012; Wormington & Juliano, 2014). Both males and females are known to alter their courtship and mating behaviours under predation risk (Acharya & McNeil, 1998; Torsekar et al., 2019). In wolf spider (*Schizocosa ocreata*), males cease courtship behaviour after detecting predatory birds’ calls and take longer to return to courting compared to non-threatening control sounds (Lohrey et al., 2009), while female túngara frogs (*Physalaemus pustulosus*) approach calling males more cautiously when exposed to bat wingbeat sounds (Bernal et al., 2007). The cost of predation during intraspecific sexual behaviours such as courtship and copulation is high, forcing individuals to switch from sexually oriented behaviours to survival behaviours.

Detecting and recognizing predatory cues are necessary for active predator avoidance behaviours, and these cues can either be visual, chemical, vibrational, or auditory. Auditory cues play a vital role in predator avoidance by prey animals, from invertebrates (Faure & Hoy, 2000; Jacobs et al., 2008; Lohrey et al., 2009; Prakash et al., 2021; Rosen et al., 2009; Triblehorn et al., 2008) to vertebrates (Bernal et al., 2007; Cantwell & Forrest, 2013; Deecke et al., 2002). In Lepidoptera (moths and butterflies), anti-predatory behaviours in moths to predators’ auditory cues have been extensively studied under various ecological contexts. Moths have evolved hearing to detect bat echolocation calls and avoid bat predation by performing aerial manoeuvrers and jamming echolocation calls (Conner & Corcoran, 2012). Both male and female moths also reduce sexual activity under bat predation pressure (Acharya & McNeil, 1998; Edomwande & Barbosa, 2020).

While moths are particularly well known for their hearing ability and anti-predator behaviours, butterflies, their day-flying relatives, are also known to have auditory organs, which may be sensitive to predator sounds (Lane et al., 2008). In particular, many species in the family *Nymphalidae* possess auditory organs on their wings, such as the blue morpho *Morpho peleides* (Lane et al., 2008; Lucas et al., 2009; Mikhail et al., 2018), common wood nymph *Cercyonis pegala* (Sun et al., 2018), the owl butterfly *Caligo eurilochus* (Lucas et al., 2014), butterflies from the genus *Erebia* (Ribarič & Gogala, 1996), and *Heliconius* butterflies (Swihart, 1967). However, unlike moths, it is generally unknown whether butterflies that possess auditory organs change their behaviour in response to their predator’s vocalizations. In this study, we used a butterfly from the genus *Heliconius* to test whether these butterflies change their behaviour in response to their predator’s vocalizations.

*Heliconius* butterflies (Family Nymphalidae), found in North, Central, and South America, are toxic, unpalatable, display aposematic colouration, form Müllerian mimicry rings, and roost communally to avoid bird and bat predation (Engler-Chaouat & Gilbert, 2007; Finkbeiner et al., 2012; Mallet & Gilbert, 1995; Pinheiro De Castro et al., 2019). Despite these anti-predatory strategies, *Heliconius* butterflies are vulnerable to predation by specialist bird predators, as well as by naïve generalist predatory birds; and their mortality is higher when young birds are learning which butterfly species are toxic and should be avoided (Chai, 1986; Langham, 2004, 2006; Pinheiro, 1996; Pinheiro & Cintra, 2017). Hence, it may be evolutionarily advantageous for *Heliconius* butterflies to detect the presence of their bird predators and change their behaviours to reduce detection, despite having multiple anti-predatory strategies. One possible way these butterflies could detect the presence of their bird predators is by using avian vocalization cues, which are often species specific (Lane et al., 2008; Lucas et al., 2009; Mikhail et al., 2018). In *Heliconius* butterflies, hearing organs located at the base of the hindwing with peak sensitivity between 0.5 to 4 KHz at 70-90 dB pressure have been described (Swihart 1967). However, the hypothesis that *Heliconius* butterflies change their behaviour in response to their predatory birds’ vocalizations has never been tested.

In this study, we tested whether *Heliconius melpomene plessini* butterflies change their behaviour in response to the vocalizations of their known bird predators. We first tested butterfly response to the vocalizations of two predatory birds with disparate calls as well as the vocalization of a frugivorous bird, to assess whether *H. m. plesseni* butterflies respond to both predator bird calls and calls of non-predatory birds. After answering that question, we then tested the response of *H. m. plessini* butterflies to predators that differ in annual patterns of predation (year-round resident or migratory), to assess whether strength of *H. m. plesseni* response is associated with degree of annual avian predator exposure. During both these experiments, we also tested whether intraspecific sexual behaviours like male courtship and female acceptance/rejection behaviours changed in response to *H. m. plessini’s* bird predatory calls.

## Materials and Methods

### Study species husbandry

*Heliconius melpomene* (Order: *Lepidoptera*, Family: *Nymphalidae*), is native to Central and South America. The subspecies *H. m. plessini* is found in the mountainous forests of Ecuador and Peru in South America (Hines et al., 2011). Live pupae of *H. m. plessini* were shipped from Ecodecision Heliconius Works in Quito, Ecuador to the University of Arkansas Biology greenhouse facility in Fayetteville AR, USA, where they were maintained at an average temperature of 27°C, average relative humidity of 70% and a 13:11 hour L:D cycle, to mimic summer tropical conditions. All pupae were hung and housed in mesh BioQuip cages (34.29 x 34.29 x 60.96 cm, Rancho Dominguez, CA, U.S.A.) until their eclosion in the greenhouse facility. Newly eclosed individuals were sexed and tagged with a unique number with a silver metallic permanent marker (SHARPIE 39108PP) and placed in sex-specific mesh BioQuip cages (60.96 x 60.96 x 142.24 cm) with *ad libitum* BIRDS choice butterfly nectar (Birdschoice, Chilton, WI, USA) and pollen from *Lantana spp* flowers. Marking butterflies with a marker does not have long term effects on their behaviour and lifespan (Gall, 1984). Female *H. m. plessini* were housed with females of two other subspecies, *H. m. malleti* and *H. m. rosina* while male *H. m. plessini* were housed on their own. Both the male and female cages were visually isolated from the opposite sex and had no more than 15 individuals in each sex specific cage at any point in time.

### Bird calls and control treatments

We used the calls of four different bird species during our experiments: three *Heliconius* predators and one frugivore as a control species. Our predatory bird species were the rufous-tailed jacamar (*Galbula ruficauda*), Eastern kingbird (*Tyrannus tyrannus*), and tropical kingbird (*Tyrannus melancholicus*) (Pinheiro, 1996, 2011; Pinheiro & Cintra, 2017). We used the non-predatory toco toucan (*Ramphastos toco*) call to test if *H. m. plessini* respond to bird calls in general, and amplified greenhouse background noise as a random noise control. We chose toco toucan as our control bird call because it is a non-predatory frugivorous bird found in the same habitat as our focal butterflies and has a naturally loud call. Playback recordings of the four bird calls (rufous-tailed jacamar, Eastern kingbird, tropical kingbird and toco toucan) with minimal disturbance from background animals were downloaded from Xeno-Canto (Xeno-Canto Foundation; www.xeno-canto.org) (for sonograms of all calls see Supplementary Figure 2). These bird calls were characterized as ‘songs’ in the original files uploaded on Xeno-Canto. All the bird calls contain elements within previously reported *Heliconius* hearing frequency 1-4 KHz (Swihart, 1967), though the main components of the kingbird calls’ are just outside that range at 5 KHz (Supplementary Figure 2).

The University of Arkansas butterfly facility has constant and continuous noise generated by fans and misters which were measured at 65 dB near the behavioural watch cage using an android sound meter application (Sound Meter-Decibel and noise Meter). To account for any butterfly behavioural responses to this background noise, or to loud noises in general, we recorded the greenhouse noise using the android voice recorder application (Voice Recorder, version 3 (42.0)) and used this recording in behavioural assays as a greenhouse background noise control. During the behavioural assays, the calls of rufous-tailed jacamar (76 dB), Eastern kingbird (79 dB), tropical kingbird (80 dB), toco toucan (80 dB) and the greenhouse background noise control (77 dB) were played at 10-15 dB louder than the actual greenhouse background noise. Bird calls in forests are always against a naturally generated background noise (by other animals; leaves rustling, waterfalls, and streams). While our constant greenhouse background noise is admittedly different from that of a forest, the presence of background noise broadly emulates such sounds generated in the forest. All calls were standardized to one minute long .mp3 files.

### Behavioural Assays

All behavioural assays were conducted between 11:00 AM and 2:00 PM, when *H. melpomene* are most active in our greenhouse (Rather et al., 2022). We conducted behavioural assays using 3-15-day-old males and females in a large behavioural cage (60.96 x 60.96 x 142.24 cm). In each assay, we used one male and one female and acclimated them in the behavioural cage for 15 minutes with a JBL_®_ Flip 4 portable blue-tooth speaker (Harman) and a *Lantana spp.* plant. We used both a male and a female in our behavioural assay to determine whether predatory bird calls had an effect on intersexual behaviours (*courtship, abdomen lifting, copulation and, sitting near*) in addition to any other types of behaviour (*wing fluttering, antennae wiggling*, *basking, flying, resting, walking*). After a 15-minute acclimation period, we recorded all the behaviours performed by the two individuals in the assay for 15 minutes prior to any playback calls. We then played one of the bird calls or the control greenhouse background noise using a JBL_®_ Flip 4 portable blue-tooth speaker from the observer’s phone (Google Pixel), placed inside the behavioural cage for 1 minute and recorded the behaviours of the two individuals during the playback of the call/background noise. After the playback, we recorded the behaviours of the two individuals for an additional 14 minutes (Supplementary Figure 1). We recorded the frequency of *fluttering* and *antenna wiggle* behaviours and the frequency and duration of *basking, flying, resting, walking, courtship, copulation, abdomen lifting,* and *sitting near each other* behaviours throughout the entire 30-minute observational period.

We defined behaviours for *H. m. plessini* as follows: *fluttering-* opening and closing of wings either while resting or walking; *antenna wiggle*- movement of antennae at 45° angle in any direction (Robertson et al., 2020); *basking*- individuals sitting with wings partially or fully open; *flying*- movement from one point to another in the air using rapid wing flaps; *resting*- individuals sitting with wings fully closed (Rather et al. 2022); *walking*- movement from one point to another along the substrate using the legs; *courtship*- sequences of behaviours where males hover, land and rapidly flap their wings next to females, and bend their abdomen to initiate copulation (Klein & De Araújo, 2010); *copulation*- where both male and female are mating; *abdomen lifting*- raising the abdomen at an angle from the normal resting body axis, usually performed by females as a courtship rejection behaviour (Chouteau et al., 2017); *sitting near each other*- where both individuals are resting or basking within one wingspan from each other (Robertson et al. 2020).

We used Spectator Go (BIOBSERVE, Fort Lee, NJ, USA) software on an Apple iPad (1^st^ generation) to manually record the frequency and duration of behaviours performed by the two individuals during the assay. This software enables the observer to record user defined behaviours in real time, separately for the two individuals, without instantly visualizing quantities during the recording, and has been used in previous studies to observe and record butterfly behaviours (Rather et al., 2022; Robertson et al., 2020; Westerman et al., 2014). To reduce observer bias, only one observer recorded all the behaviours in this study. We did not use a video camera to record behaviours as some butterfly inter-individual interactions are minute and nuanced happening at a close range, while others occupy the full three-dimensional flight area of the cage, and simultaneously capturing both of these types of behaviours is challenging for a stationary camera, but relatively straightforward for a trained human observer. Within each experiment, we tested each male-female pair with all calls with at least 24 hours between each call assay, and randomized the order of calls for each pair. If either of the butterflies in the pair died between the assays, then those pairs were eliminated from being tested for the remaining calls.

### Experiment 1: Do H. m. plesseni butterflies behaviourally respond to predator bird calls

To test whether *H. m. plessini* butterflies respond to their avian predator calls or to other birds or loud random noises in general, we subjected the butterflies to four call treatments in this experiment: rufous-tailed jacamar (N=22 pairs), Eastern kingbirds (N=22 pairs), toco toucan (N=22 pairs) and greenhouse background noise control (N=18 pairs) using the behavioural assay described above with the calls randomized. We conducted Experiment 1 from February 2019 to March 2020.

### Experiment 2: Does predator residence status influence butterfly response to bird call

Due to the results of Experiment 1 (see below), we conducted a follow up experiment to test whether predator residence status (migratory or present year-round) influenced likelihood of *H. m. plessini* butterflies changing their behaviour in response to predator call. For this experiment, we used the calls of the resident tropical kingbird and the migratory Eastern kingbird, as they have vocalizations in the same auditory frequencies, and are more closely related than the Eastern kingbird and rufous-tailed jacamar. We subjected butterflies to three call treatments: resident tropical kingbird (N=23 pairs), migratory Eastern kingbird (N=22 pairs) and the control greenhouse background noise control (N=25 pairs) using the same behavioural assay as Experiment 1, as described above. We conducted Experiment 2 from August to December 2021. We conducted the same statistical analyses for both Experiment 1 and Experiment 2, albeit separately.

### Statistical analyses

We downloaded the data from Spectator Go software and converted them into .csv files. Each file consisted of approximately 15 minutes of data, and each assay had four files (15 minutes before, and during plus after call for male and female separately). Each bout of behaviour was recorded separately by the software for the 10 behaviours described above. A *de novo* python code (supplementary material 2) was written to add each bout of a behaviour and provide the total time spent performing that particular behaviour. This way, we got the total time spent by an individual butterfly performing behaviours for the whole assay. Further, we manually extracted the behavioural states before and after the start and end of calls, as well as extracted the behaviours performed a minute before, during and after the calls. We performed three separate analyses for each experiment: behavioural state change between before and after the start and end of calls; short term (1 minute) changes in behaviours between before, during, and after calls; and long term (14 minutes) changes in behaviour before and after calls.

To determine whether butterflies changed their behavioural state in response to bird call, we compared the behaviours performed across three time points of an assay: 1) before vs after the start of call; 2) before vs after the end of call; and 3) before start vs after end of the call. We used generalized linear mixed models (GLMM) with change in behaviour between the above time points (yes or no) as the response variable, treatment (calls), and sex (male or female) as fixed predictor variables, and the order of calls as a random predictor variable. We later used a pairwise Fisher’s test to determine if the proportion of individuals that changed their behaviours were similar or different between the treatments (bird calls and noise control).

To test if the frequency and duration of short-term behaviours changed during and after a call compared to before a call, we extracted the frequency of *fluttering* and *antenna wiggle* and duration of the other eight behaviours for the minute before, minute during, and minute after the call. We performed Principal Component Analysis (PCA) for the behavioural data during these three minutes, to identify the correlation between different behaviours and identify new composite behavioural variables. We removed *abdomen lifting* from the male data set and *courtship* from the female data set as males and females respectively did not perform these behaviours. We fit a linear mixed model (LMMs), followed by an ANOVA, with treatment (bird call), state (before, during, and after call) and their interaction as fixed predictor variables, the order of the calls as a random predictor variable, and the first three principal components as the response variables. Further, we performed a Tukey HSD test to determine the pairwise differences between different combinations of treatment (bird call) and state (before, during, and after call). Later, we tested whether male *courtship, sitting near each other,* female *abdomen lifting*, *copulation* behaviours changed in response to bird calls by fitting LMMs followed by an ANOVA, with the same predictor variables. We ran these models for males and females separately, as males and females performed different behaviours. We also performed these analyses separately for experiments 1 and 2.

Next, to test if there was a prolonged long term response of butterfly behaviour to the bird calls, we extracted the frequency of *fluttering* and *antenna wiggle* and duration of the other eight behaviours for the 14 minutes before the call and the 14 minutes after the call, and performed a PCA for these 28 minutes, again removing *abdomen lifting* behaviour from male data set and *courtship* behaviour from the female data set. We fit LMM, followed by an ANOVA, with treatment (bird call), state (before, and after call), and their interaction as the fixed predictor variables, the order of the calls as a random predictor variable, and the first three principal components as the response variables for each sex. Further, we performed a Tukey HSD test to determine the pairwise differences between different combinations of treatment (bird call) and state (before, and after call). We also tested whether male *courtship, sitting near,* female *abdomen lifting*, *copulation* behaviours changed in response to bird calls by fitting LMMs followed by an ANOVA, with the same predictor variables. We again ran these models for both males and females separately, and performed these analyses separately for experiments 1 and 2.

All statistical analyses were run using R version 4.3.0 (R Core Team, 2023). All plots were generated using *ggplot2* (Wickham, 2016) package.

### Ethical Note

All butterflies used in this study were maintained in climate-controlled greenhouse conditions similar to those of their natural habitat, as stated in the U.S. Department of Agriculture, Animal and Plant Health Inspection Service permits P526P-17-00343 and P526P-20-00417. Before and after the assays, all butterflies were maintained in cages with *ad libitum* food (nectar and flowering *Lantana spp.* plants for pollen). After the assays, they were moved to breeding cages with *ad libitum* food, where they were kept until natural death. No butterflies were sacrificed for the purpose of this study.

## Experiment 1 Results

### *H. m. plessini* immediately changed their behavioural state in response to the rufous-tailed jacamar call

*H. m. plessini* butterflies immediately changed their behavioural state when the rufous-tailed jacamar call started (χ2= 16.03, p<0.01; Supplementary Figure 3A; Supplementary Table 1, 2), when the jacamar call stopped (χ2= 17.47, p<0.001; Supplementary Figure 4A; Supplementary Table 3, 4), and when compared between before the call started versus after the call ended (χ2= 27.12, p<0.001, Table 1, 2, Figure 1A). They did not significantly change their behavioural state in response to any other bird call, or in response to the noise control (Supplementary Table 1, 2, 3, 4; Table 1, 2). We did not find an effect of sex on the change in behavioural state when the calls started, when the calls stopped, or when compared between before the calls started versus after the calls ended nor was there an effect of call order on butterfly response (Supplementary Table 1, 3; Table 1).

**Figure 1:**
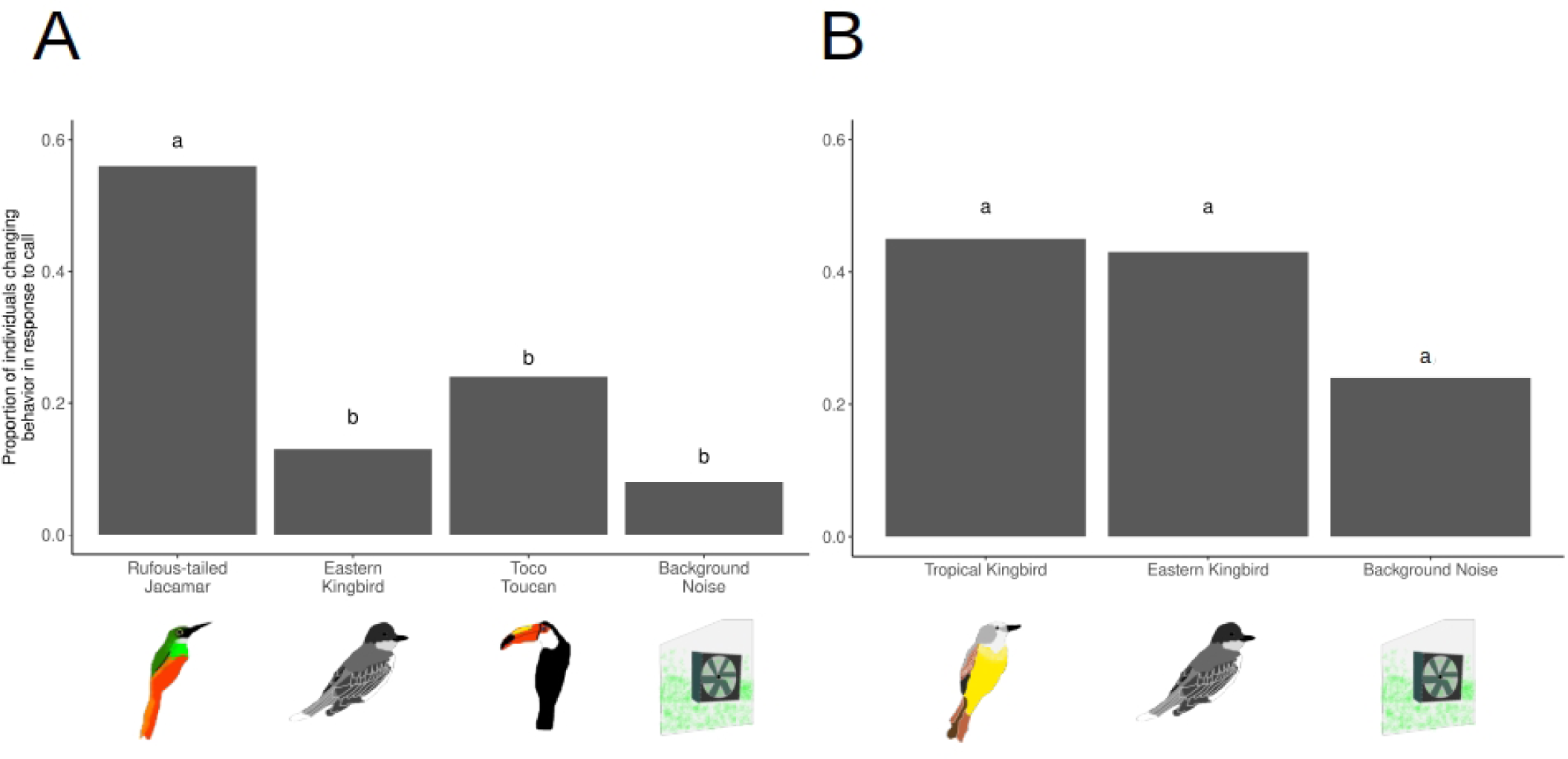
Proportion of *H. m. plessini* individuals changing behaviour in response to calls (between before start and after end of calls) for A) experiment 1; B) experiment 2; Different letters on each bars indicate statistical significance at p<0.05.

**Table 1:**
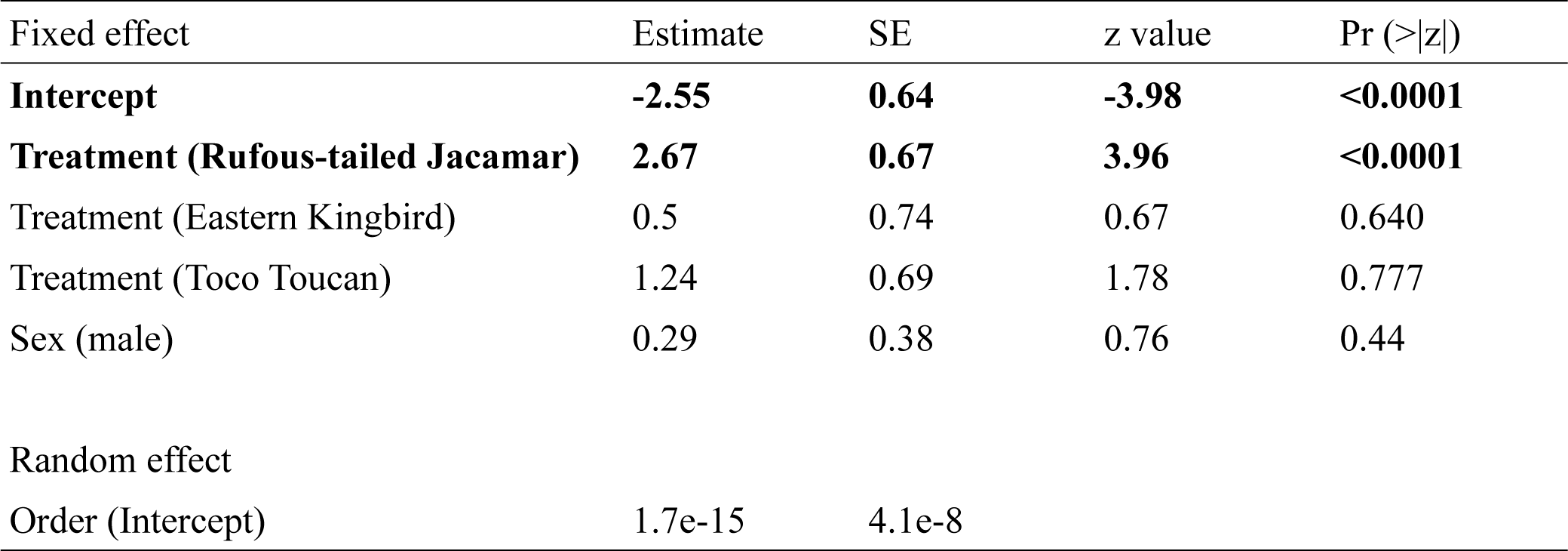
GLMM results on the effect of treatment (calls) and sex on proportion of butterflies changing their behaviour in response to calls. p<0.05 are bolded.

**Table 2:**
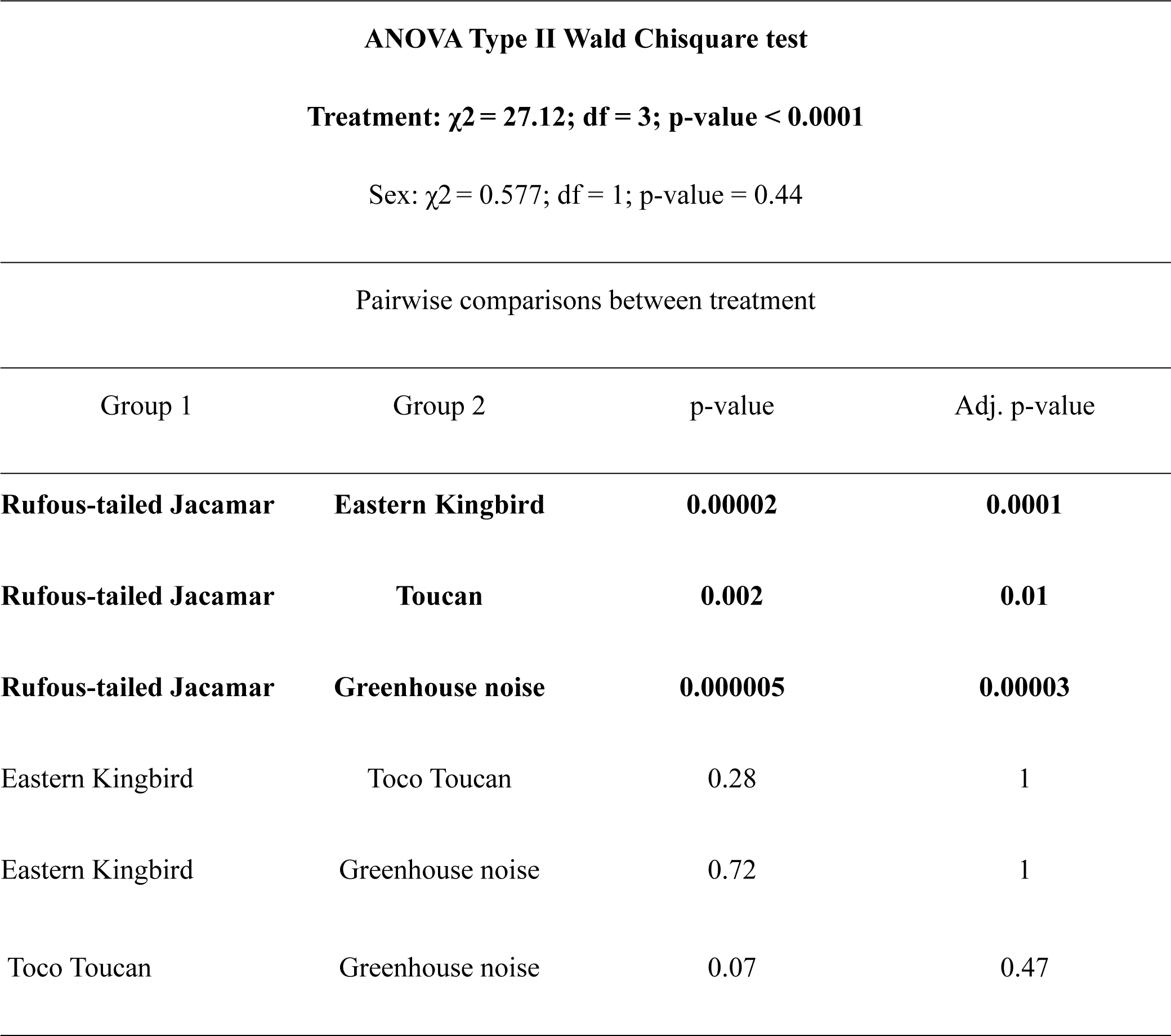
Pairwise differences in the proportion of individuals changing their behavioural state in response to calls in experiment 1. p<0.05 are bolded.

| ANOVA Type II Wald Chisquare test |  |  |  |
| --- | --- | --- | --- |
| Treatment: $\chi^2 = 27.12$; df = 3; p-value < 0.0001 | | | |
| Sex: $\chi^2 = 0.577$; df = 1; p-value = 0.44 | | | |
| Pairwise comparisons between treatment |  |  |  |
| Group 1 | Group 2 | p-value | Adj. p-value |
| **Rufous-tailed Jacamar** | **Eastern Kingbird** | **0.00002** | **0.0001** |
| **Rufous-tailed Jacamar** | **Toucan** | **0.002** | **0.01** |
| **Rufous-tailed Jacamar** | **Greenhouse noise** | **0.000005** | **0.00003** |
| Eastern Kingbird | Toco Toucan | 0.28 | 1 |
| Eastern Kingbird | Greenhouse noise | 0.72 | 1 |
| Toco Toucan | Greenhouse noise | 0.07 | 0.47 |

### *H. m. plessini* males increased their walking and fluttering behaviour during the playback of the rufous-tailed jacamar call

When combining the behavioural data for the 3 minutes before, during, and after each call in a PCA for each sex, male PC2 values were higher during the rufous-tailed jacamar call compared to before and after the rufous-tailed jacamar call, and compared to before, during, and after the Eastern kingbird, toco toucan, and greenhouse background noise (ANOVA, F= 2.336, Df= 6, p= 0.0328; Figure 2C; Table 3; Supplementary Table 7; see Supplementary Table 5 for PCA loadings). There was no effect of any of the bird calls or greenhouse background noise control on male PC1 (Figure 2A; Table 3; Supplementary Table 7, 8), male PC3 (Table 3; Supplementary Table 7), female PC1 (Figure 2B; Table 3; Supplementary Table 7; see Supplementary Table 6 for PCA loadings), female PC2 (Figure 2D; Table 2; Supplementary Table 7), or female PC3 (Table 2; Supplementary Table 7).

**Figure 2:**
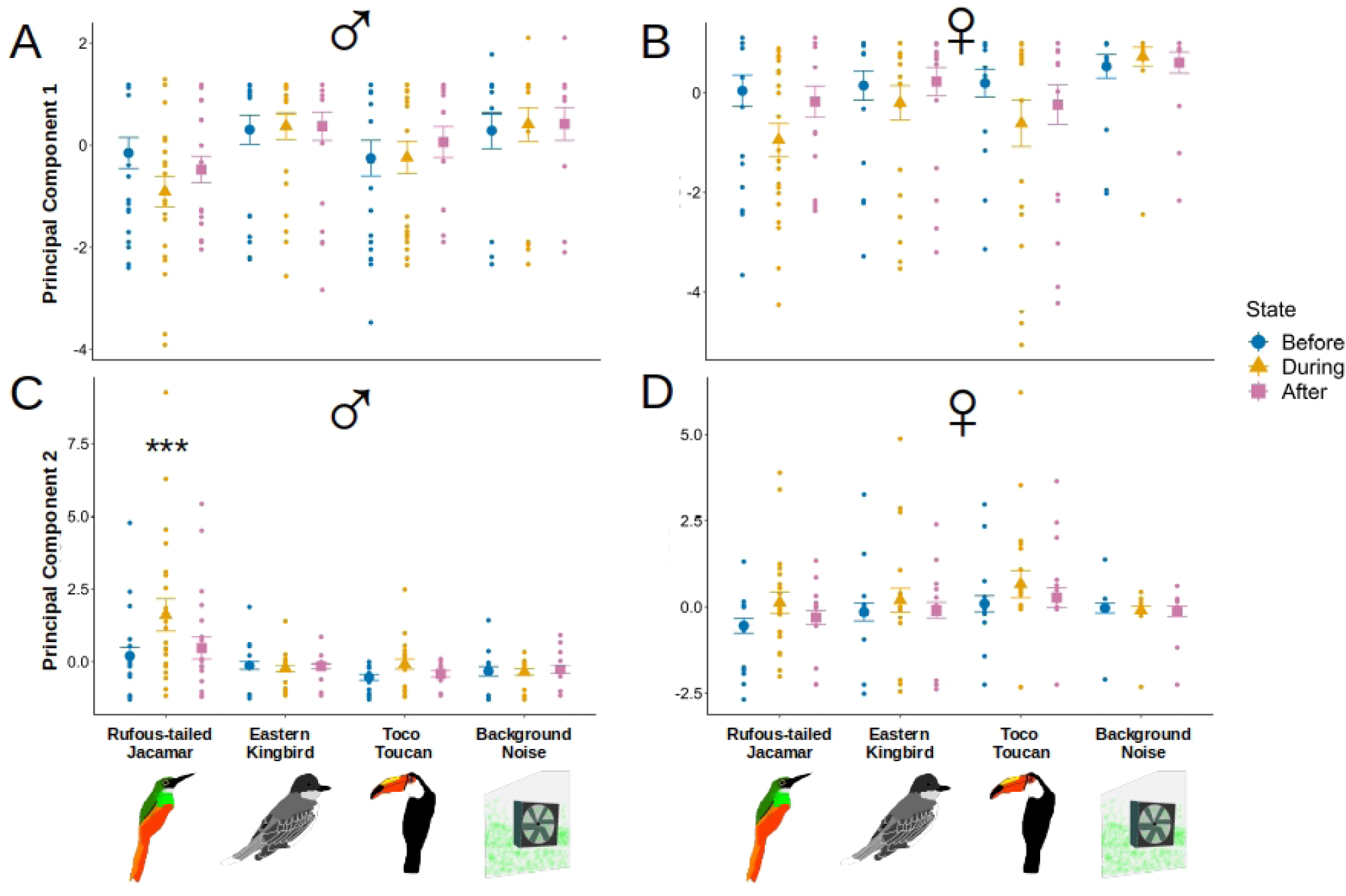
Mean ± SE of principal component variables for male and female *H. m. plessini* for a minute before, during and after calls. A) PC 1 in males for experiment 1; B) PC 1 in females for experiment 1; C) PC 2 in males for experiment 1; D) PC 2 in females for experiment 1. *** indicates significance with p<0.0001.

**Table 3:** Effect of treatment (rufous-tailed jacamar, Eastern kingbird, toco toucan and greenhouse background noise calls), state (one minute before, during and after call) and their interaction on male and female PC1 and PC2 in experiment 1. p<0.05 are bolded.

| ♂ | AIC | Df | F value | Pr (>F) | ♀ | AIC | Df | F value | Pr (>F) |
| --- | --- | --- | --- | --- | --- | --- | --- | --- | --- |
| PC1 | 884 |  |  |  | PC1 | 910 |  |  |  |
| **Treatment** |  | **3** | **5.887** | **0.0006** | **Treatment** |  | **3** | **5.090** | **0.001** |
| State |  | 2 | 0.447 | 0.640 | State |  | 2 | 2.794 | 0.063 |
| Treatment*State |  | 6 | 0.540 | 0.777 | Treatment*State |  | 6 | 0.732 | 0.624 |
| PC2 | 781 |  |  |  | PC2 | 805 |  |  |  |
| **Treatment** |  | **3** | **13.900** | **2.2e-08** | **Treatment** |  | **3** | **2.759** | **0.043** |
| **State** |  | **2** | **3.649** | **0.0275** | State |  | 2 | 2.498 | 0.084 |
| **Treatment\*State** |  | **6** | **2.336** | **0.0328** | Treatment*State |  | 6 | 0.373 | 0.896 |
| PC3 | 787 |  |  |  | PC3 | 745 |  |  |  |
| Treatment |  | 3 | 1.432 | 0.234 | Treatment |  | 3 | 2.017 | 0.112 |
| State |  | 2 | 0.856 | 0.426 | State |  | 2 | 1.071 | 0.344 |
| Treatment*State |  | 6 | 0.697 | 0.652 | Treatment*State |  | 6 | 0.577 | 0.748 |
| Courtship | 239 |  |  |  | Copulation | 1692 |  |  |  |
| Treatment |  | 3 | 0.984 | 0.401 | **Treatment** |  | **3** | **2.765** | **0.042** |
| State |  | 2 | 0.999 | 0.370 | State |  | 2 | 0 | 1 |
| Treatment*State |  | 6 | 0.984 | 0.437 | Treatment*State |  | 6 | 0 | 1 |
| Sitting near other | 1659 |  |  |  | Abdomen lift |  |  |  |  |
| **Treatment** |  | **3** | **3.500** | **0.016** | Treatment |  | 3 | 0.922 | 0.431 |
| State |  | 2 | 0.011 | 0.988 | State |  | 2 | 0.997 | 0.370 |
| Treatment*State |  | 6 | 0.003 | 1 | Treatment*State |  | 6 | 0.922 | 0.480 |

### *H. m. plessini* males and females had no long-term changes in behaviour in response to calls

When combining the behavioural data for the 14 minutes before and after each call in a PCA for each sex, there was no effect of any of the bird calls or greenhouse background noise control on male PC1 (Supplementary Figure 5A; Supplementary Table 10, 11; see Supplementary Table 8 for PCA loadings), male PC2 (Supplementary Figure 5C; Supplementary Table 10, 11), male PC3 (Supplementary Table 10, 11); female PC1 (Supplementary Figure 5B; Supplementary Table 10, 11; see Supplementary Table 9 for PCA loadings), female PC2 (Supplementary Figure 5D; Supplementary Table 10, 11), or female PC3 (Supplementary Table 10, 11).

### No effect of predatory bird calls on *H. m. plessini* intersexual behaviours

Male *courtship, sitting near each other,* female *abdomen lifting, copulation* behaviours did not have short-term or long-term changes in response to any bird calls (Table 3; Supplementary Table 10).

## Experiment 2 results

While there are a number of hypotheses as to why *H. m. plessini* did not change their behaviour in response to the migratory Eastern kingbird calls, but did change their behaviours in response to the resident jacamar calls, two we found particularly interesting were 1) that jacamars are year round residents while Eastern kingbirds are migratory; and 2) jacamars and Eastern kingbirds have different call frequencies (Hz ranges). To test the hypothesis that residence status is driving *H. m. plesseni* behavioural response while holding call frequency (Hz) constant, we then tested whether *H. m. plessini* butterflies changed their behaviour in response to the resident tropical kingbird call compared to the migratory Eastern kingbird call in Experiment 2, as these two kingbird species have vocalizations in the same auditory frequency ranges (Supplementary Figure 2).

### Residence status of kingbirds did not change *H. m. plessini* behavioural state

We found that *H. m. plessini* butterflies did not change their behavioural state when either of the resident or migratory kingbird calls or greenhouse background noise started (Supplementary Figure 3B; Supplementary Table 12, 13), when either of the kingbird calls or greenhouse background noise stopped (Supplementary Figure 4B, Supplementary Table 14, 15), and when compared between before the kingbird calls started versus after the kingbird calls ended, as well as between before the start and after the end of greenhouse background noise (Figure 1B; Supplementary Table 16, 17). We did not find an effect of sex on the change in behavioural state when calls started, when the calls stopped, and when compared between before the calls started versus after the calls ended nor was there an effect of call order on butterfly response (Supplementary Table 12, 14, 16).

### Residence status of kingbirds did not change short-term *H. m. plessini* behaviours

When combining the behavioural data for the 3 minutes before, during, and after each call in a PCA for each sex, there was no effect of any kingbird calls or greenhouse background noise control on male PC1 values (Figure 3A; Supplementary Table 20, 21; see Supplementary Table 18 for PCA loadings), on male PC2 (Figure 3C; Supplementary Table 20, 21), male PC3 (Supplementary Table 20, 21), female PC1 (Figure 3B; Supplementary Table 20, 21; see Supplementary Table 19 for PCA loadings), female PC2 (Figure 3D; Supplementary Table 20, 21), and female PC3 (Supplementary Table 20, 21) values.

**Figure 3:**
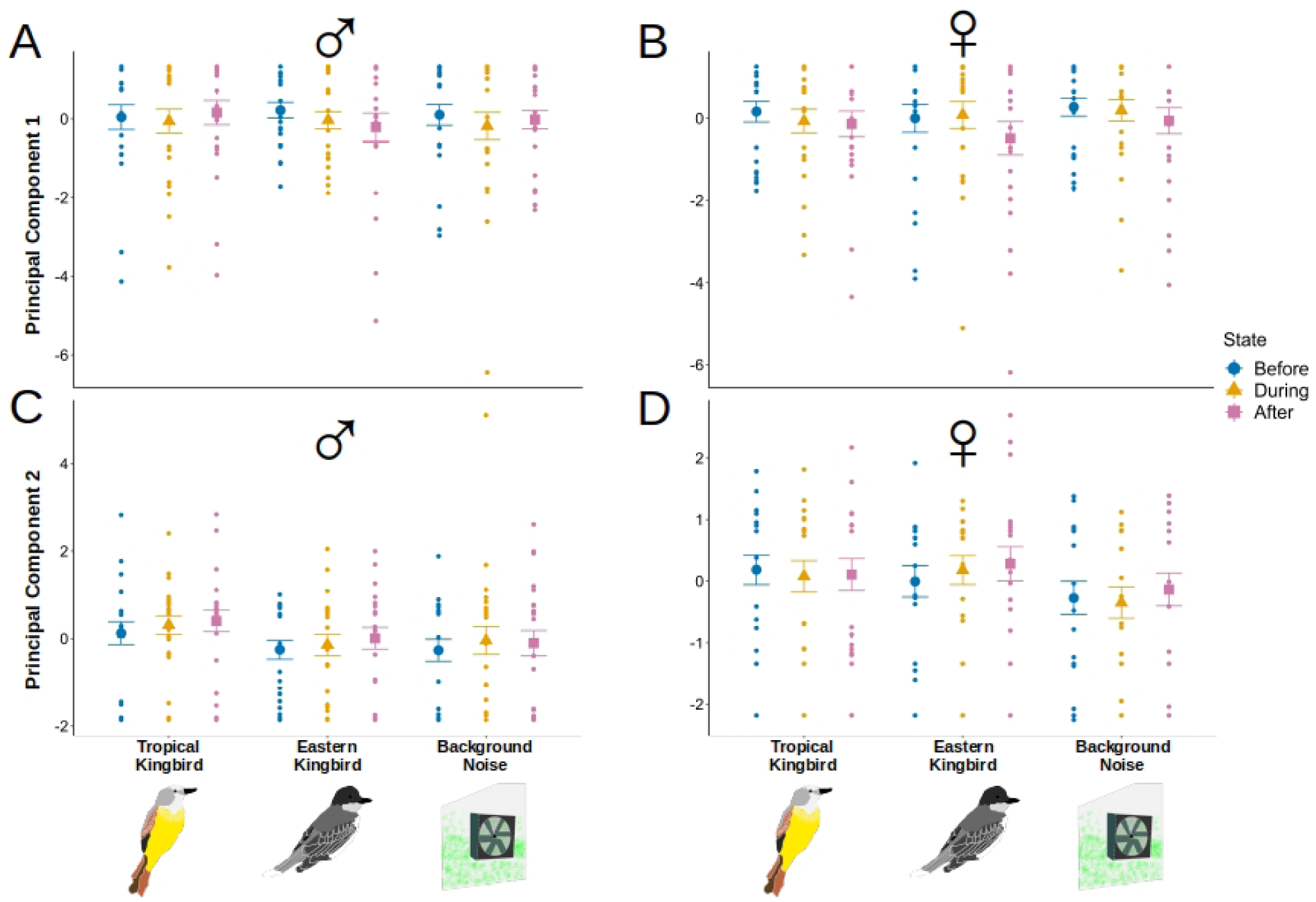
Mean ± SE of principal component variables for male and female *H. m. plessini* for a minute before, during and after calls. A) PC 1 in males for experiment 2; B) PC 1 in females for experiment 2; C) PC 2 in males for experiment 2; D) PC 2 in females for experiment 2. None of them are significantly different from each other.

### Residence status of kingbirds did not change long-term *H. m. plessini* behaviours

When combining the behavioural data for the 14 minutes before and after each call in a PCA for each sex, there was no effect of any of the kingbird calls or greenhouse background noise control on male PC1 (Supplementary Figure 6A; Supplementary Table 24, 25; see Supplementary Table 22 for PCA loadings), male PC2 (Supplementary Figure 6C; Supplementary Table 24, 25), male PC3 (Supplementary Table 24, 25); female PC1 (Supplementary Figure 6B; Supplementary Table 24, 25; see Supplementary Table 23 for PCA loadings), female PC2 (Supplementary Figure 6D, Supplementary Table 24, 25), or female PC3 (Supplementary Table 24, 25).

### No effect of predatory kingbird calls on *H. m. plessini* intersexual behaviours

Male *courtship, sitting near each other,* female *abdomen lifting, copulation* behaviours did not have short-term or long-term changes in response to any bird calls (Supplementary Table 20, 24).

## Discussion

*Heliconius melpomene plessini* butterflies changed their behaviour in response to predatory rufous-tailed jacamar calls but did not change their behaviour in response to predatory Eastern kingbird or tropical kingbird calls. We found a sex-specific difference in behaviour, where males, but not females, increased their fluttering and walking behaviours during the playback of the rufous-tailed jacamar calls. The observed behavioural changes in response to rufous-tailed jacamar calls are short-term and do not persist over an extended duration of time.

A major finding of this study is that toxic, unpalatable, and aposematic *Heliconius melpomene plessini* butterfly changed their behaviour in response to the predatory rufous-tailed jacamar calls. Contrary to our expectations, *H. m. plessini* butterflies did not change their behaviour in response to either the Eastern kingbird or tropical kingbird calls. Two non-mutually exclusive hypotheses can be postulated to explain these results: 1) There may be reduced predation pressure from both the Eastern and tropical kingbirds compared to rufous-tailed jacamar, which has led to an evolved behavioural response to the rufous-tailed jacamar but not to two kingbird species, and/or 2) *H. m. plessini* may be incapable of hearing the Eastern and tropical kingbird calls. Since Eastern kingbirds are migratory and tropical kingbirds are year-round residents, we had hypothesized that, if *H.m. plessini* could hear the Eastern and tropical kingbird calls, they might respond to the resident tropical kingbird due to their year-round presence, but not the migratory Eastern kingbird. Tropical kingbird calls are similar in frequency to the calls of Eastern kingbird (Supplementary Figure 2). However, we found that *H. m. plessini* butterflies did not change their behaviour in response to either of the kingbird calls, suggesting that between kingbird species variation in predation pressure was not sufficient to induce *H. m. plesseni* variation in response to Eastern and tropical kingbird calls.

*H. m. plessini* may be under reduced predation pressure from kingbirds relative to rufous-tailed jacamars. While the rufous-tailed jacamar is a year-round resident of *H. m. plesseni’s* habitat, the Eastern kingbird is migratory and is not present during half of the year in South America, where *H. m. plessini* is found. Eastern kingbird is also frugivorous during their migration over Central and South America (Blake & Loiselle, 1992; Morton, 1971). While this does not explain the lack of response to the tropical kingbird, an additional possibility is that *Heliconius melpomene* may be differentially palatable for rufous tailed jacamars and tropical kingbirds. Future studies should explore whether there is variability in toxicity across different subspecies of *H. melpomene,* or variability in predator sensitivity to *Heliconius* toxicity. Although there is no current support for this hypothesis in *Heliconius,* the aposematic striped skunks (*Mephitis mephitis*) perform anti-predatory behaviour in response to the calls of the great horned owl (*Bubo virginianus*) from which they are not chemically defended, but not in response to the calls of the coyote (*Canis latrans*), from which they are chemically defended (Fisher & Stankowich, 2018). Moreover, we found that *H. m. plessini* did not change their behaviour in response to the frugivorous control toco toucan bird call, despite the toucan calls being in the range of *Heliconius* hearing, which may indicate that *Heliconius* butterflies are capable of differentiating between predatory and non-predatory bird calls.

An alternative hypothesis is that *H. m. plessini* butterflies may not be capable of detecting kingbird calls but are able to detect the rufous tailed jacamar calls. Rufous tailed jacamar calls have a peak frequency below 4 kHz (Mikhail et al. 2018; Supplementary Figure 2), whereas both the Eastern and tropical kingbirds have a peak call frequency above 4 kHz (Supplementary Figure 2). Previous electrophysiological tests of the auditory organ in *H. erato* found that *H. erato* butterflies have the best hearing capabilities below 4 kHz at 70-90 dB power (Swihart, 1967). Any calls with frequencies above 4 kHz will require a higher decibel power to hear, which may be the case with the kingbird calls, as their peak call frequency is between 5-8 kHz. Similar trends have been observed in the blue morpho (*Morpho peleides*), and common wood nymph (*Cercyonis pegala*) butterflies, where a higher decibel power is required for higher frequency calls to elicit a response, and that these butterflies are tuned to hear sounds below 5 kHz (Fournier et al., 2013; Mikhail et al., 2018; Sun et al., 2018). Future studies in *Heliconius* can test this hypothesis by recording the butterfly responses to reduced frequency (below 4 kHz) kingbird calls and enhanced frequency (above 5kHz) rufous-tailed jacamar calls, and observe whether *H. m. plesseni* butterflies behaviourally respond to the altered kingbird and jacamar calls. We also found that *H. m. plessini* did not change their behaviour in response to the toco toucan calls and greenhouse background noise despite their calls being below 4 KHz, suggesting that *H. m. plessini* are able to distinguish between bird calls within their hearing range.

Our study is testing the hypothesis that *Heliconius* change their behaviours in response to predatory bird calls. Although an auditory organ has not yet been described in *Heliconius melpomene*, the auditory organ is described in a closely related butterfly *Heliconius erato* (Swihart, 1967). Here we do provide evidence that *H. m. plesssini* changed their behaviour after hearing their predatory rufous-tailed jacamar calls. Future work can explore the presence of a morphological hearing structure in *Heliconius melpomene plessini* and their electrophysiological range like that performed in other butterflies (Lane et al. 2008; Lucas et al. 2009; Mikhail et al. 2018), to enhance our understanding of the physiological mechanisms *H. m. plesseni* may be using to facilitate their response to the rufous-tailed jacamar.

We found that males, but not females, changed their behaviour in response to the rufous-tailed jacamar calls. This male-specific response to predators is similar to that found in other species, and may reflect sexual dimorphic predation pressures. Previous studies in wolf spiders (*Pardosa milvina*) have found that males, but not females, used a predatory chemical cue experience to decrease predation from a live predator (Sitvarin and Rypstra, 2012). Similarly, male yellow-billed marmots (*Marmota flaviventris*) decreased foraging followed by a playback of alarm calls (Lea and Blumstein, 2011). The sex-specific differences observed in the response of *H. m. plessini* might reflect differences in predation pressures between the sexes. Male *Heliconius* butterflies in the wild spend greater time flying in the middle of the forest canopy, and mostly near their larval/food plants in search of females or for foraging whereas female *Heliconius* spend time fluttering near the understory in search of host plants for egg-laying (Mallet and Gilbert, 1995). Jacamars and kingbirds are “aerial hawking” predators that catch insects in flight (Fitzpatrick, 1980), and flying male butterflies might be at greater risk of predation. This may be the reason for increased fluttering and walking during the jacamar calls. Moreover, Swihart observed fast wing flutters in *H. erato* when he exposed them to loudspeaker generated sound (Swihart, 1967), indicating that butterflies may have an innate wing fluttering response to sound cues. Similar results have been found in *Erebia* butterflies, where they flutter in response to sound (Ribaric and Gogala 1996) and in the peacock butterflies (*Inachis io*) where they walk and flutter to avoid rodent predation during winter hibernation (Olofsson et al., 2011). In *Heliconius*, fluttering may advertise aposematic colouration and could reinforce the birds’ learned behaviour to avoid brightly coloured butterflies (Langham, 2006). Similar to the mimicry of aposematic colours among *Heliconius* species, there is also evidence of locomotor mimicry in the flight of unpalatable *Heliconius*, including flight measures associated with response to jacamars (Chai & Srygley, 1990; Srygley, 1994). Future studies of the responses in *H. melpomene,* their model *H. erato* and other species of the same aposematic mimicry rings could inform us if certain predators have influenced the evolution of mimetic behavioural responses.

Palatability experiments with jacamars have found that experienced birds sight-reject flying *Heliconius* butterflies (Pinheiro & Campos, 2019). Therefore, flying, fluttering and walking behaviours could be advantageous under different ecological contexts (for example bird predator community and experience) as an immediate response to predator’s presence, which may be another reason why we did not see the behavioural changes over a long-term (14 minutes) period. Future studies could look at the advantages of these behaviours under different ecological contexts such as microhabitats (Dell’Aglio et al., 2022), as well as test the behavioural responses of the butterflies using other predatory birds.

## Conclusions

We found that unpalatable and brightly coloured *Heliconius melpomene plessini* butterflies respond and change their behaviour during the playback of the rufous-tailed jacamar call. This change in behaviour is sex-specific, where males, but not females, increase their walking and fluttering behaviour over a short time-frame. Males reverted back to their original behaviour after the call ended. *H. m. plessini* did not change their behaviour in response to the two kingbird and the toco toucan calls. Our study opens avenues for future research in the field of butterfly auditory anti-predatory behaviour response, its mechanistic underpinnings and ecological and evolutionary consequences, especially in the context of mimicry.

## Acknowledgments

We thank David A. Ernst, Deonna N. Robertson, Grace Hirzel, Matthew Murphy, Yi Ting Ter, Kiana Kasmaii, and Keity Farfán Pira for their contribution towards *Heliconius* butterfly husbandry and reviewing the manuscript. We thank Brian Counterman for providing valuable inputs and reviewing this manuscript. We also thank Pooja Panwar for fruitful discussions during the conceptualization of this project.

## Data Availability

Analyses reported in this article can be reproduced using the data provided by the authors in Dryad (link: XXXX)

## CRediT author contributions

SP- Conceptualization, Methodology, Investigation, Data curation, Formal analysis, Funding acquisition, Writing- original draft, Writing- review and editing; MD- Data curation, Software, Writing- review and editing; ELW- Conceptualization, Methodology, Supervision, Resources, Funding acquisition, Writing- original draft, Writing- review and editing.

## Supplementary Material 1

**Supplementary Figure 1:**
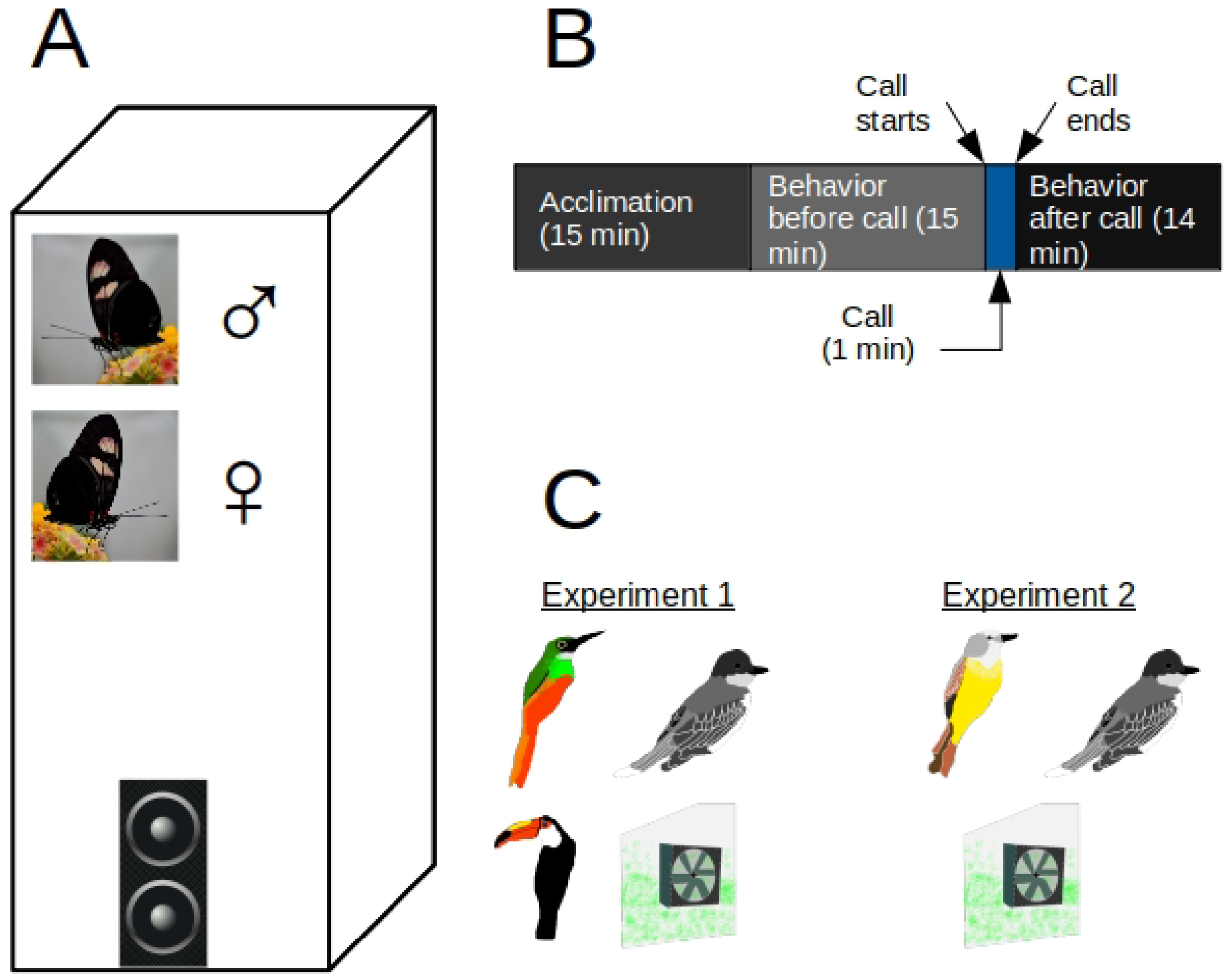
Experimental design. A) 3-15-day male and female *H. m. plessini* butterflies were subjected in an experimental cage with a blue tooth speaker and a *Lantana spp.* plant during each experimental assay. B) The timeline of each assay conducted where the butterflies were acclimated for 15 minutes and their behaviours recorded for the next 30 minutes. During the 16 ^th^ minute, a call was randomly played for a minute. C) The calls used in the two experiments in this study. Clockwise from top left in experiment 1: rufous-tailed jacamar, Eastern kingbird, greenhouse background noise, and toco toucan. Clockwise from top left in experiment 2: tropical kingbird, Eastern kingbird, and greenhouse background noise.

**Supplementary Figure 2:**
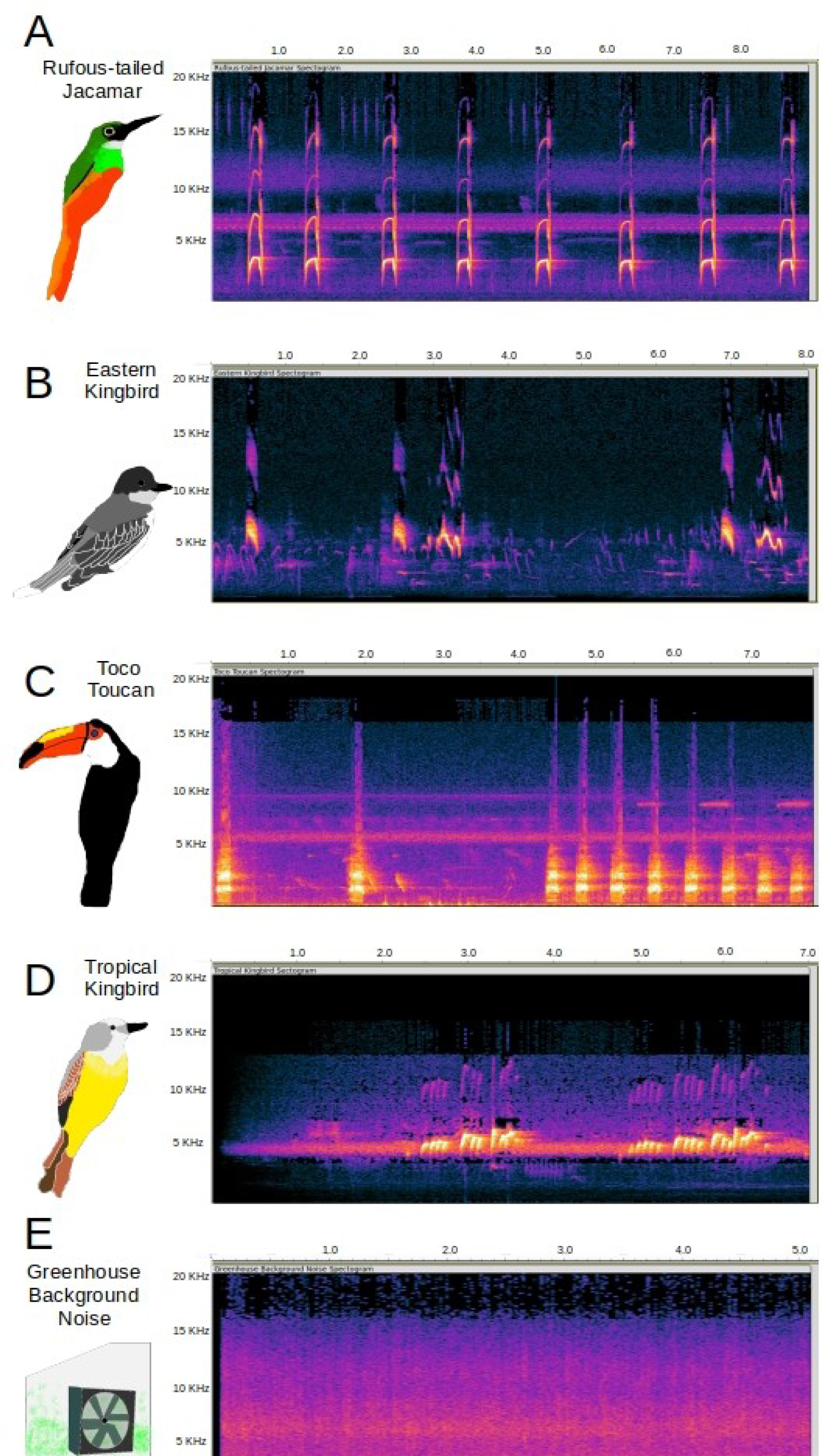
Spectograms of the calls used during this study A) rufous-tailed jacamar; B) Eastern kingbird; C) toco toucan; D) tropical kingbird; E) greenhouse background noise.

**Supplementary Figure 3:**
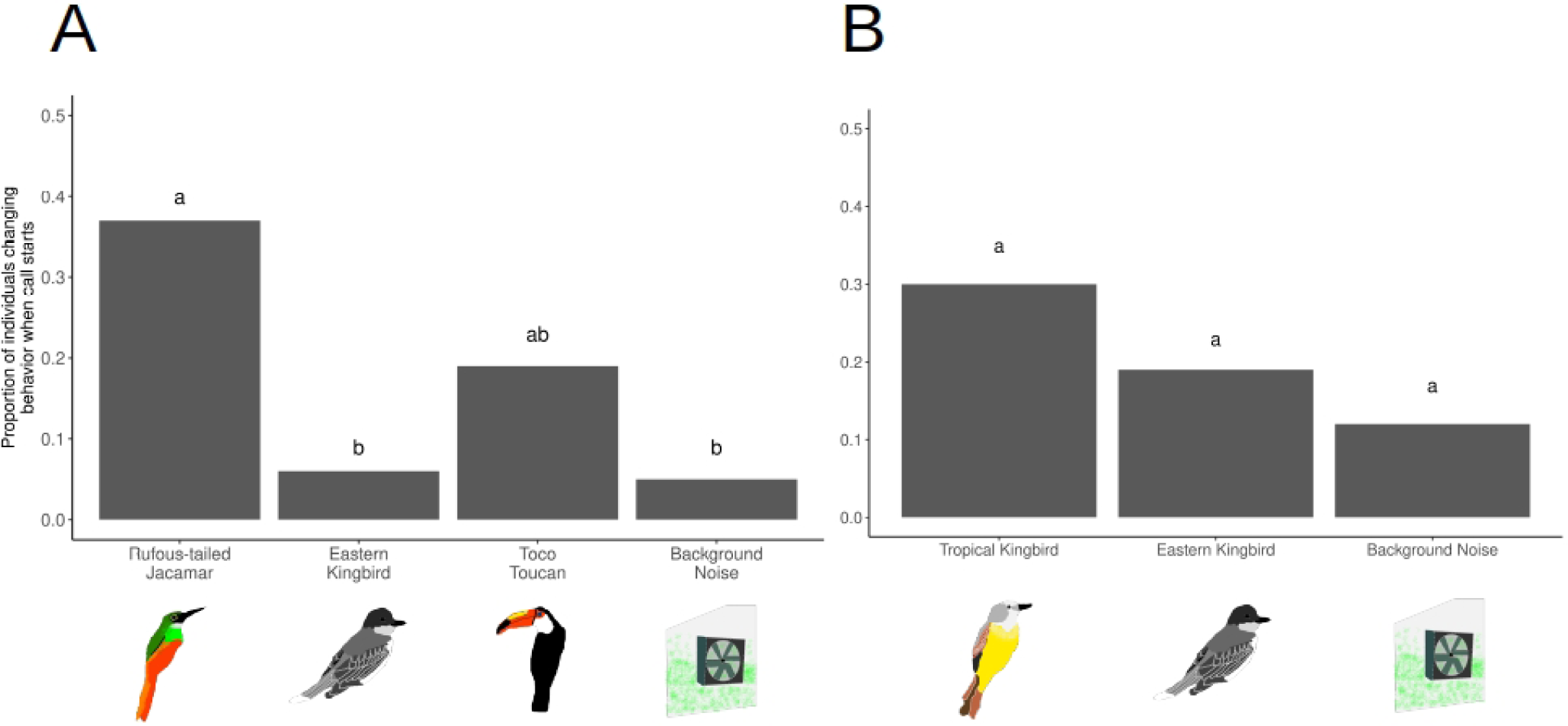
Proportion of *H. m. plessini* individuals changing behaviour in response to the start of the calls (between before start and after start of calls) for A) experiment 1; B) experiment 2; Different letters on each bars indicate statistical significance at p<0.05.

**Supplementary Figure 4:**
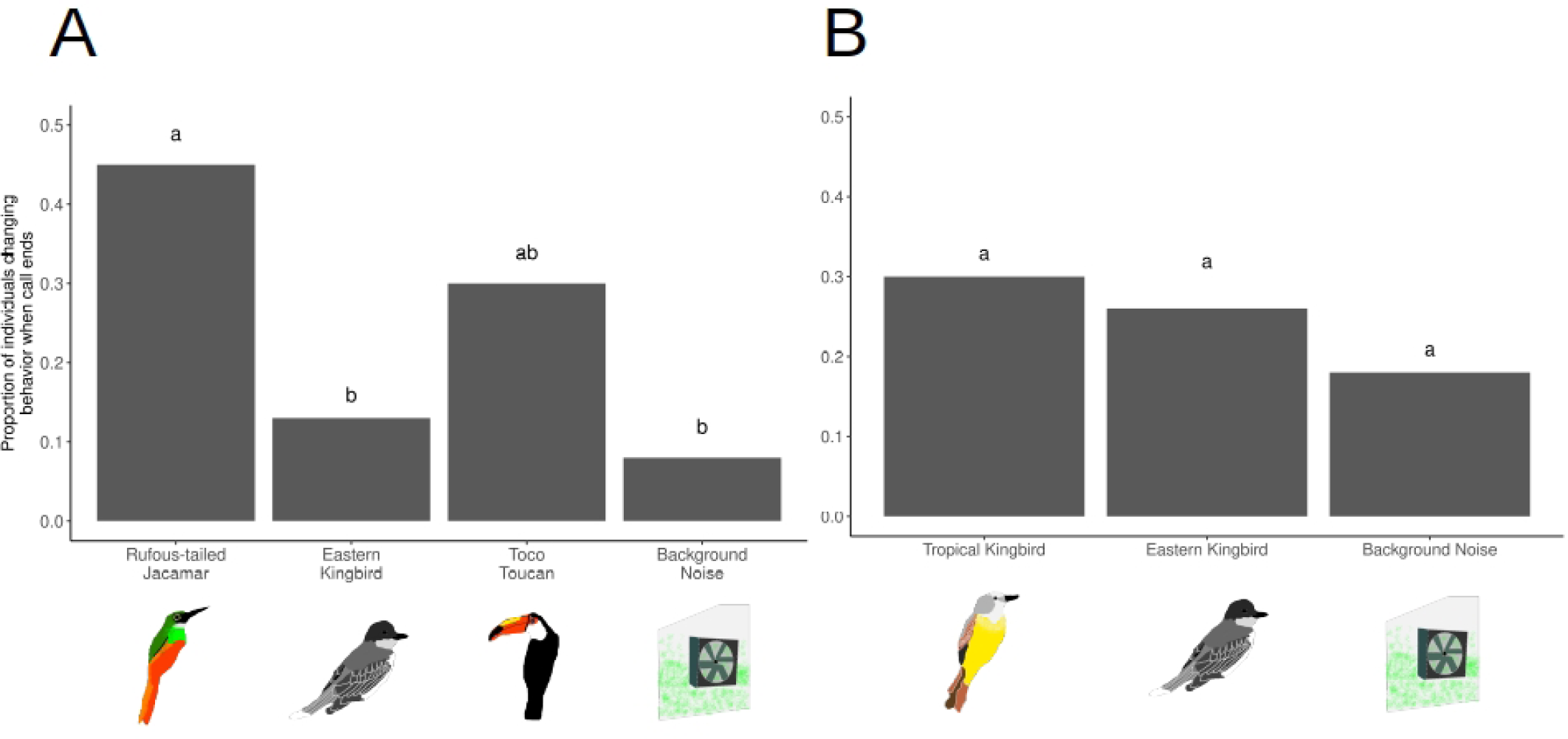
Proportion of *H. m. plessini* individuals changing behaviour in response to the end of the calls (between before end and after end of calls) for A) experiment 1; B) experiment 2; Different letters on each bars indicate statistical significance at p<0.05.

**Supplementary Figure 5:**
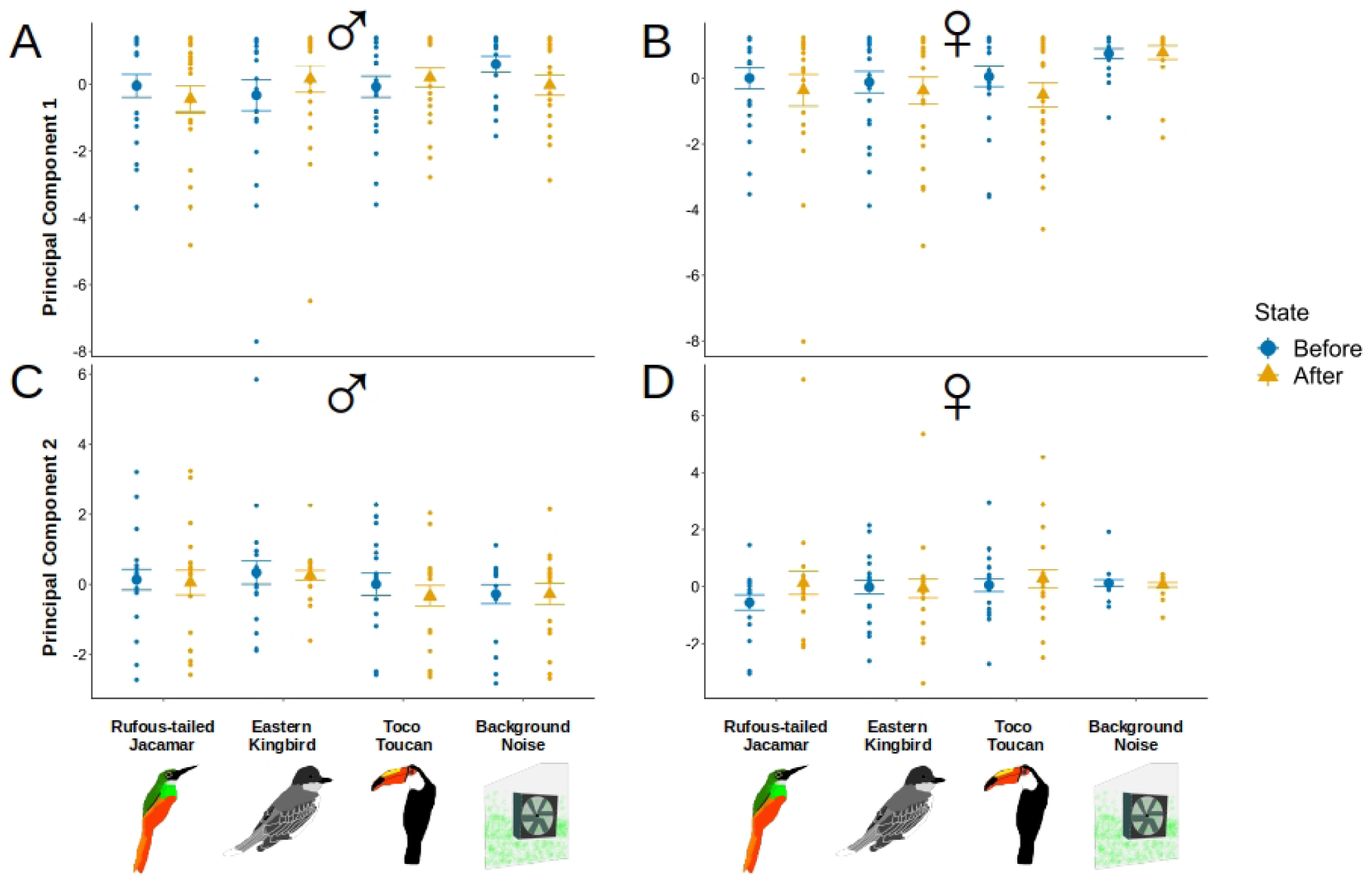
Mean ± SE of principal component variables for male and female *H. m. plessini* for 14 minutes before, and after calls. A) PC 1 in males for experiment 1; B) PC 1 in females for experiment 1; C) PC 2 in males for experiment 1; D) PC 2 in females for experiment 1. None of them are significantly different from each other.

**Supplementary Figure 6:**
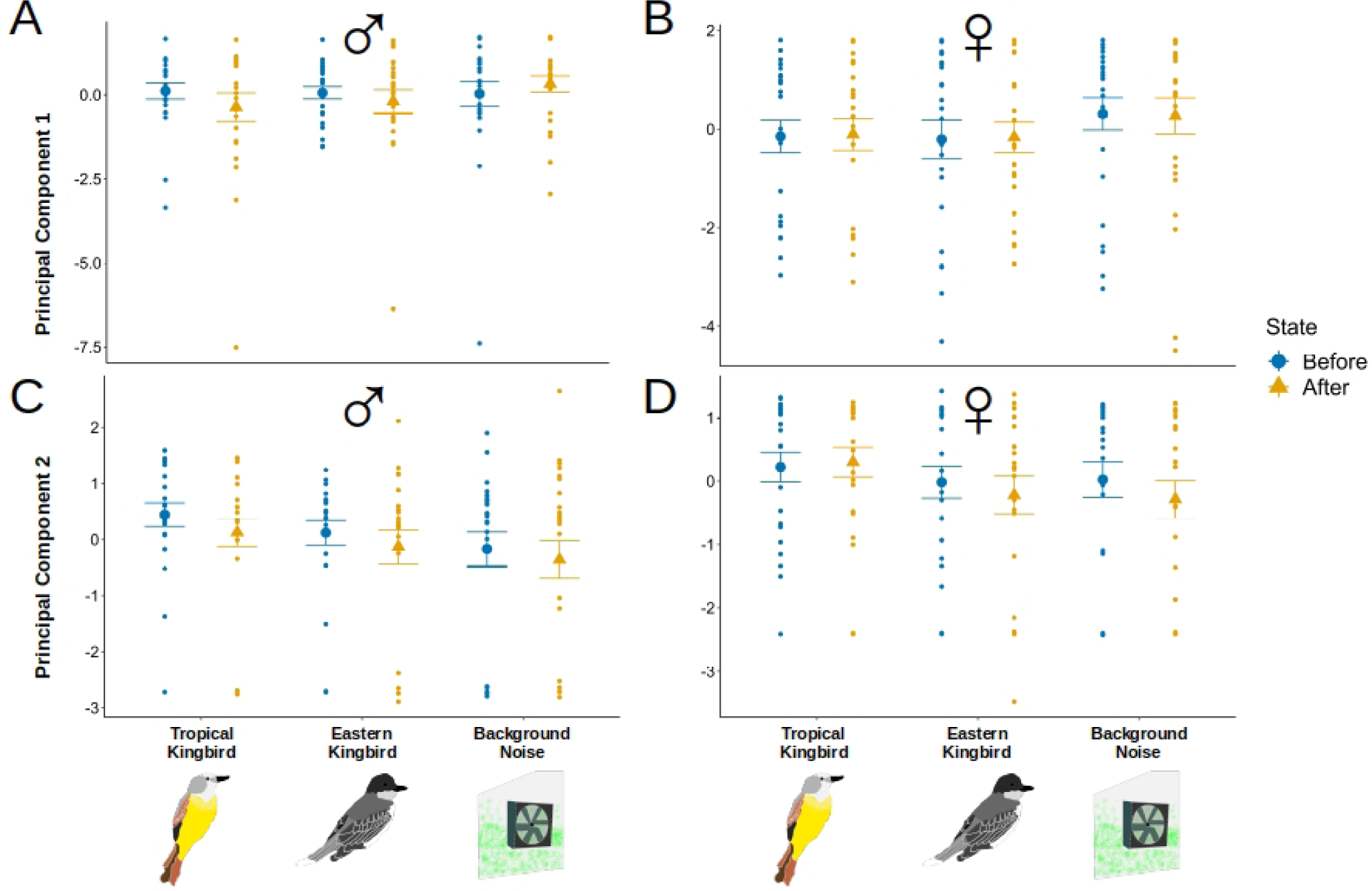
Mean ± SE of principal component variables for male and female *H. m. plessini* for 14 minutes before, and after calls. A) PC 1 in males for experiment 2; B) PC 1 in females for experiment 2; C) PC 2 in males for experiment 2; D) PC 2 in females for experiment 2. None of them are significantly different from each other.

**Supplementary Table 1:**
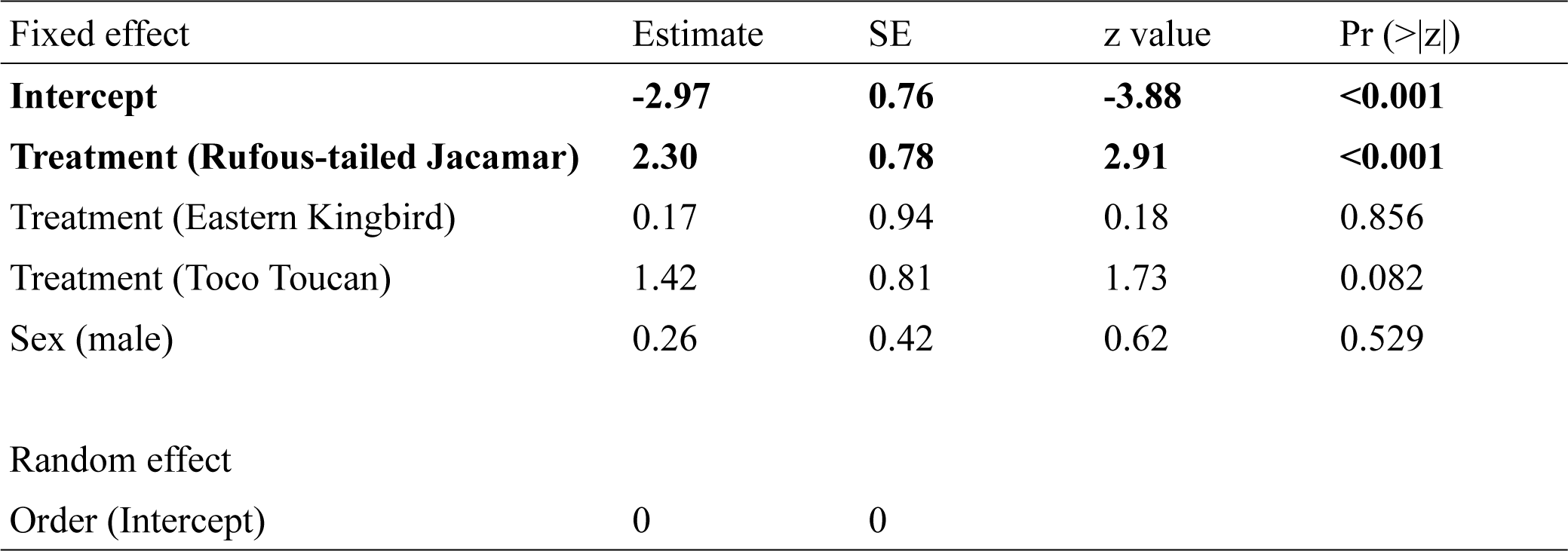
GLMM results on the effect of treatment (calls) and sex on proportion of butterflies changing their behaviour at the start of calls in experiment 1. p<0.05 are bolded.

**Supplementary Table 2:**
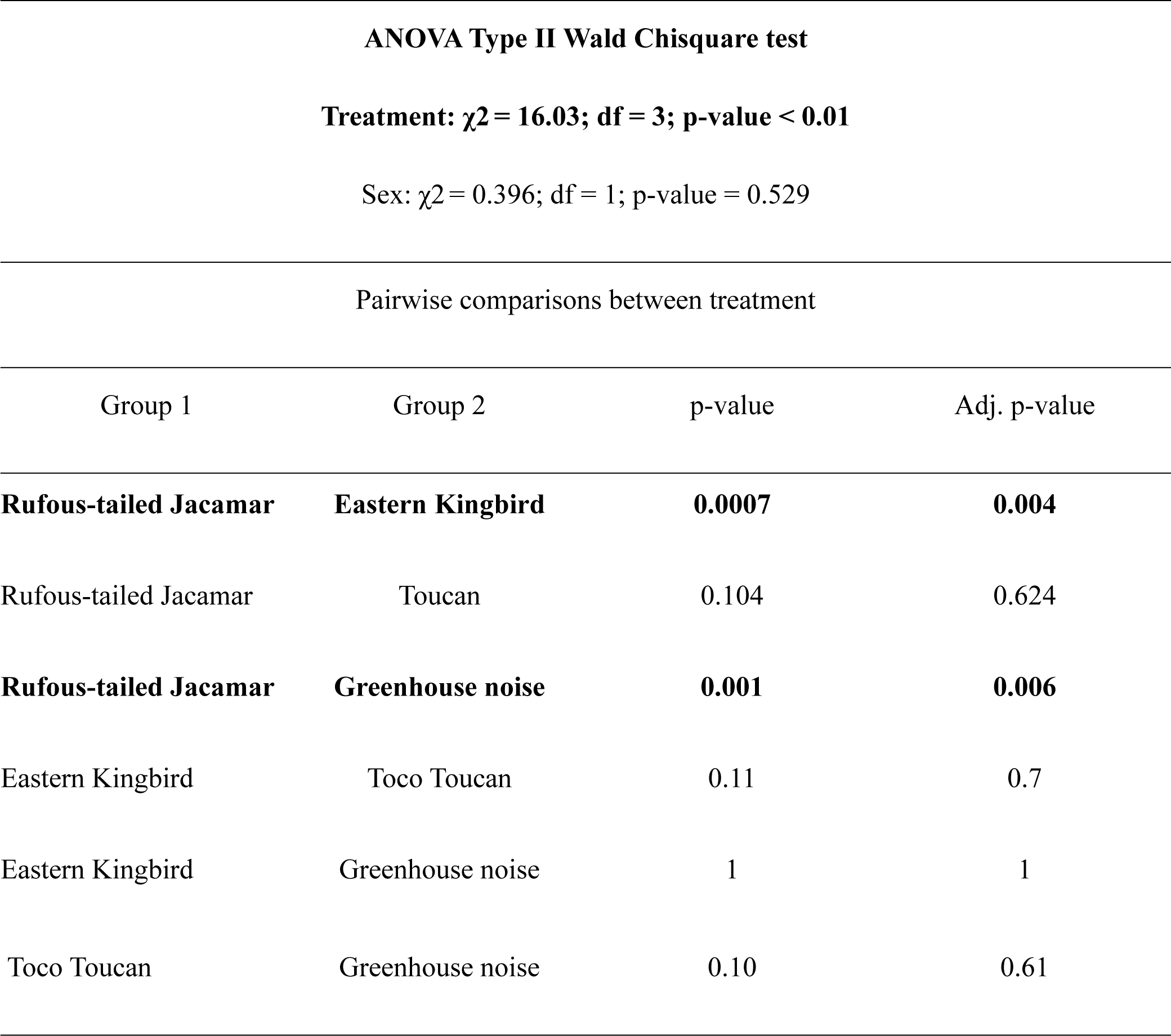
Pairwise differences in the proportion of individuals changing their behavioural state in at the start of calls in experiment 1. p<0.05 are bolded.

**Supplementary Table 3:**
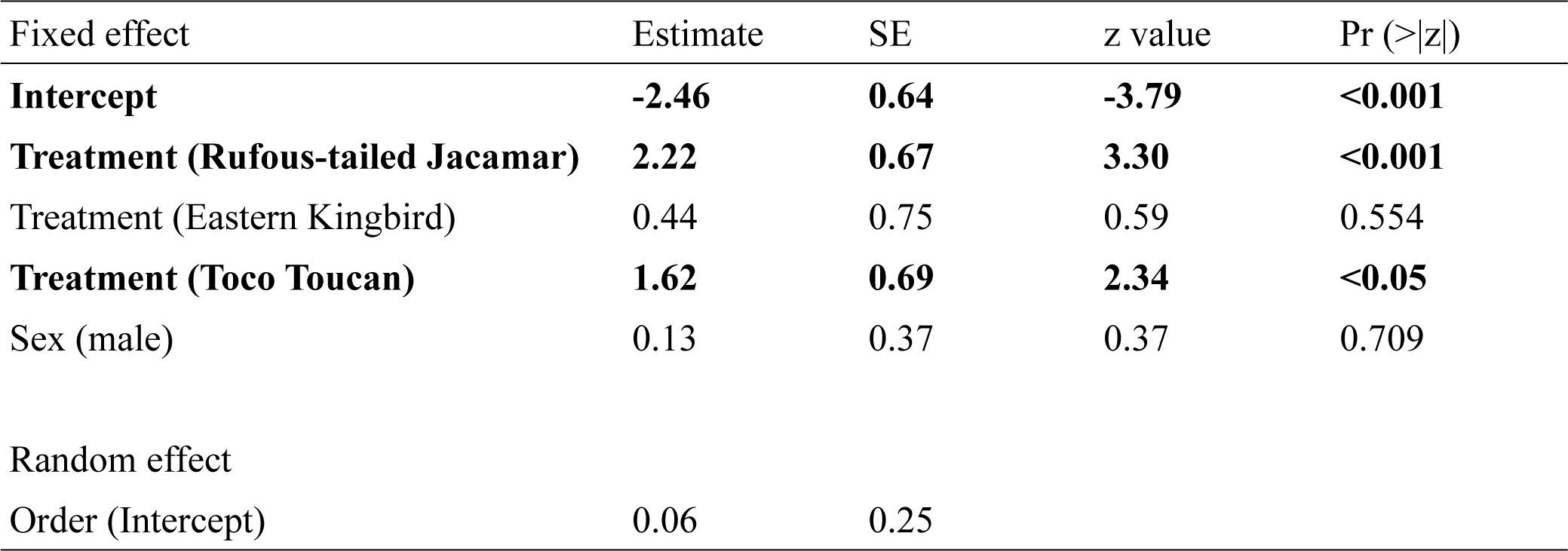
GLMM results on the effect of treatment (calls) and sex on proportion of butterflies changing their behaviour at the end of calls in experiment 1. p<0.05 are bolded.

**Supplementary Table 4:**
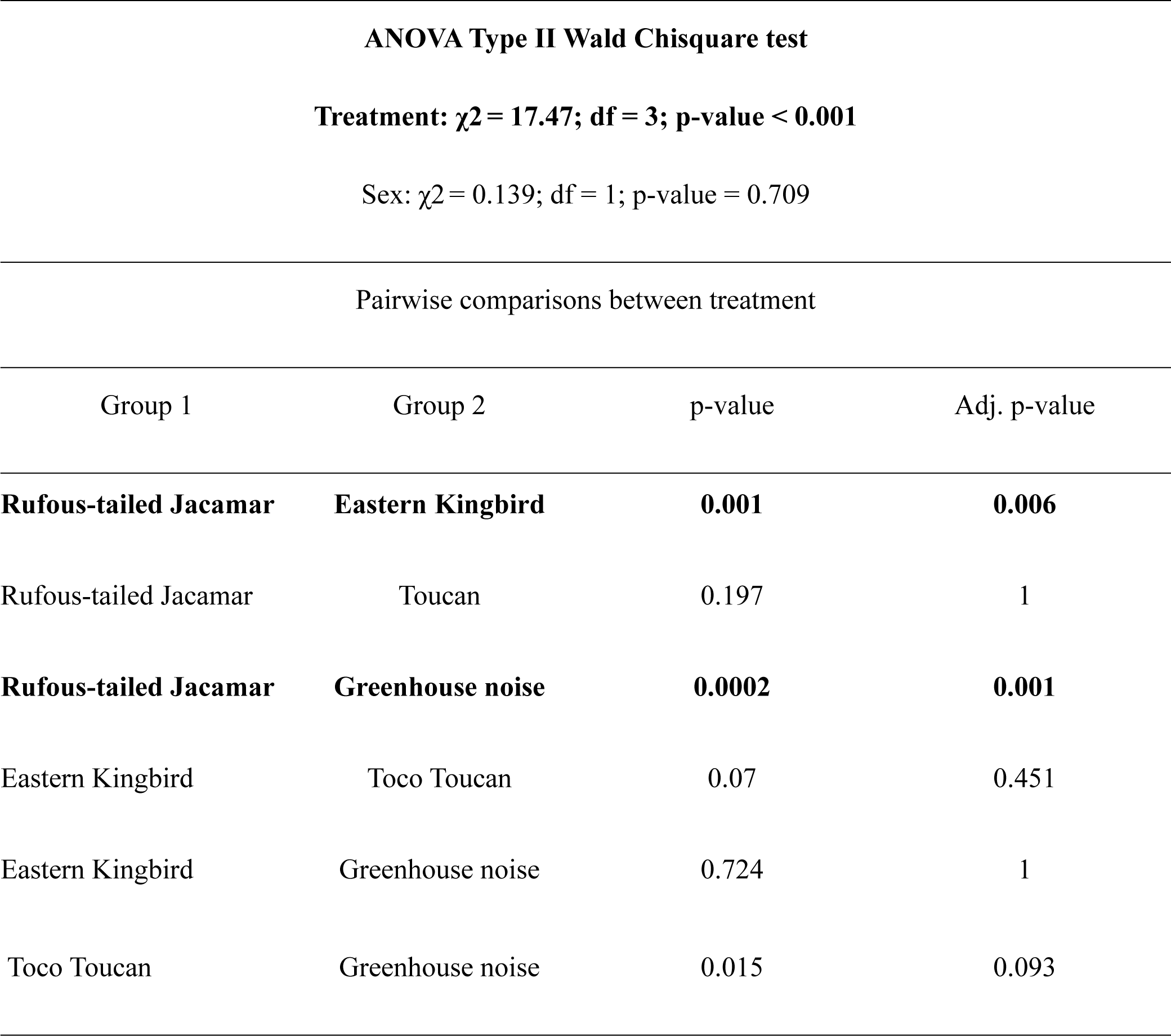
Pairwise differences in the proportion of individuals changing their behavioural state in at the end of calls in experiment 1. p<0.05 are bolded.

**Supplementary Table 5:**
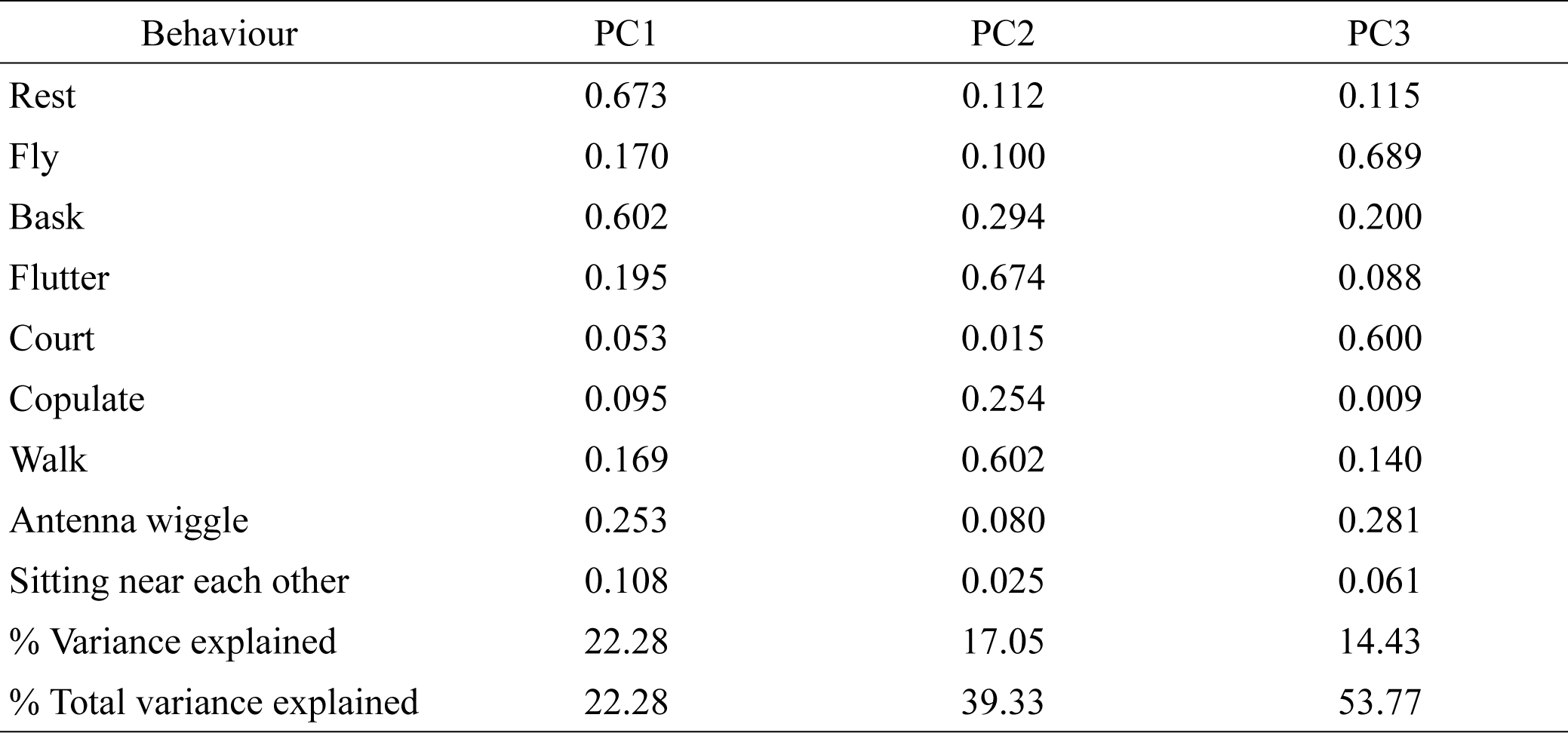
Loadings of each behaviour in Principal Component (PC) composite variables for males in a minute before, during, and after calls in experiment 1.

**Supplementary Table 6:**
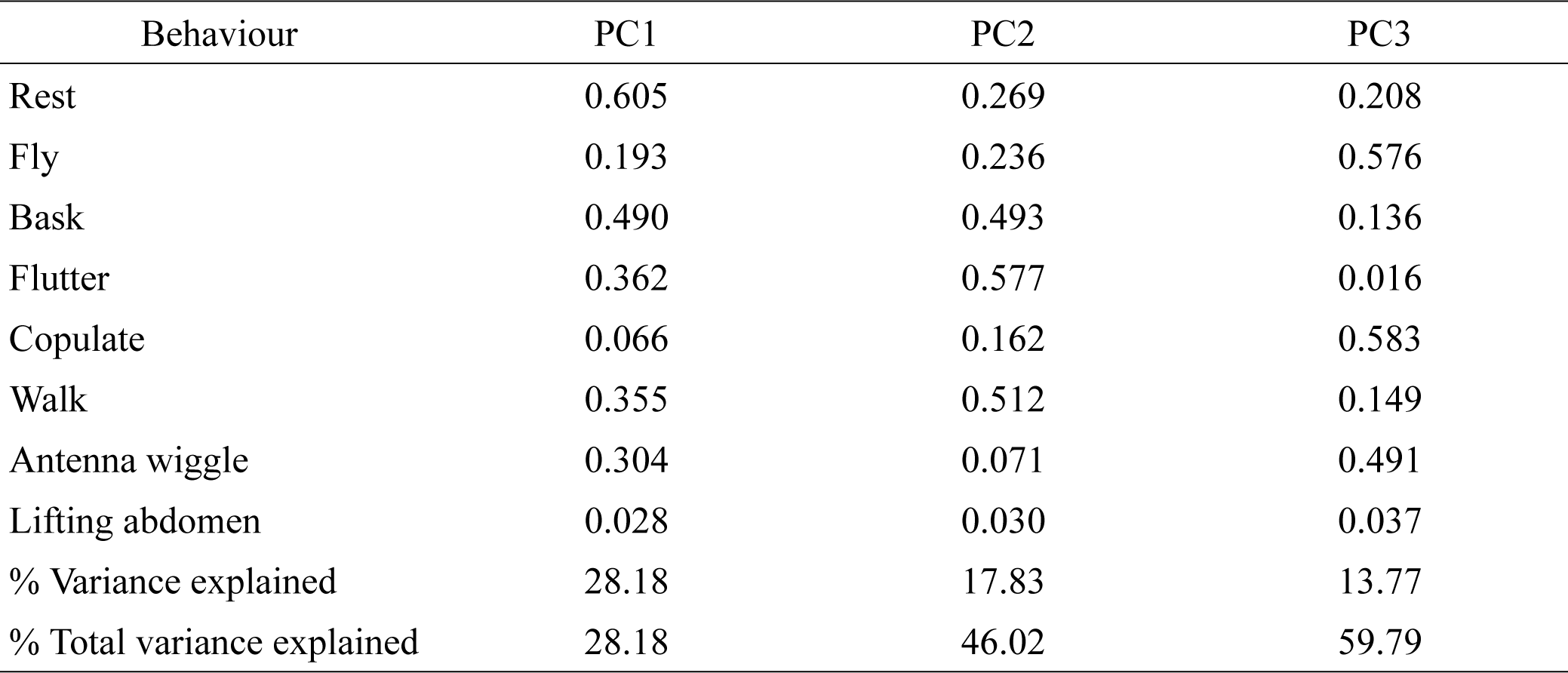
Loadings of each behaviour in Principal Component (PC) composite variables for females in a minute before, during and after calls in experiment 1.

**Supplementary Table 7:**
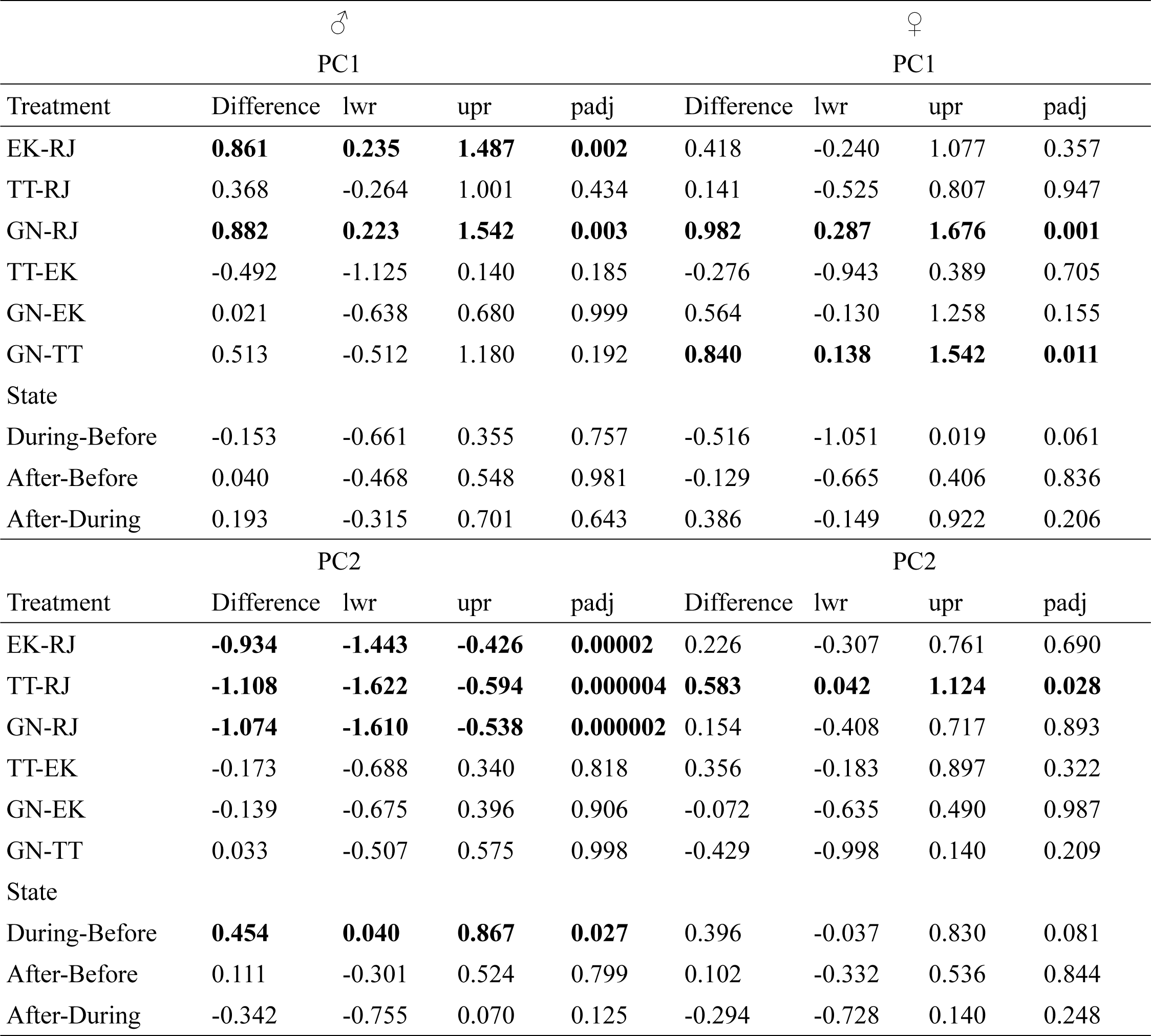
ANOVA Post-hoc test results with PC as dependent variables and the treatments (calls), state (before, during, and after) and their interaction as response variables for males and females in a minute before, during, and after calls in experiment 1. EK= Eastern kingbird; RJ=Rufous-tailed jacamar; TT=Toco toucan; GN=Greenhouse background noise; Difference=pairwise difference; lwr=lower range; upr=upper range; padj= adjusted p-value.

**Supplementary Table 8:**
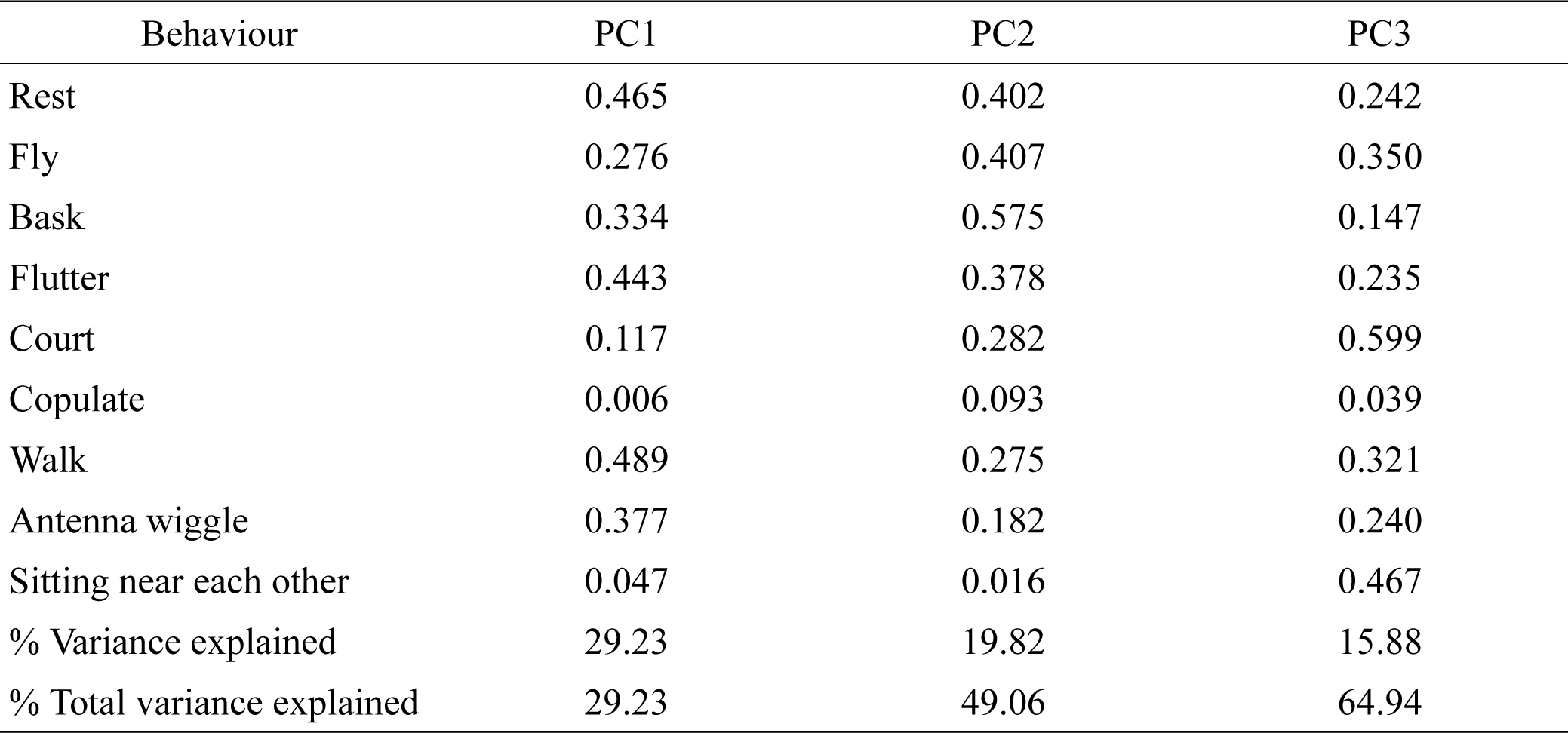
Loadings of each behaviour in Principal Component (PC) composite variables for males in 14 minutes before and after calls in experiment 1.

**Supplementary Table 9:**
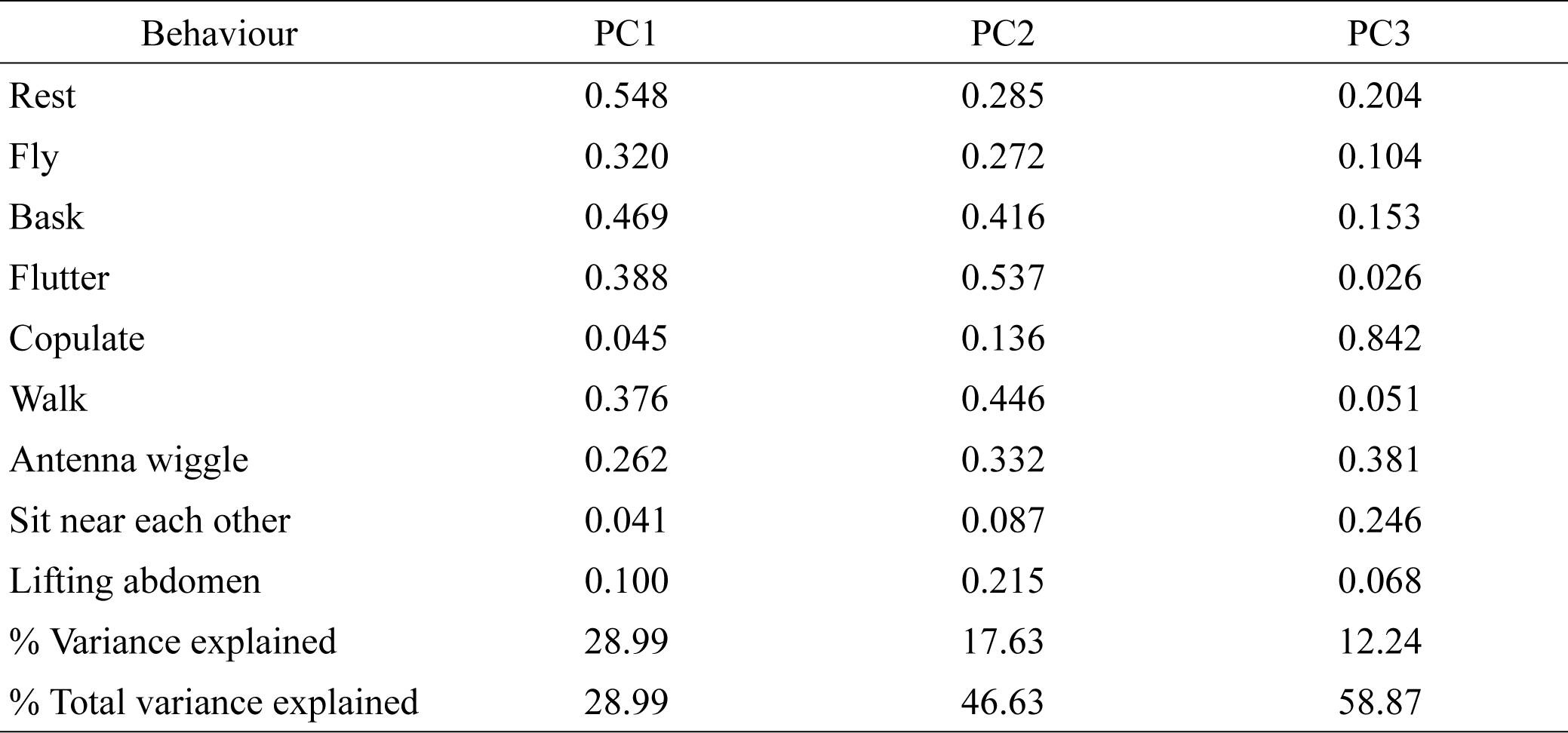
Loadings of each behaviour in Principal Component (PC) composite variables for females in 14 minutes before and after calls in experiment 1.

**Supplementary Table 10:**
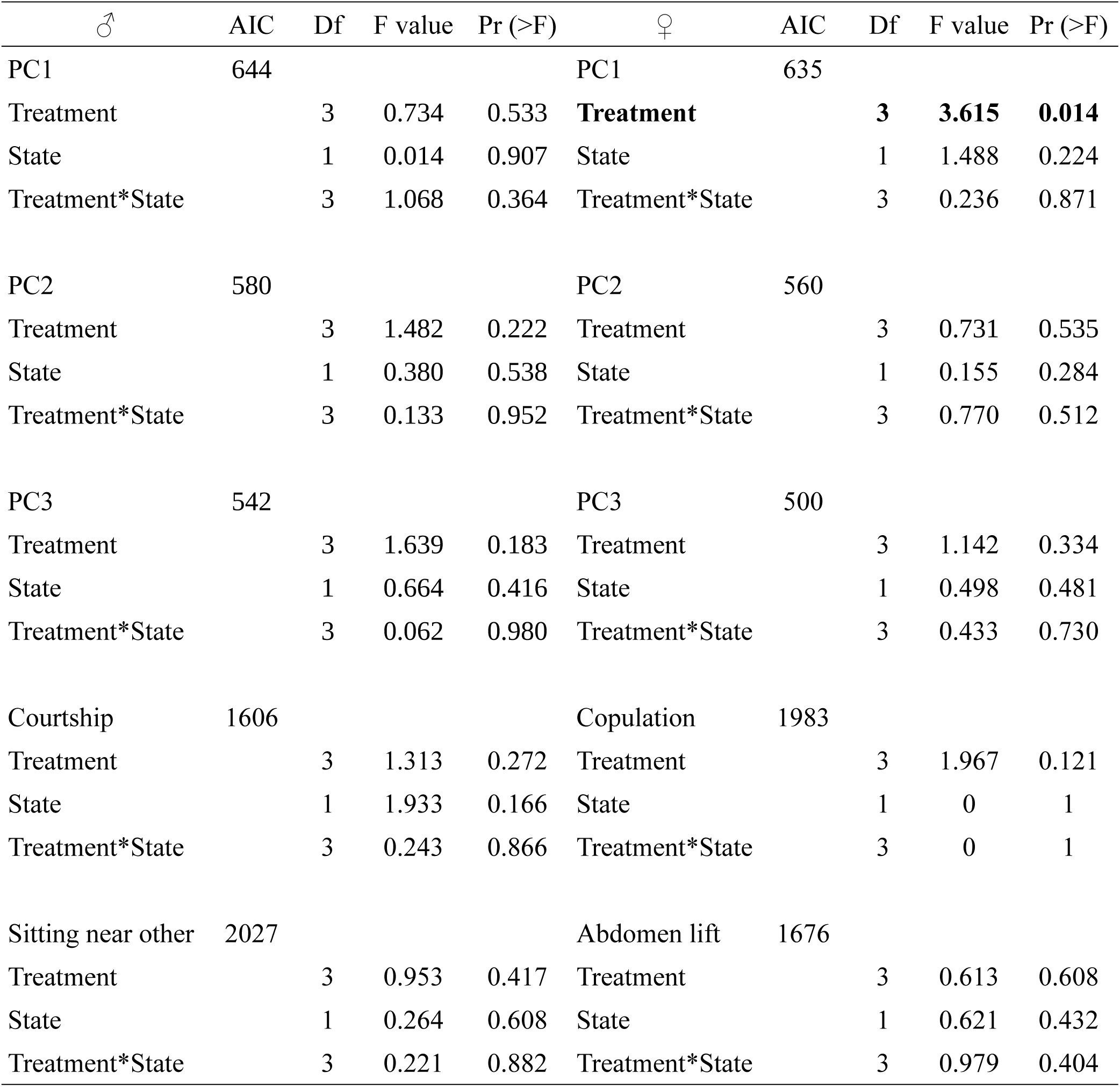
Effect of treatment (Rufous-tailed Jacamar, Eastern Kingbird, Toco Toucan and Greenhouse background noise calls), state (before call and after call) and their interaction on 14 minute behaviours before and after call, and male PC1, PC2, PC3, *courtship, copulation, sit near* and female PC1, PC2, PC3, *abdomen lifting* behaviours in experiment 1.

**Supplementary Table 11:**
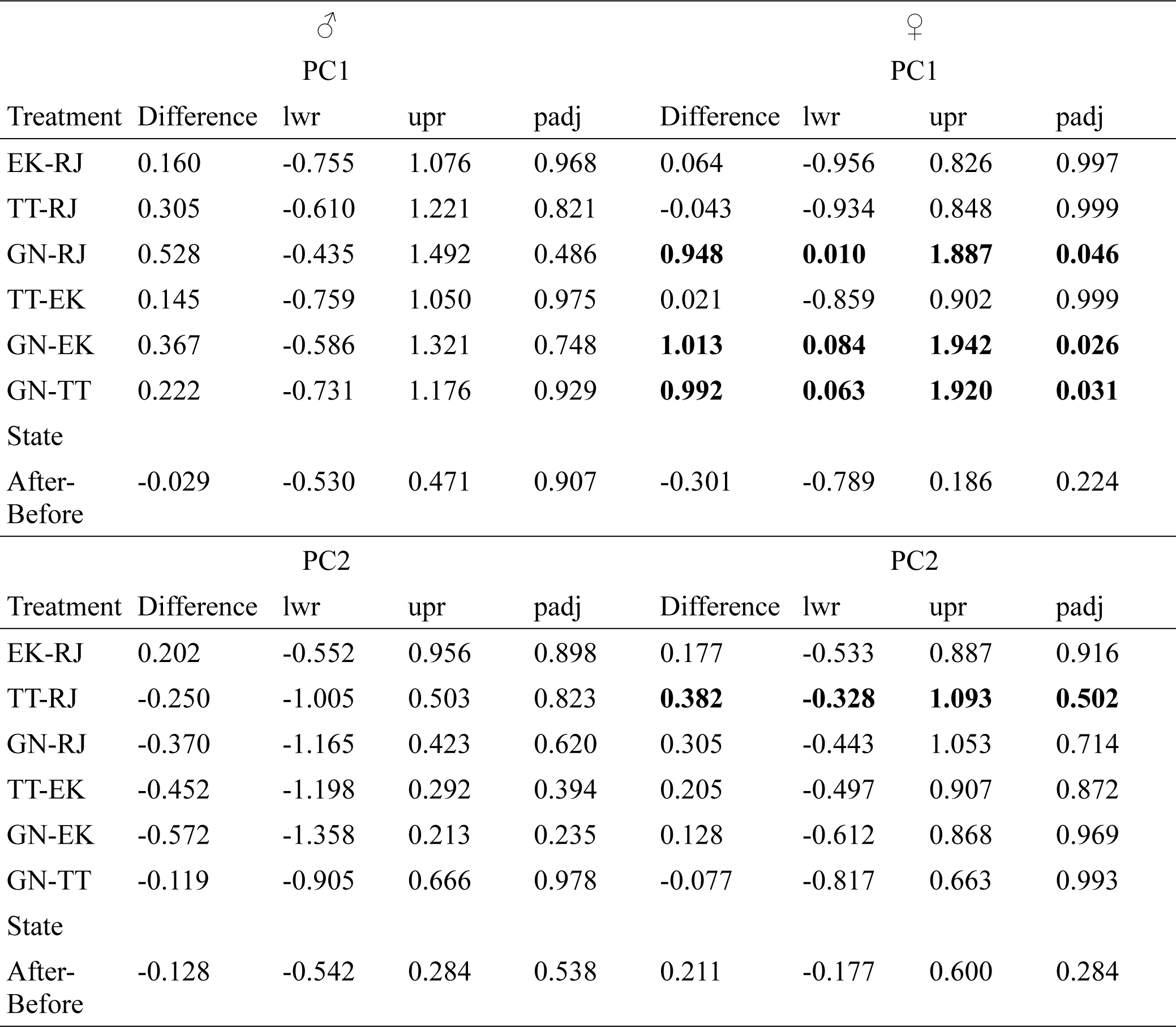
ANOVA post-hoc test results with PC as dependent variables and the treatments (calls), state (before and after) and their interaction as response variables for males and females in 14 minutes before and after calls in experiment 1. EK= Eastern kingbird; RJ=Rufous-tailed jacamar; TT=Toco toucan; GN=Greenhouse background noise; Difference= pairwise difference; lwr=lower range; upr=upper range; padj= adjusted p-value.

**Supplementary Table 12:**
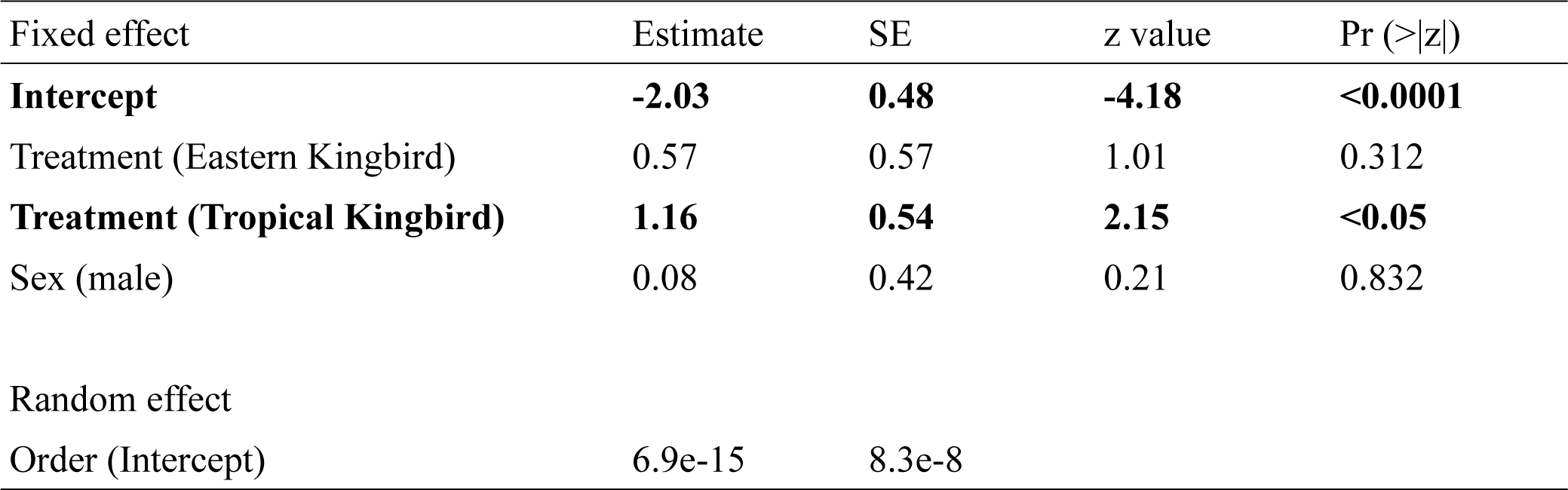
GLMM results on the effect of treatment (calls) and sex on proportion of butterflies changing their behaviour at the start of calls in experiment 2. p<0.05 are bolded.

**Supplementary Table 13:**
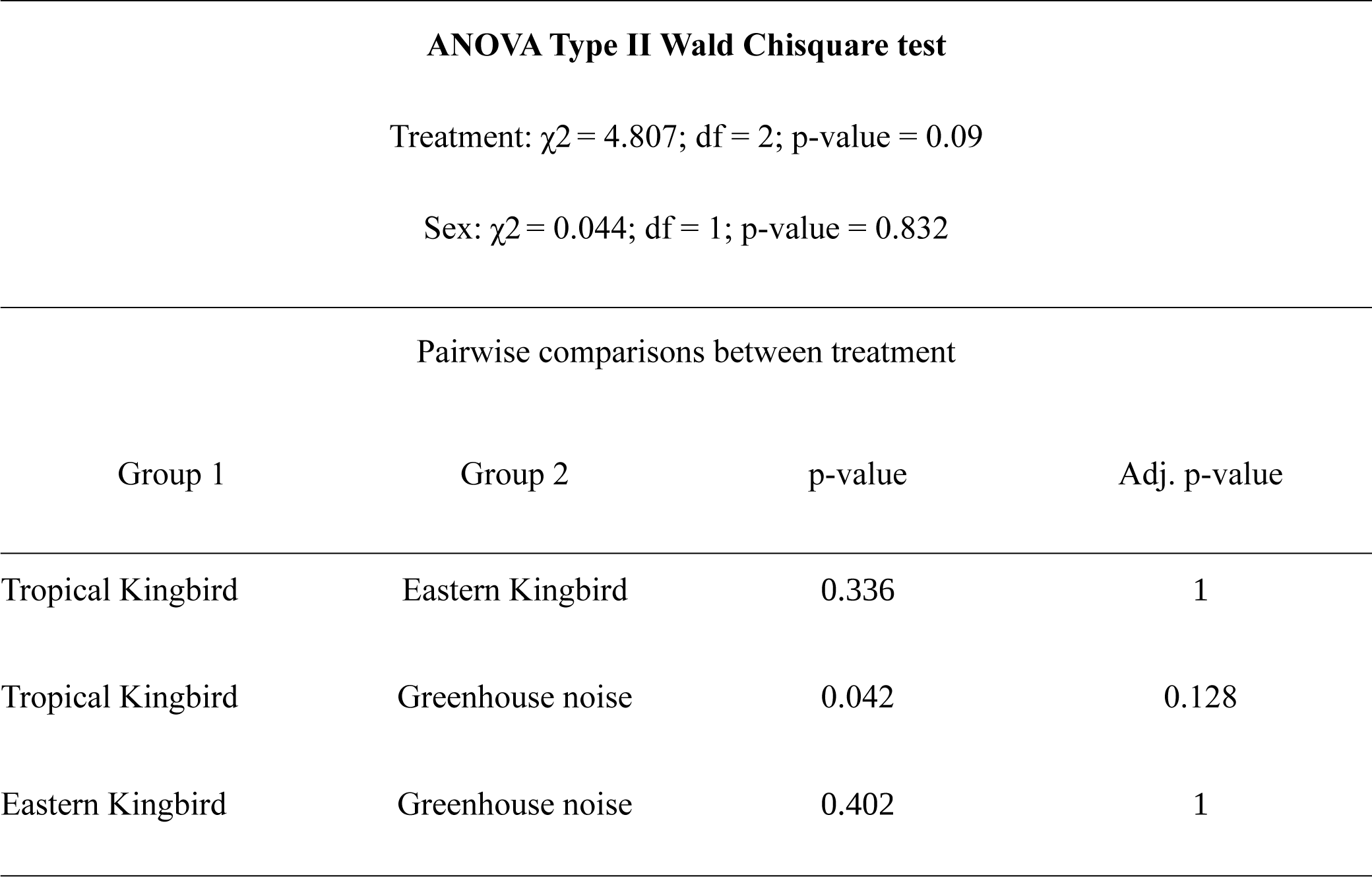
Pairwise differences in the proportion of butterflies changing their behavioural state in response to the start of calls in experiment 2.

**Supplementary Table 14:**
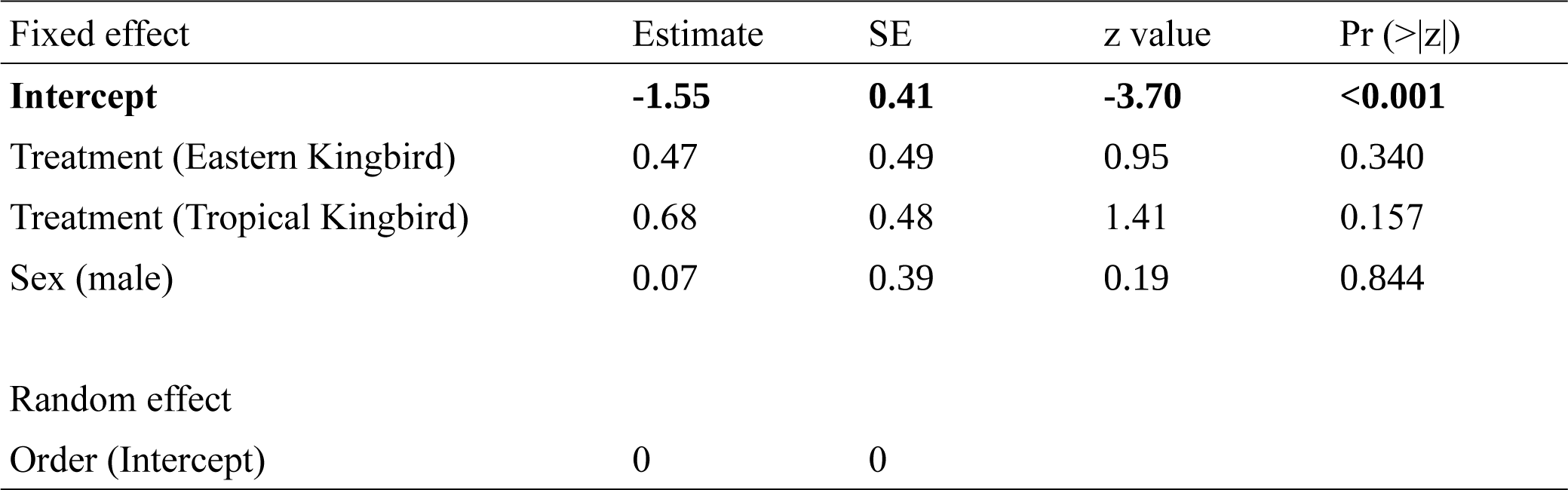
GLMM results of the effect of treatment (calls) and sex on proportion of butterflies changing their behaviour at the end of calls in experiment 2. p<0.05 are bolded.

**Supplementary table 15:**
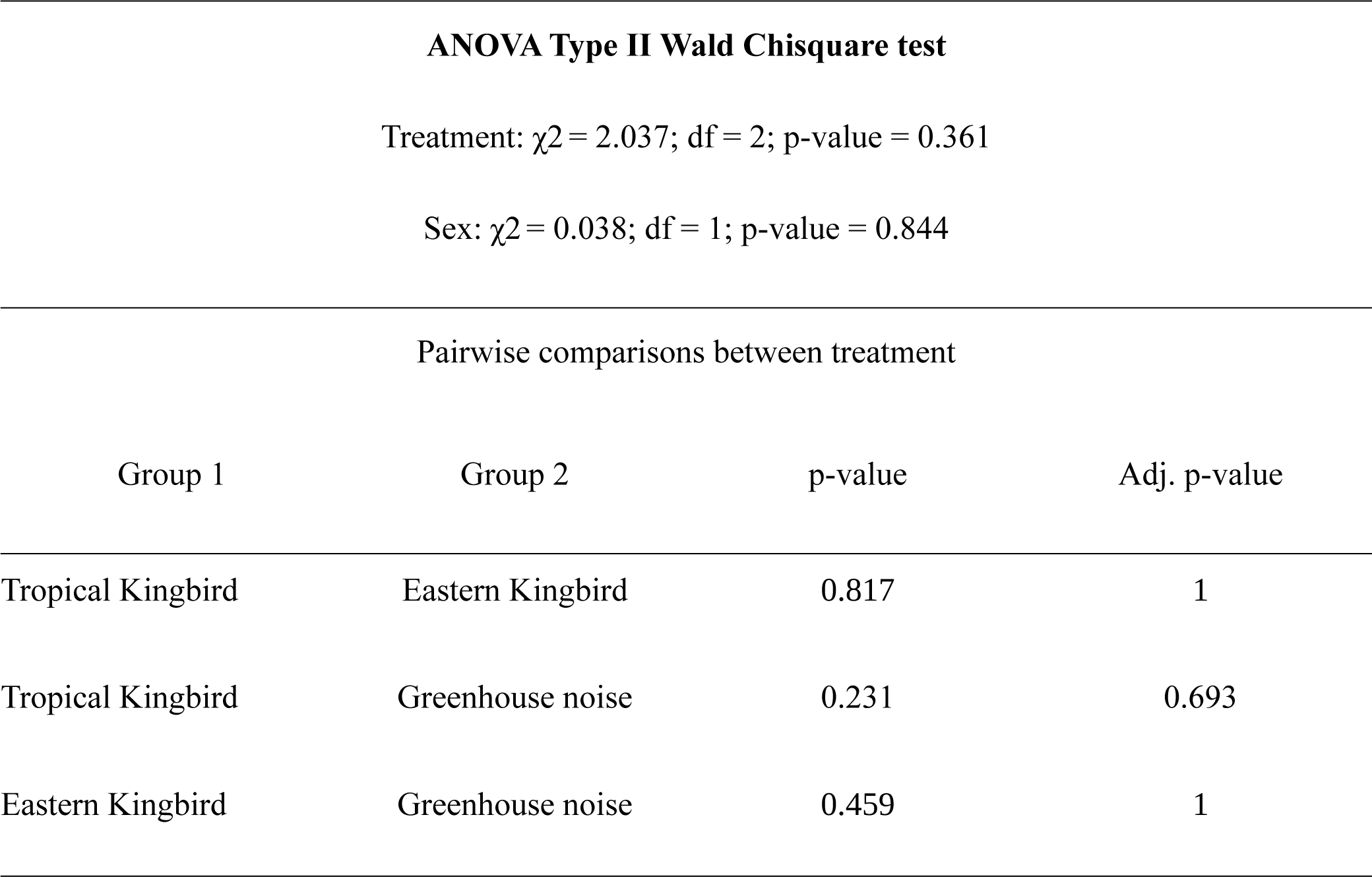
Pairwise differences in the proportion of males and females changing their behavioural state in response to the end of calls in experiment 2.

**Supplementary Table 16:**
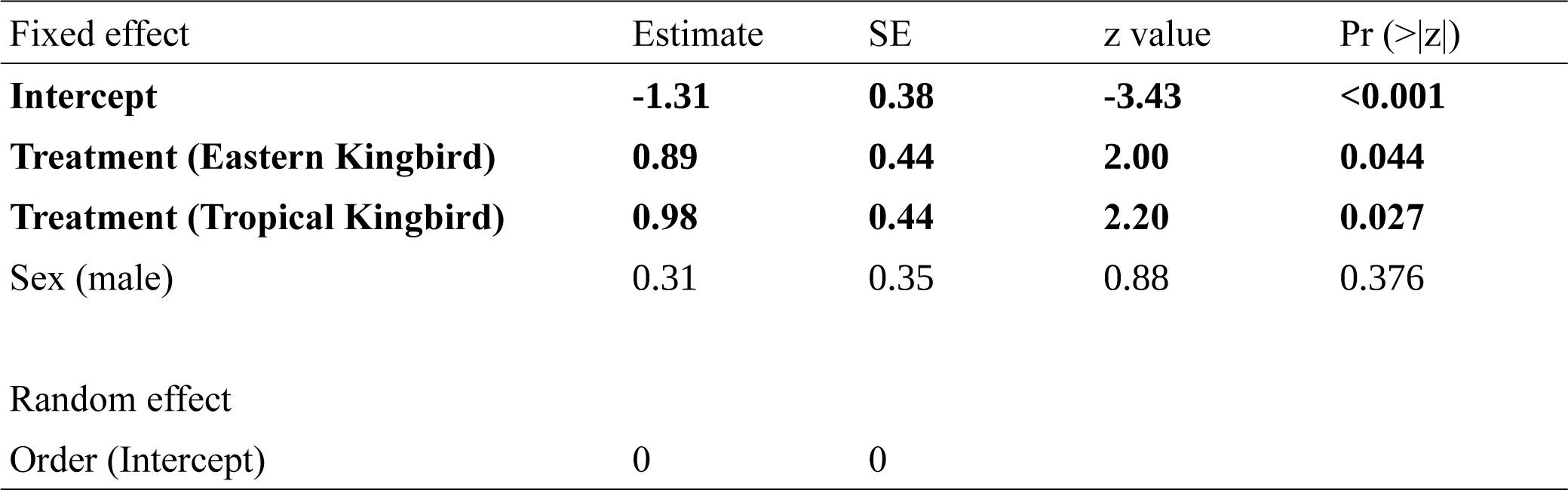
GLMM results of the effect of treatment (calls) and sex on proportion of butterflies changing their behaviour in response to calls in experiment 2. p<0.05 are bolded.

**Supplementary table 17:**
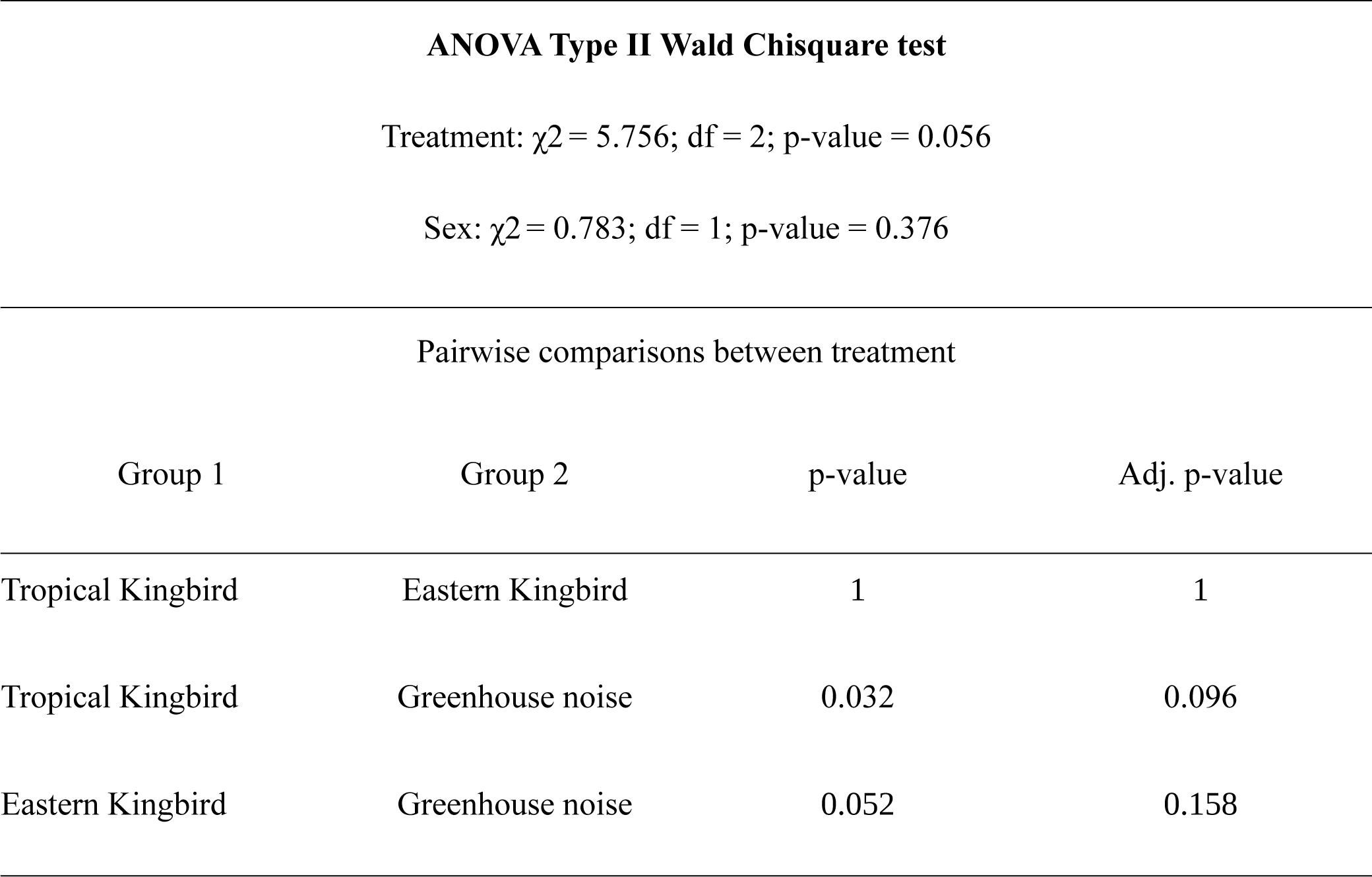
Pairwise differences in the proportion of individuals changing their behavioural state in response to the calls in experiment 2.

**Supplementary Table 18:**
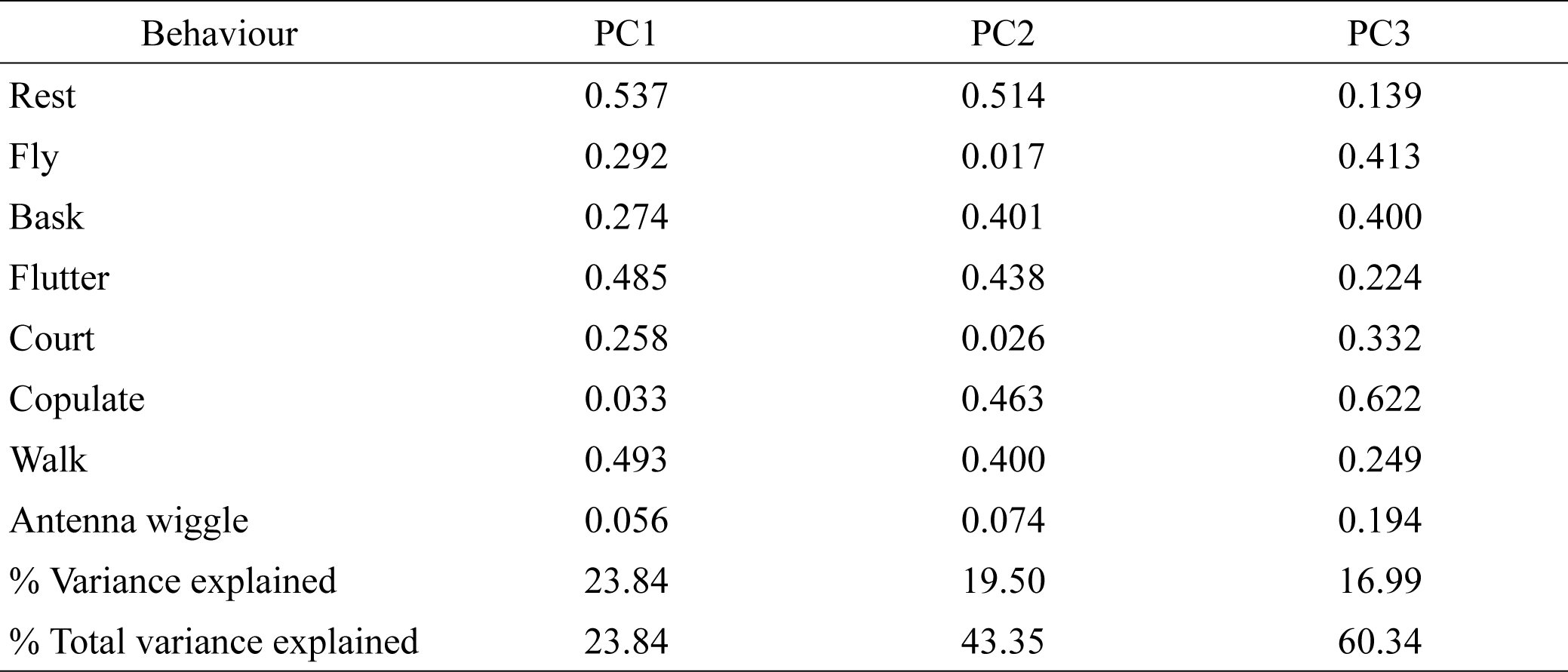
Loadings of each behaviour in Principal Component (PC) composite variables for males in a minute before, during and after calls in experiment 2.

**Supplementary Table 19:**
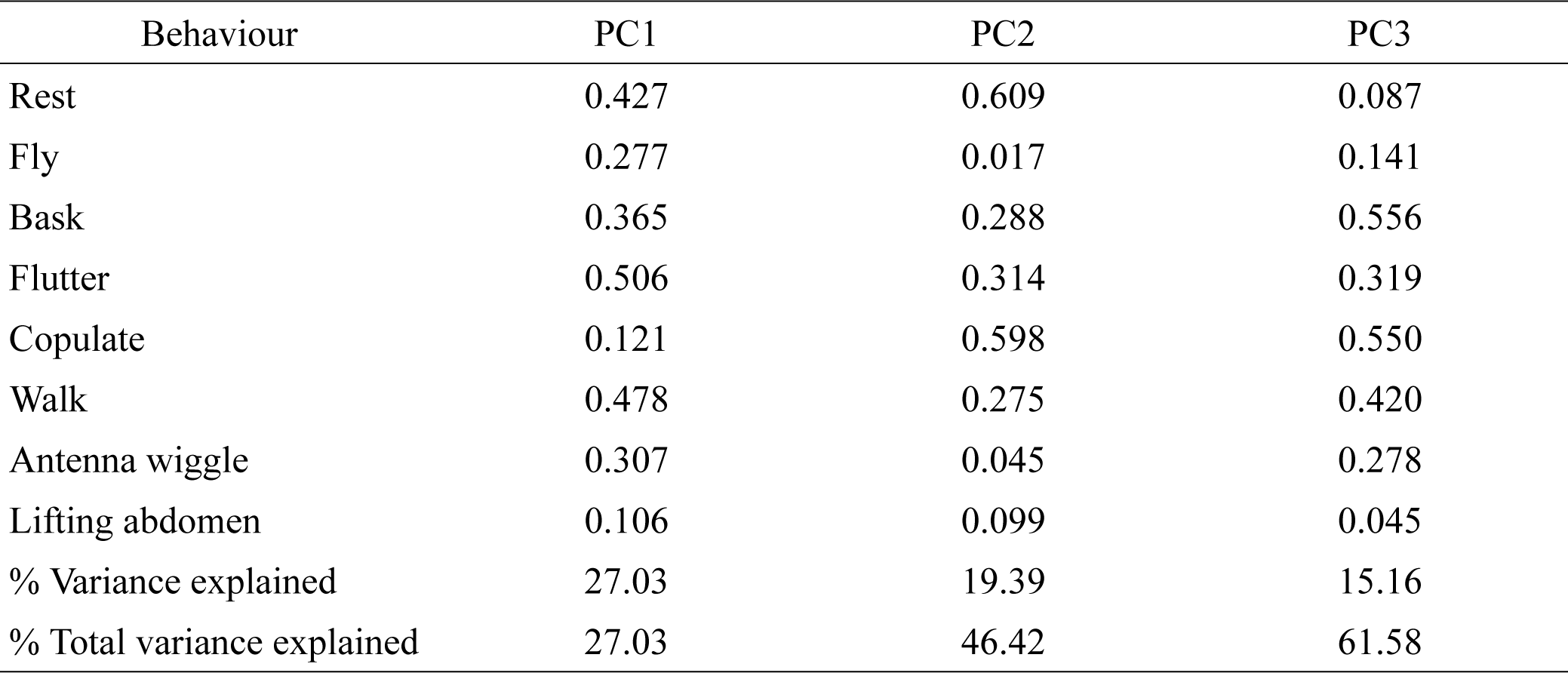
Loadings of each behaviour in Principal Component (PC) composite variables for females in a minute before, during and after calls in experiment 2.

**Supplementary Table 20:**
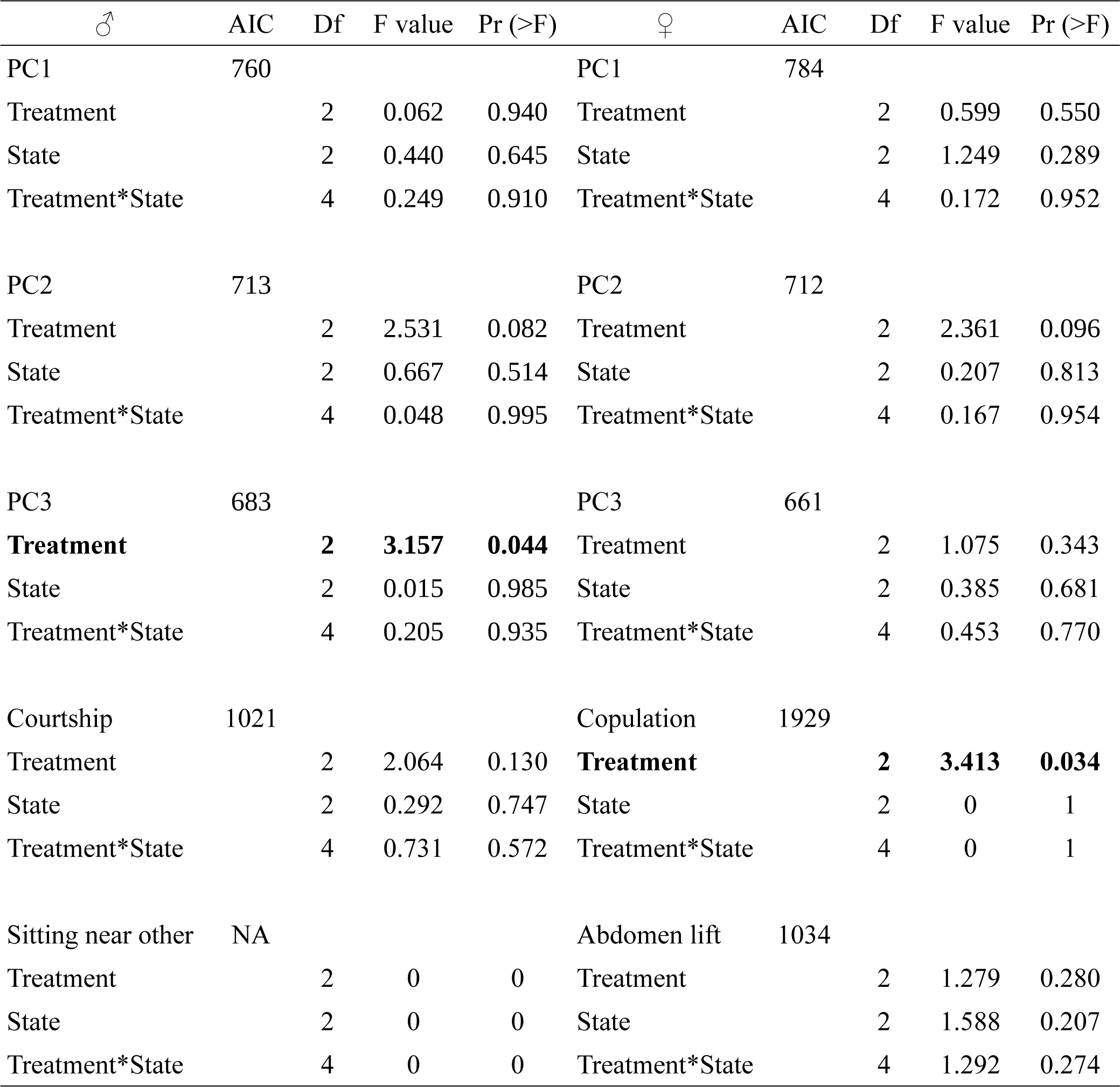
Effect of treatment (Tropical Kingbird, Eastern Kingbird, and Greenhouse background noise calls), state (before, during, and after call) and their interaction on male PC1, PC2, PC3, *courtship, sit near* and female PC1, PC2, PC3, *copulation*, *abdomen lifting* behaviours in experiment 2. p<0.05 bolded. For male *sit near* behaviour, there were zero occurrences.

**Supplementary Table 21:**
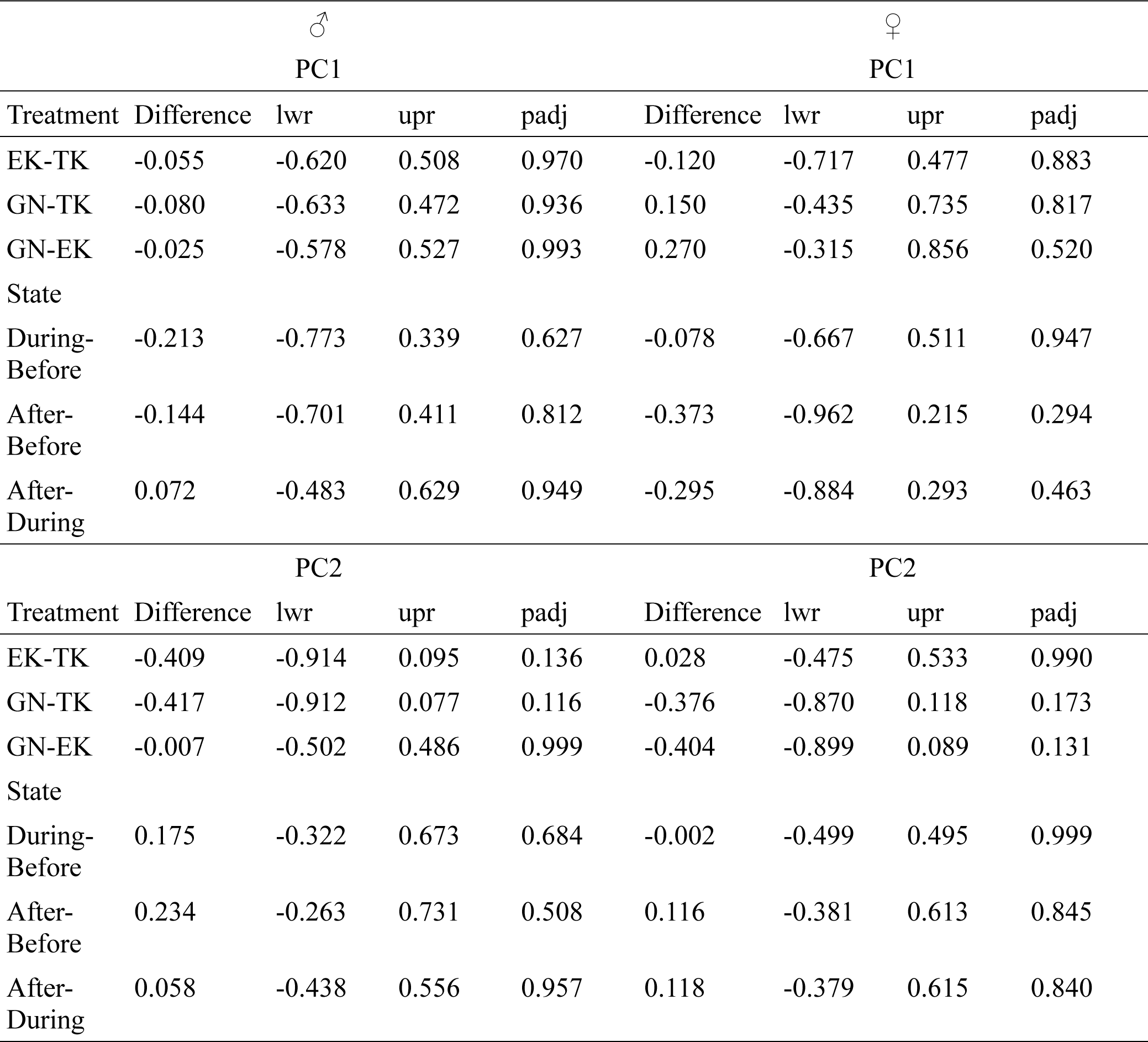
ANOVA post-hoc test results with PC as dependent variables and the treatments (calls), state (before, during, and after) and their interaction as response variables for males and females in a minute before, during, and after calls in experiment 2. EK= Eastern kingbird; TK=Tropical kingbird; GN=Greenhouse background noise; Difference= pairwise difference; lwr=lower range; upr=upper range; padj= adjusted p-value.

**Supplementary Table 22:**
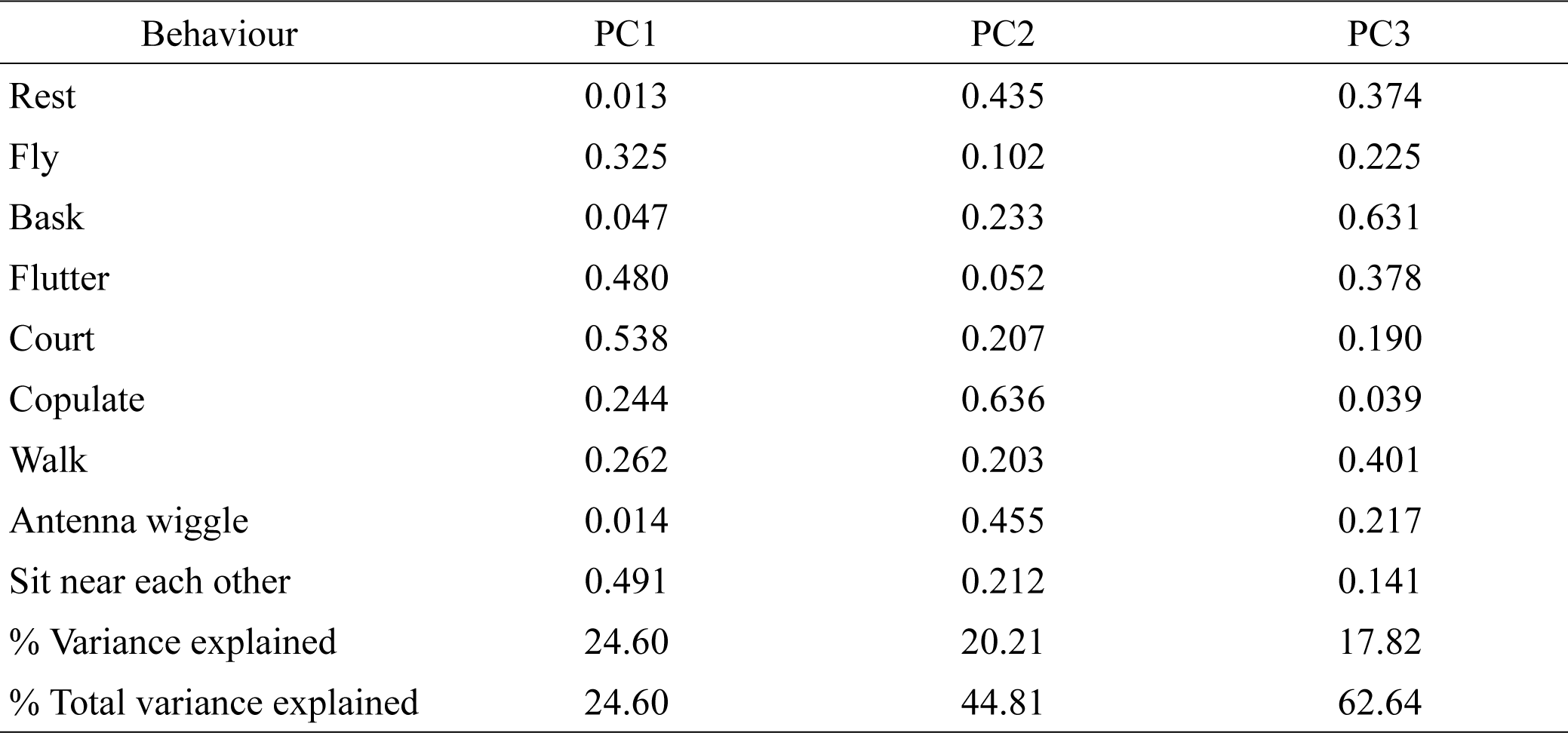
Loadings of each behaviour in Principal Component (PC) composite variables for males in 14 minutes before and after calls in experiment 2.

**Supplementary Table 23:**
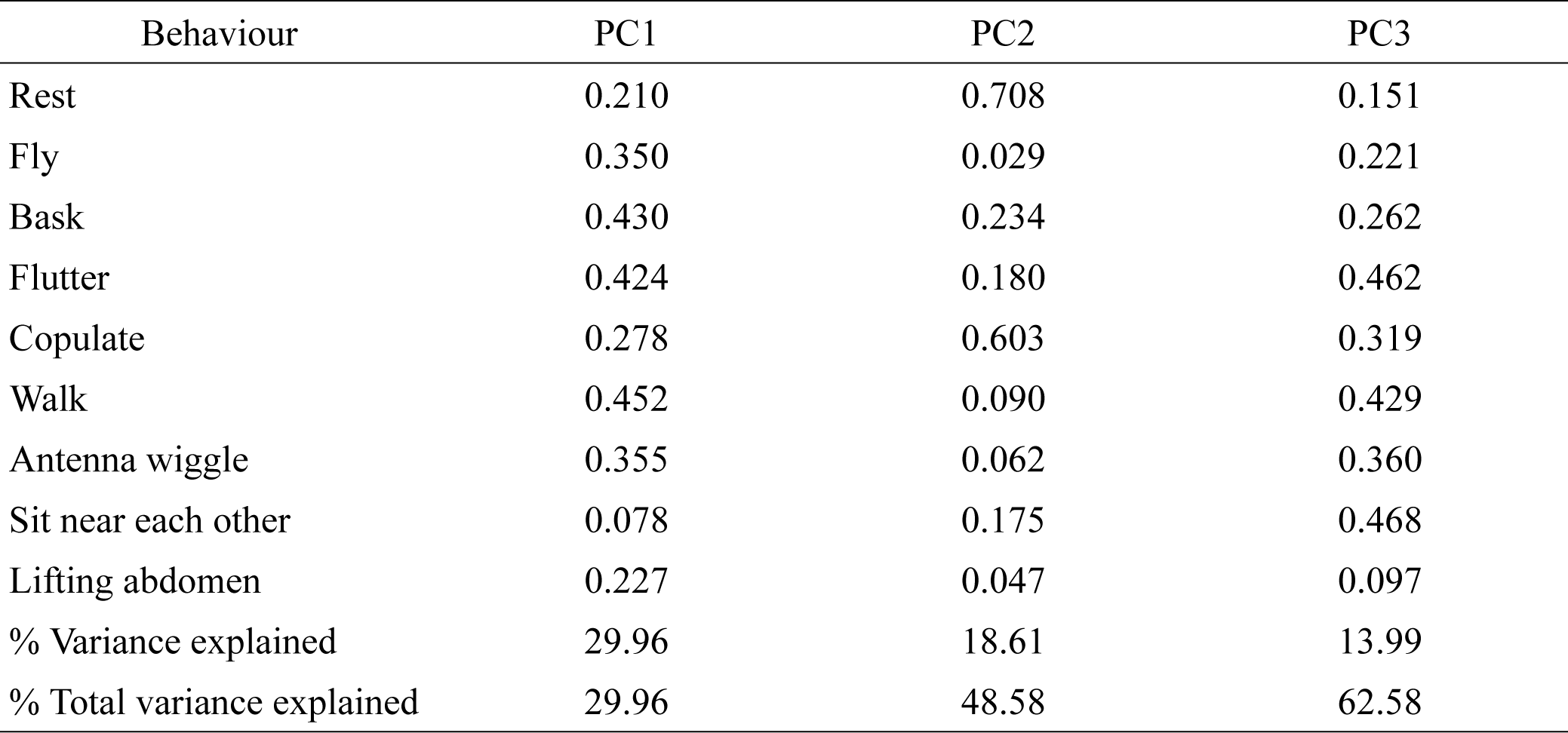
Loadings of each behaviour in Principal Component (PC) composite variables for females in 14 minutes before and after calls in experiment 2.

**Supplementary Table 24:**
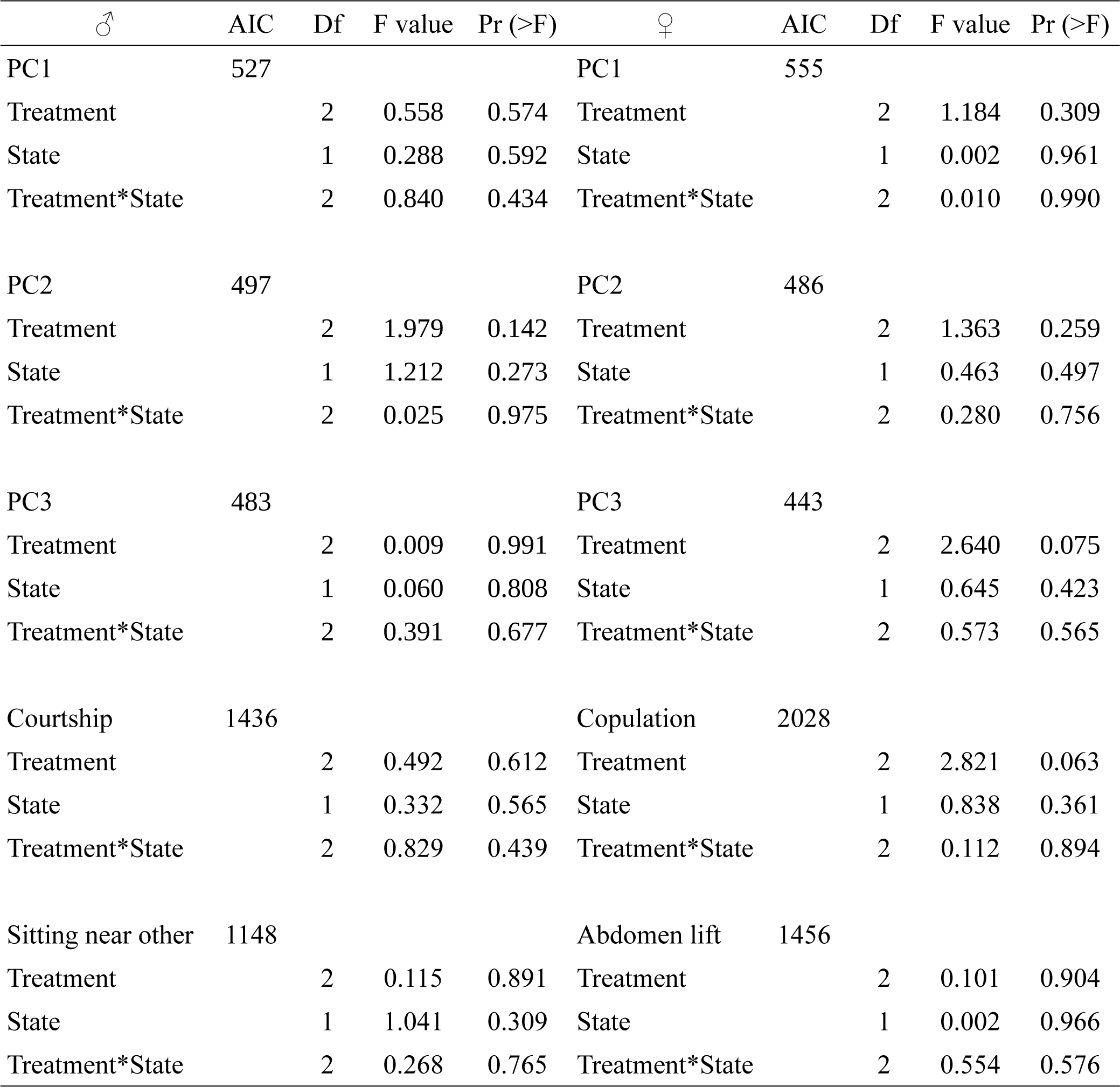
Effect of treatment (Rufous-tailed Jacamar, Eastern Kingbird, Toco Toucan and Greenhouse background noise calls), state (before call and after call) and their interaction on 14 minute behaviors before and after call, and male PC1, PC2, PC3, *courtship, copulation, sit near* and female PC1, PC2, PC3, *abdomen lifting* behaviours in experiment 2.

**Supplementary Table 25:**
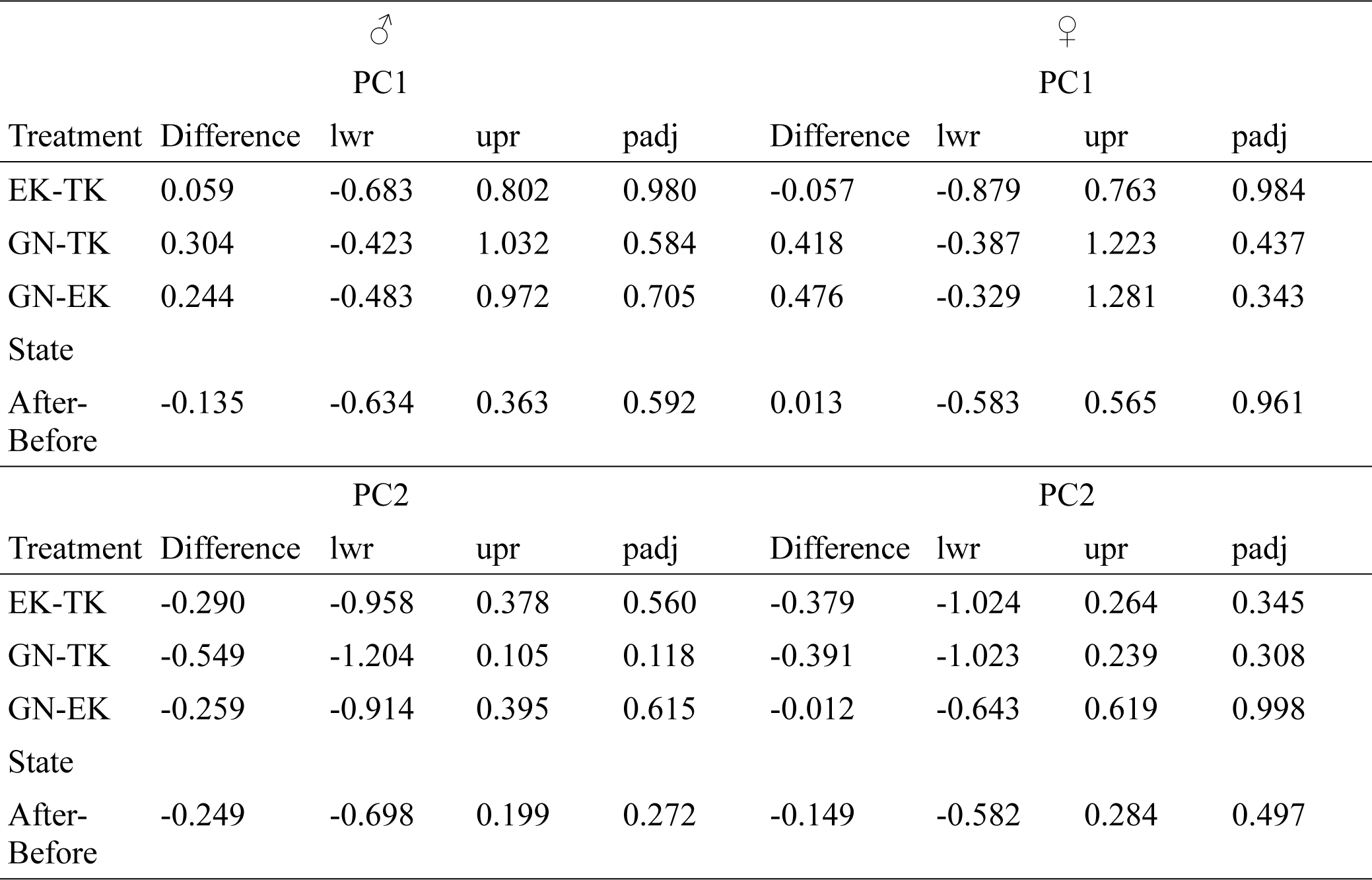
ANOVA post-hoc test results with PC as dependent variables and the treatments (calls), state (before and after) and their interaction as response variables for males and females in 14 minutes before, during, and after calls in experiment 2. EK= Eastern kingbird; TK=Tropical kingbird; GN=Greenhouse background noise; Difference= pairwise difference; lwr=lower range; upr=upper range; padj= adjusted p-value.

## References

Acharya, L., & McNeil, J. N. (1998). Predation risk and mating behavior: The responses of moths to bat-like ultrasound. Behavioral Ecology, 9(6), 552–558. 10.1093/beheco/9.6.552

Apfelbach, R., Blanchard, C. D., Blanchard, R. J., Hayes, R. A., & McGregor, I. S. (2005). The effects of predator odors in mammalian prey species: A review of field and laboratory studies. Neuroscience & Biobehavioral Reviews, 29(8), 1123– 1144. 10.1016/j.neubiorev.2005.05.005

Bernal, X. E., Stanley Rand, A., & Ryan, M. J. (2007). Sexual differences in the behavioral response of Túngara Frogs, *Physalaemus pustulosus*, to cues associated with increased predation risk: Sex differences in behavioral response to increased predation risk. Ethology, 113(8), 755–763. 10.1111/j.1439-0310.2007.01374.x

Blake, J. G., & Loiselle, B. A. (1992). Fruits in the diets of neotropical migrant birds in Costa Rica. Biotropica, 24(2), 200. 10.2307/2388674

Cantwell, L. R., & Forrest, T. G. (2013). Response of *Anolis sagrei* to acoustic calls from predatory and nonpredatory birds. Journal of Herpetology, 47(2), 293–298. 10.1670/11-184

Chai, P. (1986). Field observations and feeding experiments on the responses of rufous-tailed jacamars (*Galbula ruficauda*) to free-flying butterflies in a tropical rainforest. Biological Journal of the Linnean Society, 29(3), 161–189. 10.1111/j.1095-8312.1986.tb01772.x

Chai, P., & Srygley, R. B. (1990). Predation and the flight, morphology, and temperature of neotropical rain-forest butterflies. The American Naturalist, 135(6), 748–765. 10.1086/285072

Chouteau, M., Llaurens, V., Piron-Prunier, F., & Joron, M. (2017). Polymorphism at a mimicry supergene maintained by opposing frequency-dependent selection pressures. Proceedings of the National Academy of Sciences, 114(31), 8325–8329. 10.1073/pnas.1702482114

Conner, W. E., & Corcoran, A. J. (2012). Sound strategies: The 65-Million-Year-Old battle between bats and insects. Annual Review of Entomology, 57(1), 21–39. 10.1146/annurev-ento-121510-133537

Core Team, R. (2023). R: A Language and Environment for Statistical Computing. R Foundation for Statistical Computing. https://www.R-project.org/

Curlis, J. D., Macklem, D. C., Davis, R., & Cox, C. L. (2016). Sex-specific antipredator response to auditory cues in the black spiny-tailed iguana. Journal of Zoology, 299(1), 68–74. 10.1111/jzo.12326

Deecke, V. B., Slater, P. J. B., & Ford, J. K. B. (2002). Selective habituation shapes acoustic predator recognition in harbour seals. Nature, 420(6912), 171–173. 10.1038/nature01030

Dell’Aglio, D. D., Mena, S., Mauxion, R., McMillan, W. O., & Montgomery, S. H. (2022). Divergence in *Heliconius* flight behaviour is associated with local adaptation to different forest structures. Journal of Animal Ecology, 91(4), 727– 737. 10.1111/1365-2656.13675

Edomwande, C., & Barbosa, F. (2020). The influence of predation risk on mate signaling and mate choice in the lesser waxmoth *Achroia grisella*. Scientific Reports, 10(1), 524. 10.1038/s41598-020-57481-1

Engler-Chaouat, H. S., & Gilbert, L. E. (2007). De novo Synthesis vs. Sequestration: Negatively correlated metabolic traits and the evolution of host plant specialization in cyanogenic butterflies. Journal of Chemical Ecology, 33(1), 25–42. 10.1007/s10886-006-9207-8

Faure, P. A., & Hoy, R. R. (2000). The sounds of silence: Cessation of singing and song pausing are ultrasound-induced acoustic startle behaviors in the katydid *Neoconocephalus ensiger* (Orthoptera; Tettigoniidae). Journal of Comparative Physiology A: Sensory, Neural, and Behavioral Physiology, 186(2), 129–142. 10.1007/s003590050013

Finkbeiner, S. D., Briscoe, A. D., & Reed, R. D. (2012). The benefit of being a social butterfly: Communal roosting deters predation. Proceedings of the Royal Society B: Biological Sciences, 279(1739), 2769–2776. 10.1098/rspb.2012.0203

Fisher, K. A., & Stankowich, T. (2018). Antipredator strategies of striped skunks in response to cues of aerial and terrestrial predators. Animal Behaviour, 143, 25–34. 10.1016/j.anbehav.2018.06.023

Fitzpatrick, J. W. (1980). Foraging behavior of neotropical tyrant flycatchers. The Condor, 82(1), 43–57. 10.2307/1366784

Fournier, J. P., Dawson, J. W., Mikhail, A., & Yack, J. E. (2013). If a bird flies in the forest, does an insect hear it? Biology Letters, 9(5), 20130319. 10.1098/rsbl.2013.0319

Gall, L. F. (1984). The effects of capturing and marking on subsequent activity in *Boloria acrocnema* (Lepidoptera: Nymphalidae), with a comparison of different numerical models that estimate population size. Biological Conservation, 28(2), 139–154. 10.1016/0006-3207(84)90032-6

Greene, E., & Meagher, T. (1998). Red squirrels, *Tamiasciurus hudsonicus*, produce predator-class specific alarm calls. Animal Behaviour, 55(3), 511–518. 10.1006/anbe.1997.0620

Hines, H. M., Counterman, B. A., Papa, R., Albuquerque De Moura, P., Cardoso, M. Z., Linares, M., Mallet, J., Reed, R. D., Jiggins, C. D., Kronforst, M. R., & McMillan, W. O. (2011). Wing patterning gene redefines the mimetic history of *Heliconius* butterflies. Proceedings of the National Academy of Sciences, 108(49), 19666–19671. 10.1073/pnas.1110096108

Jacobs, D. S., Ratcliffe, J. M., & Fullard, J. H. (2008). Beware of bats, beware of birds: The auditory responses of eared moths to bat and bird predation. Behavioral Ecology, 19(6), 1333–1342. 10.1093/beheco/arn071

Klein, A. L., & De Araújo, A. M. (2010). Courtship behavior of *Heliconius erato phyllis* (Lepidoptera, Nymphalidae) towards virgin and mated females: Conflict between attraction and repulsion signals? Journal of Ethology, 28(3), 409–420. 10.1007/s10164-010-0209-1

Lane, K. A., Lucas, K. M., & Yack, J. E. (2008). Hearing in a diurnal, mute butterfly,Morpho peleides (Papilionoidea, Nymphalidae). The Journal of Comparative Neurology, 508(5), 677–686. 10.1002/cne.21675

Langham, G. M. (2004). Specialized avian predators repeatedly attack novel color morphs of *Heliconius* butterflies. Evolution, 58(12), 2783–2787.

Langham, G. M. (2006). Rufous-tailed jacamars and aposematic butterflies: Do older birds attack novel prey? Behavioral Ecology, 17(2), 285–290. 10.1093/beheco/arj027

Lea, A. J., & Blumstein, D. T. (2011). Age and sex influence marmot antipredator behavior during periods of heightened risk. Behavioral Ecology and Sociobiology, 65(8), 1525–1533. 10.1007/s00265-011-1162-x

Lind, J., & Cresswell, W. (2005). Determining the fitness consequences of antipredation behavior. Behavioral Ecology, 16(5), 945–956. 10.1093/beheco/ari075

Lohrey, A. K., Clark, D. L., Gordon, S. D., & Uetz, G. W. (2009). Antipredator responses of wolf spiders (Araneae: Lycosidae) to sensory cues representing an avian predator. Animal Behaviour, 77(4), 813–821. 10.1016/j.anbehav.2008.12.025

Lucas, K. M., Mongrain, J. K., Windmill, J. F. C., Robert, D., & Yack, J. E. (2014). Hearing in the crepuscular owl butterfly (*Caligo eurilochus*, Nymphalidae). Journal of Comparative Physiology A, 200(10), 891–898. 10.1007/s00359-014-0933-z

Lucas, K. M., Windmill, J. F. C., Robert, D., & Yack, J. E. (2009). Auditory mechanics and sensitivity in the tropical butterfly *Morpho peleides* (Papilionoidea, Nymphalidae). Journal of Experimental Biology, 212(21), 3533–3541. 10.1242/jeb.032425

Mallet, J., & Gilbert, L. E. (1995). Why are there so many mimicry rings? Correlations between habitat, behaviour and mimicry in *Heliconius* butterflies. Biological Journal of the Linnean Society, 55, 159–180.

Mikhail, A., Lewis, J. E., & Yack, J. E. (2018). What does a butterfly hear? Physiological characterization of auditory afferents in *Morpho peleides* (Nymphalidae). Journal of Comparative Physiology A, 204(9–10), 791–799. 10.1007/s00359-018-1280-2

Morton, E. S. (1971). Food and migration habits of the Eastern kingbird in Panama. The Auk, 88(4), 925–926. 10.2307/4083855

Olofsson, M., Eriksson, S., Jakobsson, S., & Wiklund, C. (2012). Deimatic display in the European swallowtail butterfly as a secondary defence against attacks from Great tits. PLoS ONE, 7(10), e47092. 10.1371/journal.pone.0047092

Olofsson, M., Vallin, A., Jakobsson, S., & Wiklund, C. (2011). Winter predation on two species of hibernating butterflies: Monitoring rodent attacks with infrared cameras. Animal Behaviour, 81(3), 529–534. 10.1016/j.anbehav.2010.12.012

Palmer, M. S., & Packer, C. (2021). Reactive anti-predator behavioral strategy shaped by predator characteristics. PLOS ONE, 16(8), e0256147. 10.1371/journal.pone.0256147

Pinheiro, C. E. G. (1996). Palatablility and escaping ability in neotropical butterflies: Tests with wild kingbirds (*Tyrannus melancholicus*, Tyrannidae). Biological Journal of the Linnean Society, 59(4), 351–365. 10.1111/j.1095-8312.1996.tb01471.x

Pinheiro, C. E. G. (2011). On the evolution of warning coloration, Batesian and Müllerian mimicry in Neotropical butterflies: The role of jacamars (Galbulidae) and tyrant-flycatchers (Tyrannidae). Journal of Avian Biology, 42(4), 277–281. 10.1111/j.1600-048X.2011.05435.x

Pinheiro, C. E. G., & Campos, V. C. (2019). The responses of wild jacamars (*Galbula ruficauda*, Galbulidae) to aposematic, aposematic and cryptic, and cryptic butterflies in central Brazil. Ecological Entomology, 44(4), 441–450. 10.1111/een.12723

Pinheiro, C. E. G., & Cintra, R. (2017). Butterfly predators in the neotropics: Which birds are involved? Journal of the Lepidopterists’ Society, 71(2), 109–114. 10.18473/lepi.71i2.a5

Pinheiro De Castro, É. C., Zagrobelny, M., Zurano, J. P., Zikan Cardoso, M., Feyereisen, R., & Bak, S. (2019). Sequestration and biosynthesis of cyanogenic glucosides in passion vine butterflies and consequences for the diversification of their host plants. Ecology and Evolution, 9, 5079–5093. 10.1002/ece3.5062

Prakash, H., Greif, S., Yovel, Y., & Balakrishnan, R. (2021). Acoustically eavesdropping bat predators take longer to capture katydid prey signalling in aggregation. Journal of Experimental Biology, 224(10), jeb233262. 10.1242/jeb.233262

Rather, P. A., Herzog, A. E., Ernst, D. A., & Westerman, E. L. (2022). Effect of experience on mating behaviour in male *Heliconius melpomene* butterflies. Animal Behaviour, 183, 139–149. 10.1016/j.anbehav.2021.11.004

Ribarič, D., & Gogala, M. (1996). Acoustic behaviour of some butterfly species of the Genus *Erebia* (Lepidoptera: Satyridae). Acta Entomologica Slovenica, 4(1), 5–12.

Robertson, D. N., Sullivan, T. J., & Westerman, E. L. (2020). Lack of sibling avoidance during mate selection in the butterfly *Bicyclus anynana*. Behavioural Processes, 173, 104062. 10.1016/j.beproc.2020.104062

Rojas, B., Mappes, J., & Burdfield-Steel, E. (2019). Multiple modalities in insect warning displays have additive effects against wild avian predators. Behavioral Ecology and Sociobiology, 73(3), 37. 10.1007/s00265-019-2643-6

Rosen, M. J., Levin, E. C., & Hoy, R. R. (2009). The cost of assuming the life history of a host: Acoustic startle in the parasitoid fly *Ormia ochracea*. Journal of Experimental Biology, 212(24), 4056–4064. 10.1242/jeb.033183

Sitvarin, M. I., & Rypstra, A. L. (2012). Sex-Specific response of *Pardosa milvina* (Araneae: Lycosidae) to experience with a chemotactile predation cue. Ethology, 118(12), 1230–1239. 10.1111/eth.12029

Srygley, R. B. (1994). Locomotor mimicry in butterflies? The associations of positions of Centres of Mass among Groups of Mimetic, Unprofitable Prey. Philosophical Transactions: Biological Sciences, 343(1304), 145–155.

Sun, P., Mhatre, N., Mason, A. C., & Yack, J. E. (2018). In that vein: Inflated wing veins contribute to butterfly hearing. Biology Letters, 14(10), 20180496. 10.1098/rsbl.2018.0496

Swihart, S. L. (1967). Hearing in Butterflies (Nymphalidae: *Heliconius*, Ageronia). Journal of Insect Physiology, 13, 469–476.

Thaker, M., Lima, S. L., & Hews, D. K. (2009). Alternative antipredator tactics in tree lizard morphs: Hormonal and behavioural responses to a predator encounter. Animal Behaviour, 77(2), 395–401. 10.1016/j.anbehav.2008.10.014

Torsekar, V. R., Isvaran, K., & Balakrishnan, R. (2019). Is the predation risk of mate-searching different between the sexes? Evolutionary Ecology, 33(3), 329–343. 10.1007/s10682-019-09982-3

Triblehorn, J. D., Ghose, K., Bohn, K., Moss, C. F., & Yager, D. D. (2008). Free-flight encounters between praying mantids (*Parasphendale agrionina*) and bats (*Eptesicus fuscus*). Journal of Experimental Biology, 211(4), 555–562. 10.1242/jeb.005736

Umbers, K. D. L., & Mappes, J. (2015). Postattack deimatic display in the mountain katydid, *Acripeza reticulata*. Animal Behaviour, 100, 68–73. 10.1016/j.anbehav.2014.11.009

Vallin, A., Jakobsson, S., Lind, J., & Wiklund, C. (2006). Crypsis versus intimidation— Anti-predation defence in three closely related butterflies. Behavioral Ecology and Sociobiology, 59(3), 455–459. 10.1007/s00265-005-0069-9

Westerman, E. L., Chirathivat, N., Schyling, E., & Monteiro, A. (2014). Mate preference for a phenotypically plastic trait is learned, and may facilitate preference-phenotype matching. Evolution, 68(6), 1661–1670. 10.1111/evo.12381

Wickham, H. (2016). *ggplot2: Elegant Graphics for Data Analysis*. Springer-Verlag New York. https://ggplot2.tidyverse.org

Wormington, J. D., & Juliano, S. A. (2014). Hunger-dependent and sex-specific antipredator behaviour of larvae of a size-dimorphic mosquito. Ecological Entomology, 39(5), 548–555. 10.1111/een.12129

Zaguri, M., & Hawlena, D. (2020). Odours of non-predatory species help prey moderate their risk assessment. Functional Ecology, 34(4), 830–839. 10.1111/1365-2435.13509

