## supplementary material 1 for "Behavioural changes in aposematic *Heliconius melpomene* butterflies in response to their predatory bird calls"

### Table of Content

| Name | Brief description | Pg.no |
|---|---|---|
| Supplementary Figure 1 | Experimental design | 3 |
| Supplementary Figure 2 | Spectograms of calls | 4 |
| Supplementary Figure 3 | Proportion changing behaviour at start of call | 6 |
| Supplementary Figure 4 | Proportion changing behaviour at end of call | 7 |
| Supplementary Figure 5 | Long term effect of calls on PC1 and PC2 behavior in expt 1 | 8 |
| Supplementary Figure 6 | Long term effect of calls on PC1 and PC2 behavior in expt 2 | 9 |
| Supplementary Table 1 | GLMM results for behavioural change at start of call in expt 1 | 10 |
| Supplementary Table 2 | Pairwise difference in behavioural change at start of call in expt 1 | 11 |
| Supplementary Table 3 | GLMM results for behavioural change at end of call in expt 1 | 12 |
| Supplementary Table 4 | Pairwise difference in behavioural change at end of call in expt 1 | 13 |
| Supplementary Table 5 | PC loadings for males for 3 minutes in expt 1 | 14 |
| Supplementary Table 6 | PC loadings for females for 3 minutes in expt 1 | 15 |
| Supplementary Table 7 | ANOVA post-hoc test for 3 minutes in expt 1 | 16 |
| Supplementary Table 8 | PC loadings for males for 28 minutes in expt 1 | 17 |
| Supplementary Table 9 | PC loadings for females for 28 minutes in expt 1 | 18 |
| Supplementary Table 10 | ANOVA models for PC1, PC2, PC3, and inter-sexual behaviours for 28 minutes in expt 1 | 19 |
| Supplementary Table 11 | ANOVA post-hoc test for 28 minutes in expt 1 | 20 |
| Supplementary Table 12 | GLMM results for behavioural change at start of call in expt 2 | 21 |
| Supplementary Table 13 | Pairwise difference in behavioural change at start of call in expt 2 | 22 |
| Supplementary Table 14 | GLMM results for behavioural change at end of call in expt 2 | 23 |
| Supplementary Table 15 | Pairwise difference in behavioural change at end of call in expt 2 | 24 |
| Supplementary Table 16 | GLMM results for behavioural change in response to call in expt 2 | 25 |
| Supplementary Table 17 | Pairwise difference in behavioural change in response to call in expt 2 | 26 |
| Supplementary Table 18 | PC loadings for males for 3 minutes in expt 2 | 27 |
| Supplementary Table 19 | PC loadings for females for 3 minutes in expt 2 | 28 |

| | | |
|---|---|---|
| Supplementary Table 20 | ANOVA models for PC1, PC2, PC3, and inter-sexual behaviours for 3 minutes in expt 2 | 29 |
| Supplementary Table 21 | ANOVA post-hoc test for 3 minutes in expt 2 | 30 |
| Supplementary Table 22 | PC loadings for males for 28 minutes in expt 2 | 31 |
| Supplementary Table 23 | PC loadings for females for 28 minutes in expt 2 | 32 |
| Supplementary Table 24 | ANOVA models for PC1, PC2, PC3, and inter-sexual behaviours for 28 minutes in expt 2 | 33 |
| Supplementary Table 25 | ANOVA post-hoc test for 28 minutes in expt 2 | 34 |

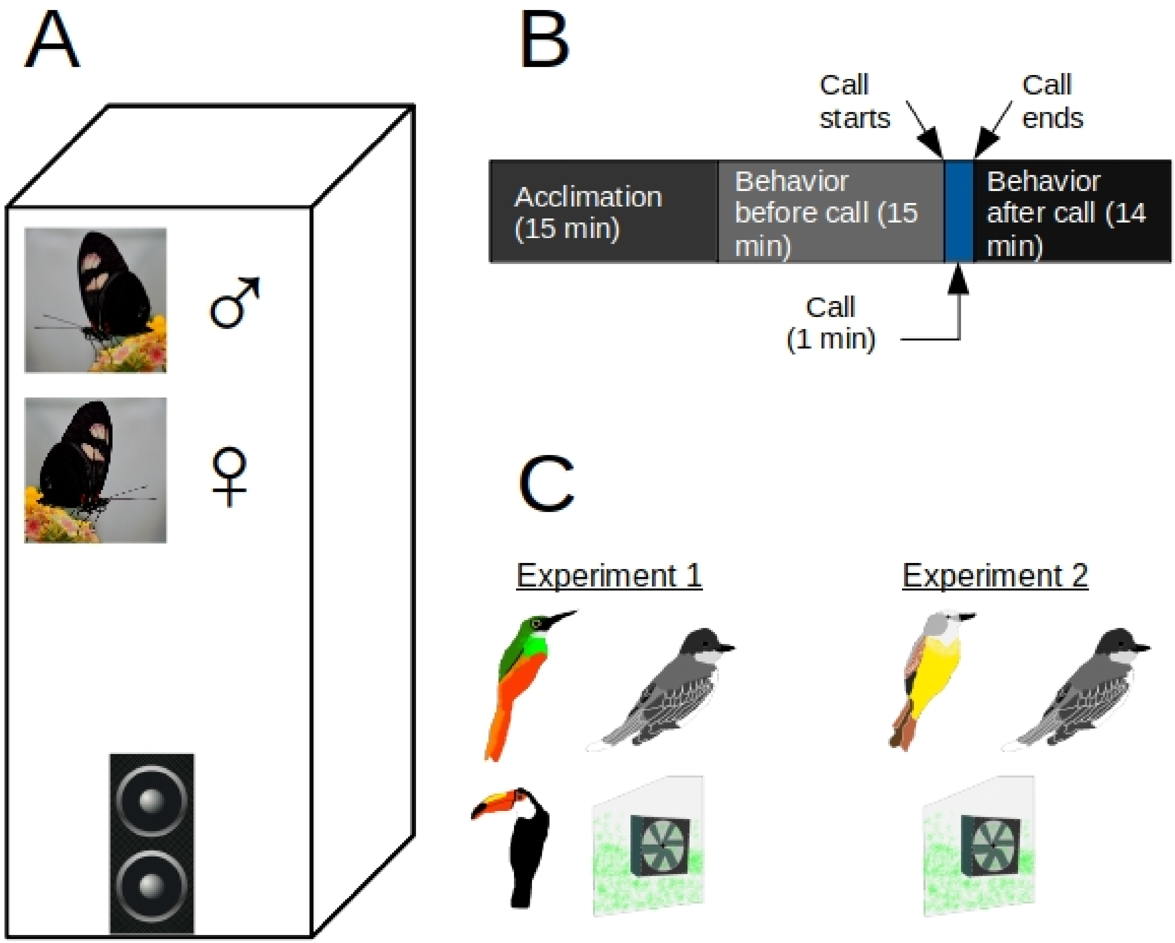

**Supplementary Figure 1:** Experimental design. A) 3-15-day male and female *H. m. plessini* butterflies were subjected in an experimental cage with a blue tooth speaker and a *Lantana spp.* plant during each experimental assay. B) The timeline of each assay conducted where the butterflies were acclimated for 15 minutes and their behaviours recorded for the next 30 minutes. During the 16$^{th}$ minute, a call was randomly played for a minute. C) The calls used in the two experiments in this study. Clockwise from top left in experiment 1: rufous-tailed jacamar, Eastern kingbird, greenhouse background noise, and toco toucan. Clockwise from top left in experiment 2: tropical kingbird, Eastern kingbird, and greenhouse background noise.

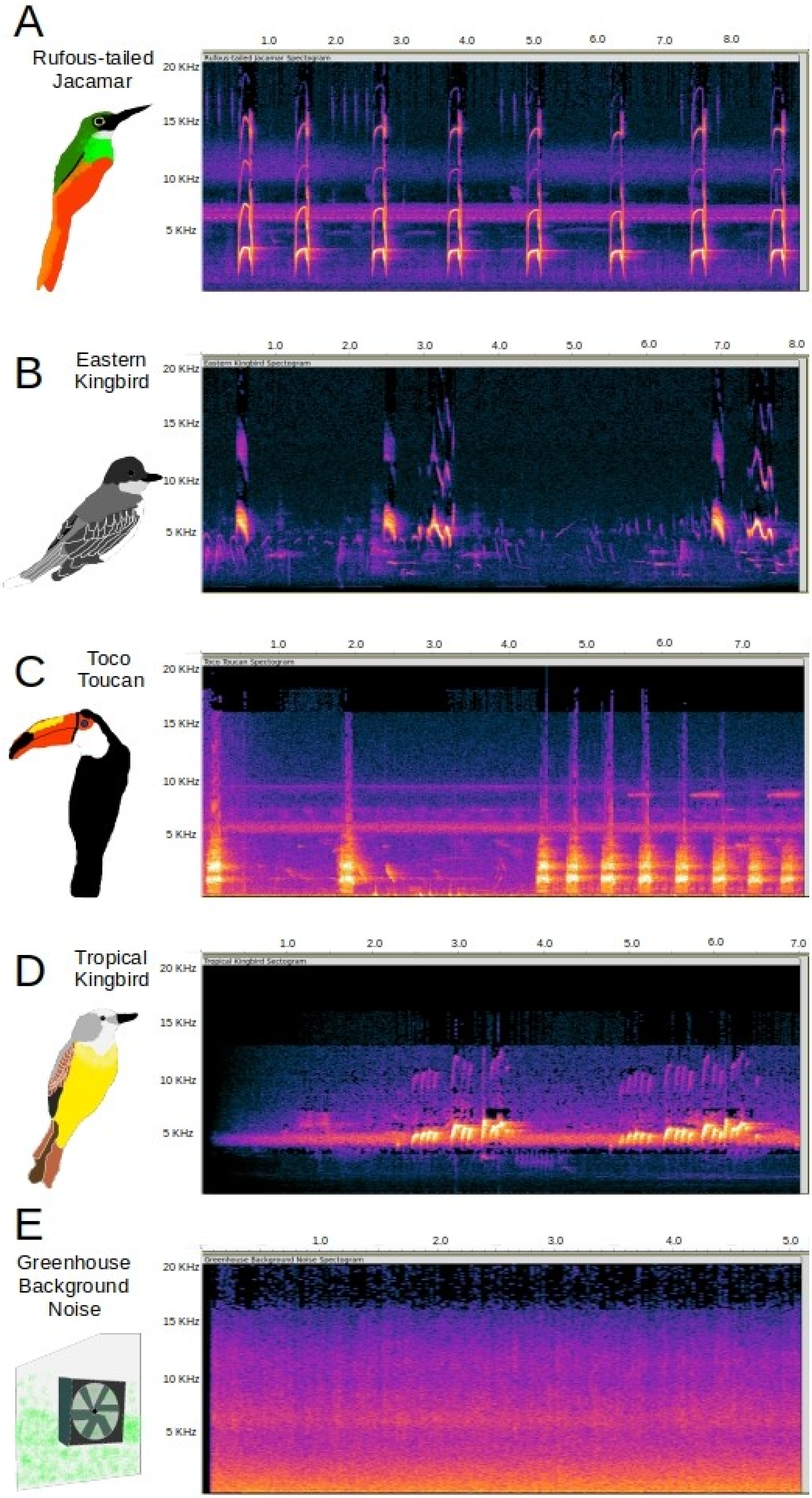

A
Rufous-tailed Jacamar
B
Eastern Kingbird
C
Toco Toucan
D
Tropical Kingbird
E
Greenhouse Background Noise
20 KHz
15 KHz
10 KHz
5 KHz
1.0
2.0
3.0
4.0
5.0
6.0
7.0
8.0

**Supplementary Figure 2:** Spectograms of the calls used during this study A) rufous-tailed jacamar; B) Eastern kingbird; C) toco toucan; D) tropical kingbird; E) greenhouse background noise.

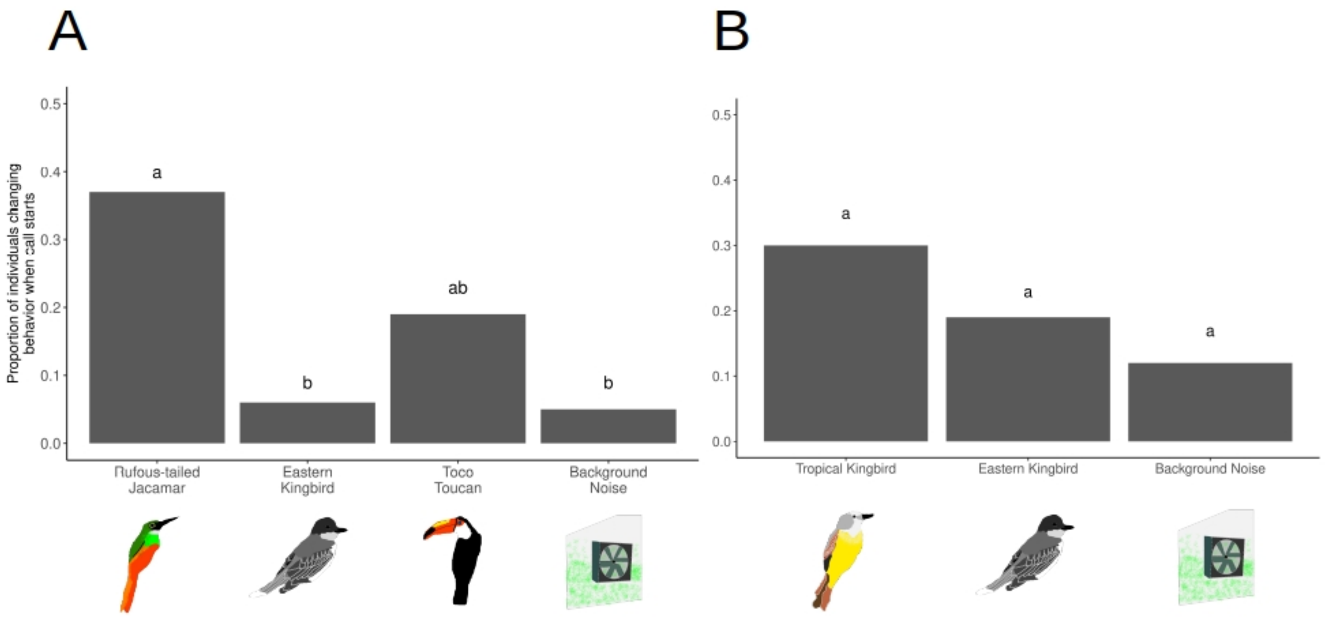

**Supplementary Figure 3:** Proportion of *H. m. plessini* individuals changing behaviour in response to the start of the calls (between before start and after start of calls) for A) experiment 1; B) experiment 2; Different letters on each bars indicate statistical significance at $p<0.05$.

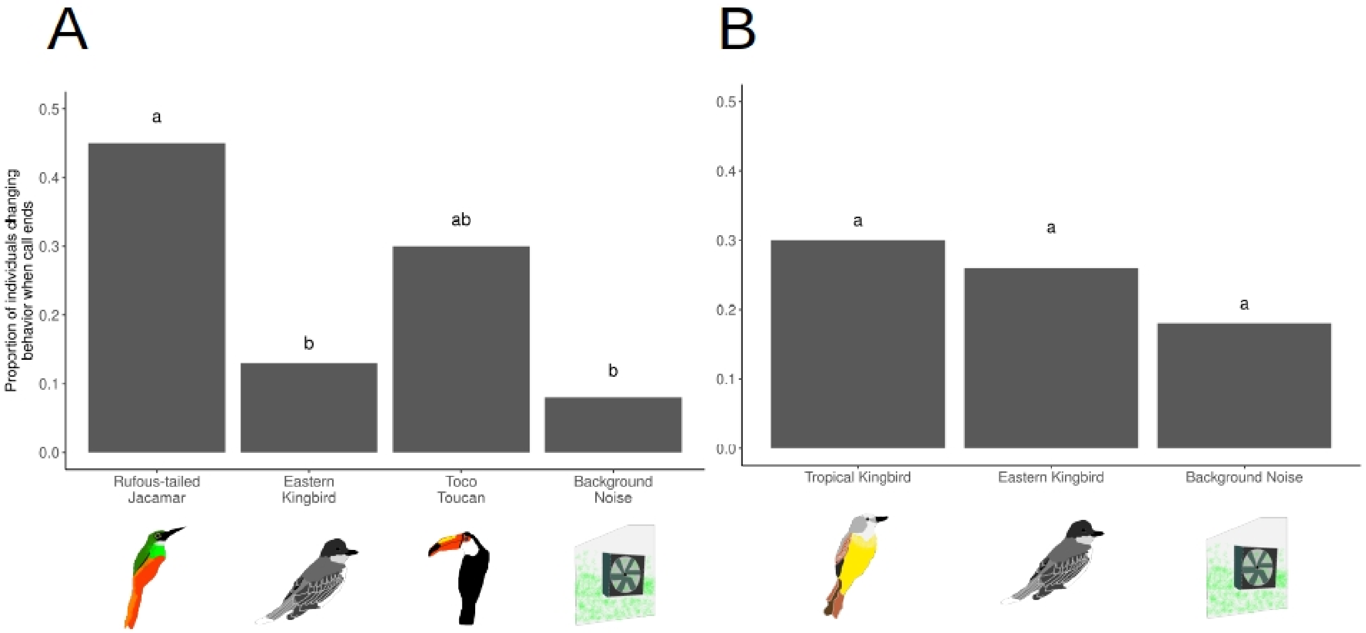

**Supplementary Figure 4:** Proportion of *H. m. plessini* individuals changing behaviour in response to the end of the calls (between before end and after end of calls) for A) experiment 1; B) experiment 2; Different letters on each bars indicate statistical significance at $p<0.05$.

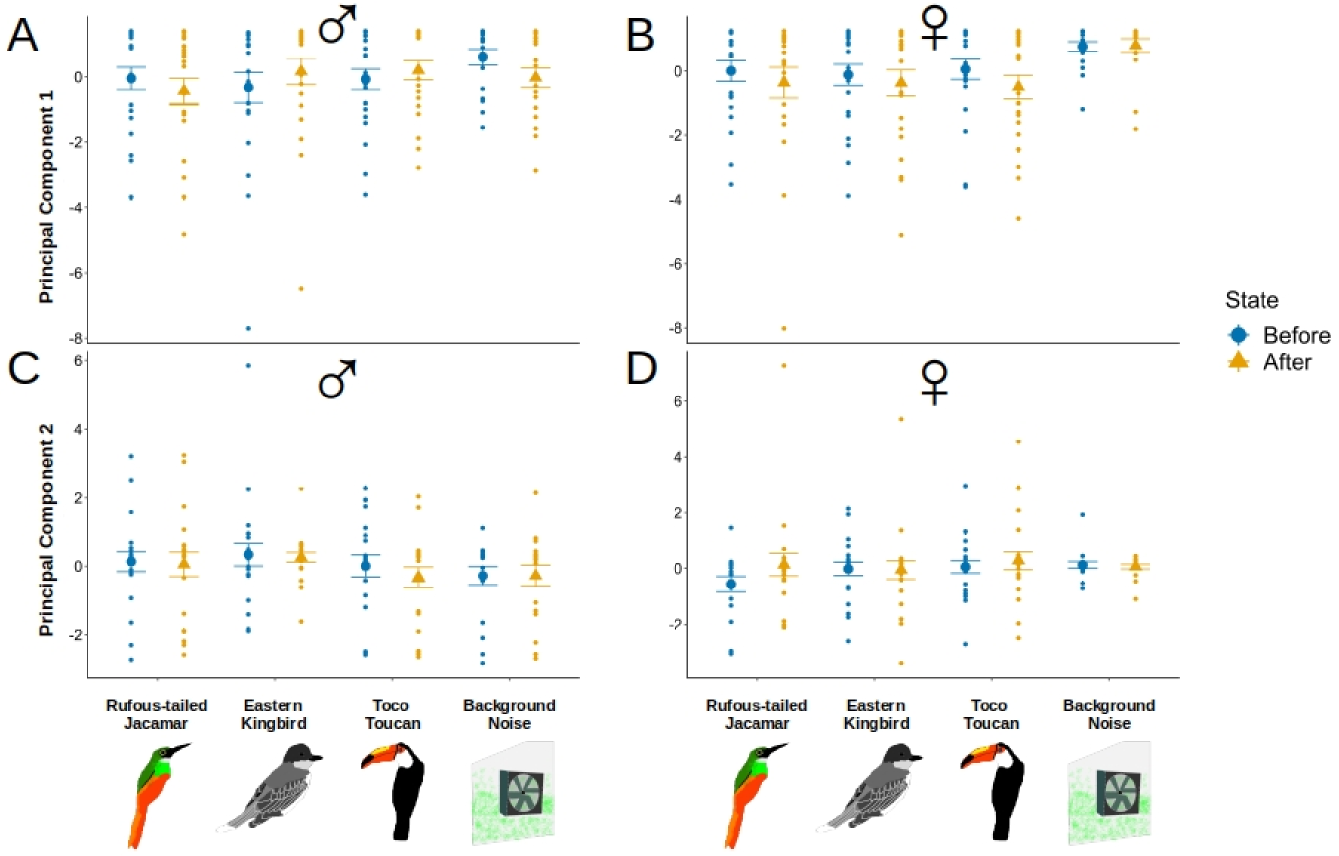

**Supplementary Figure 5:** Mean ± SE of principal component variables for male and female *H. m. plessini* for 14 minutes before, and after calls. A) PC 1 in males for experiment 1; B) PC 1 in females for experiment 1; C) PC 2 in males for experiment 1; D) PC 2 in females for experiment 1. None of them are significantly different from each other.

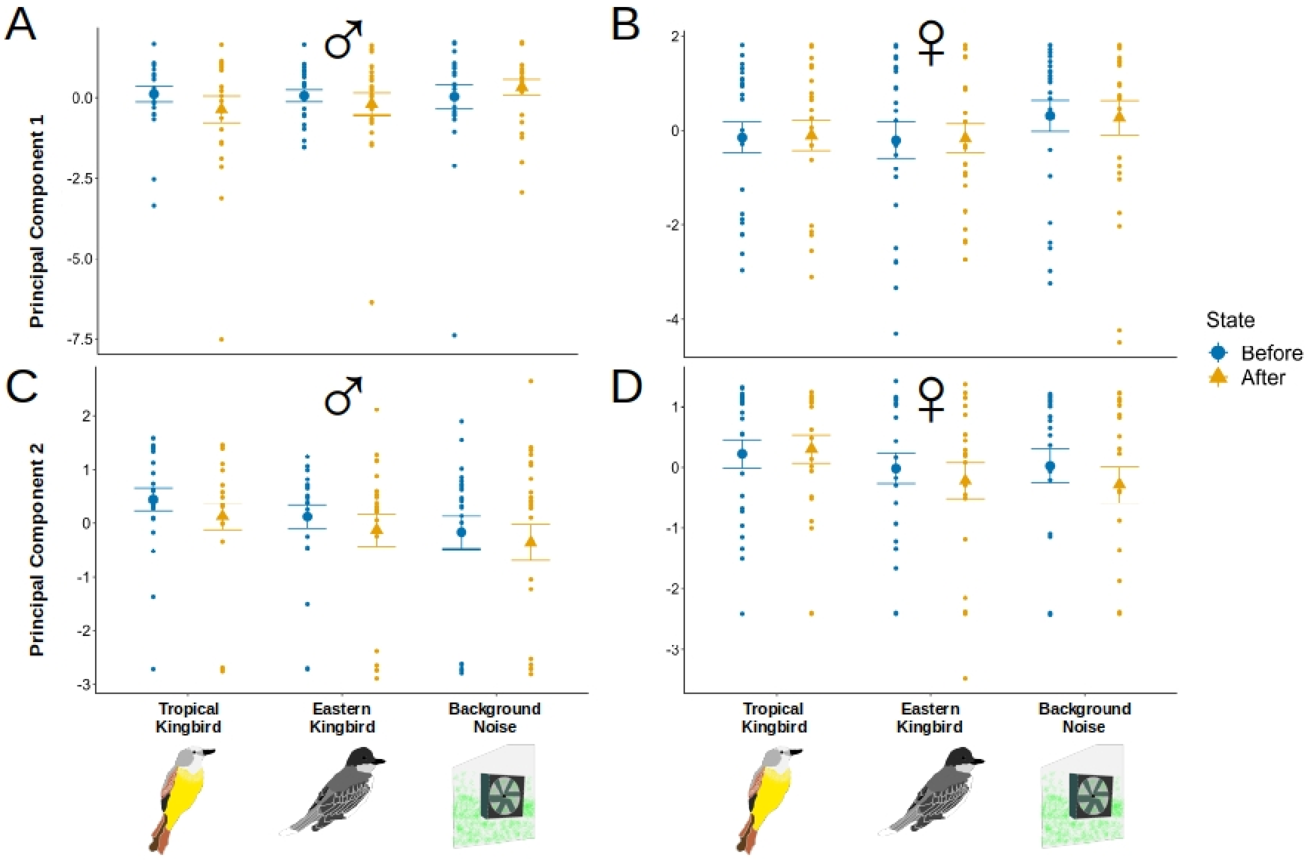

**Supplementary Figure 6:** Mean ± SE of principal component variables for male and female *H. m. plessini* for 14 minutes before, and after calls. A) PC 1 in males for experiment 2; B) PC 1 in females for experiment 2; C) PC 2 in males for experiment 2; D) PC 2 in females for experiment 2. None of them are significantly different from each other.

**Supplementary Table 1: GLMM results on the effect of treatment (calls) and sex on proportion of butterflies changing their behaviour at the start of calls in experiment 1. p<0.05 are bolded**

| Fixed effect | Estimate | SE | z value | Pr (>\|z\|) |
|---|---|---|---|---|
| **Intercept** | **-2.97** | **0.76** | **-3.88** | **<0.001** |
| **Treatment (Rufous-tailed Jacamar)** | **2.30** | **0.78** | **2.91** | **<0.001** |
| Treatment (Eastern Kingbird) | 0.17 | 0.94 | 0.18 | 0.856 |
| Treatment (Toco Toucan) | 1.42 | 0.81 | 1.73 | 0.082 |
| Sex (male) | 0.26 | 0.42 | 0.62 | 0.529 |
| Random effect | | | | |
| Order (Intercept) | 0 | 0 | | |

**Supplementary Table 2: Pairwise differences in the proportion of individuals changing their behavioural state in at the start of calls in experiment 1. p<0.05 are bolded**

**ANOVA Type II Wald Chisquare test**

**Treatment: $\chi2 = 16.03$; df = 3; p-value < 0.01**

Sex: $\chi2 = 0.396$; df = 1; p-value = 0.529

Pairwise comparisons between treatment

| Group 1 | Group 2 | p-value | Adj. p-value |
|---|---|---|---|
| **Rufous-tailed Jacamar** | **Eastern Kingbird** | **0.0007** | **0.004** |
| Rufous-tailed Jacamar | Toucan | 0.104 | 0.624 |
| **Rufous-tailed Jacamar** | **Greenhouse noise** | **0.001** | **0.006** |
| Eastern Kingbird | Toco Toucan | 0.11 | 0.7 |
| Eastern Kingbird | Greenhouse noise | 1 | 1 |
| Toco Toucan | Greenhouse noise | 0.10 | 0.61 |

**Supplementary Table 3: GLMM results on the effect of treatment (calls) and sex on proportion of butterflies changing their behaviour at the end of calls in experiment 1. p<0.05 are bolded**

| Fixed effect | Estimate | SE | z value | Pr (>\|z\|) |
|---|---|---|---|---|
| **Intercept** | **-2.46** | **0.64** | **-3.79** | **<0.001** |
| **Treatment (Rufous-tailed Jacamar)** | **2.22** | **0.67** | **3.30** | **<0.001** |
| Treatment (Eastern Kingbird) | 0.44 | 0.75 | 0.59 | 0.554 |
| **Treatment (Toco Toucan)** | **1.62** | **0.69** | **2.34** | **<0.05** |
| Sex (male) | 0.13 | 0.37 | 0.37 | 0.709 |
| Random effect | | | | |
| Order (Intercept) | 0.06 | 0.25 | | |

**Supplementary Table 4: Pairwise differences in the proportion of individuals changing their behavioural state in at the end of calls in experiment 1. p<0.05 are bolded**

**ANOVA Type II Wald Chisquare test**

**Treatment: $\chi 2 = 17.47$; df = 3; p-value < 0.001**

Sex: $\chi 2 = 0.139$; df = 1; p-value = 0.709

Pairwise comparisons between treatment

| Group 1 | Group 2 | p-value | Adj. p-value |
|---|---|---|---|
| **Rufous-tailed Jacamar** | **Eastern Kingbird** | **0.001** | **0.006** |
| Rufous-tailed Jacamar | Toucan | 0.197 | 1 |
| **Rufous-tailed Jacamar** | **Greenhouse noise** | **0.0002** | **0.001** |
| Eastern Kingbird | Toco Toucan | 0.07 | 0.451 |
| Eastern Kingbird | Greenhouse noise | 0.724 | 1 |
| Toco Toucan | Greenhouse noise | 0.015 | 0.093 |

**Supplementary Table 5: Loadings of each behaviour in Principal Component (PC) composite variables for males in a minute before, during, and after calls in experiment 1**

| Behaviour | PC1 | PC2 | PC3 |
|---|---|---|---|
| Rest | 0.673 | 0.112 | 0.115 |
| Fly | 0.170 | 0.100 | 0.689 |
| Bask | 0.602 | 0.294 | 0.200 |
| Flutter | 0.195 | 0.674 | 0.088 |
| Court | 0.053 | 0.015 | 0.600 |
| Copulate | 0.095 | 0.254 | 0.009 |
| Walk | 0.169 | 0.602 | 0.140 |
| Antenna wiggle | 0.253 | 0.080 | 0.281 |
| Sitting near each other | 0.108 | 0.025 | 0.061 |
| % Variance explained | 22.28 | 17.05 | 14.43 |
| % Total variance explained | 22.28 | 39.33 | 53.77 |

**Supplementary Table 6: Loadings of each behaviour in Principal Component (PC) composite variables for females in a minute before, during and after calls in experiment 1**

| Behaviour | PC1 | PC2 | PC3 |
|---|---|---|---|
| Rest | 0.605 | 0.269 | 0.208 |
| Fly | 0.193 | 0.236 | 0.576 |
| Bask | 0.490 | 0.493 | 0.136 |
| Flutter | 0.362 | 0.577 | 0.016 |
| Copulate | 0.066 | 0.162 | 0.583 |
| Walk | 0.355 | 0.512 | 0.149 |
| Antenna wiggle | 0.304 | 0.071 | 0.491 |
| Lifting abdomen | 0.028 | 0.030 | 0.037 |
| % Variance explained | 28.18 | 17.83 | 13.77 |
| % Total variance explained | 28.18 | 46.02 | 59.79 |

**Supplementary Table 7: ANOVA Post-hoc test results with PC as dependent variables and the treatments (calls), state (before, during, and after) and their interaction as response variables for males and females in a minute before, during, and after calls in experiment 1. EK= Eastern kingbird; RJ=Rufous-tailed jacamar; TT=Toco toucan; GN=Greenhouse background noise; Difference=pairwise difference; lwr=lower range; upr=upper range; padj= adjusted p-value.**

| | ♂ | | | | ♀ | | | |
|---|---|---|---|---|---|---|---|---|
| | PC1 | | | | PC1 | | | |
| Treatment | Difference | lwr | upr | padj | Difference | lwr | upr | padj |
| EK-RJ | **0.861** | **0.235** | **1.487** | **0.002** | 0.418 | -0.240 | 1.077 | 0.357 |
| TT-RJ | 0.368 | -0.264 | 1.001 | 0.434 | 0.141 | -0.525 | 0.807 | 0.947 |
| GN-RJ | **0.882** | **0.223** | **1.542** | **0.003** | **0.982** | **0.287** | **1.676** | **0.001** |
| TT-EK | -0.492 | -1.125 | 0.140 | 0.185 | -0.276 | -0.943 | 0.389 | 0.705 |
| GN-EK | 0.021 | -0.638 | 0.680 | 0.999 | 0.564 | -0.130 | 1.258 | 0.155 |
| GN-TT | 0.513 | -0.512 | 1.180 | 0.192 | **0.840** | **0.138** | **1.542** | **0.011** |
| State | | | | | | | | |
| During-Before | -0.153 | -0.661 | 0.355 | 0.757 | -0.516 | -1.051 | 0.019 | 0.061 |
| After-Before | 0.040 | -0.468 | 0.548 | 0.981 | -0.129 | -0.665 | 0.406 | 0.836 |
| After-During | 0.193 | -0.315 | 0.701 | 0.643 | 0.386 | -0.149 | 0.922 | 0.206 |
| | PC2 | | | | PC2 | | | |
| Treatment | Difference | lwr | upr | padj | Difference | lwr | upr | padj |
| EK-RJ | **-0.934** | **-1.443** | **-0.426** | **0.00002** | 0.226 | -0.307 | 0.761 | 0.690 |
| TT-RJ | **-1.108** | **-1.622** | **-0.594** | **0.000004** | **0.583** | **0.042** | **1.124** | **0.028** |
| GN-RJ | **-1.074** | **-1.610** | **-0.538** | **0.000002** | 0.154 | -0.408 | 0.717 | 0.893 |
| TT-EK | -0.173 | -0.688 | 0.340 | 0.818 | 0.356 | -0.183 | 0.897 | 0.322 |
| GN-EK | -0.139 | -0.675 | 0.396 | 0.906 | -0.072 | -0.635 | 0.490 | 0.987 |
| GN-TT | 0.033 | -0.507 | 0.575 | 0.998 | -0.429 | -0.998 | 0.140 | 0.209 |
| State | | | | | | | | |
| During-Before | **0.454** | **0.040** | **0.867** | **0.027** | 0.396 | -0.037 | 0.830 | 0.081 |
| After-Before | 0.111 | -0.301 | 0.524 | 0.799 | 0.102 | -0.332 | 0.536 | 0.844 |
| After-During | -0.342 | -0.755 | 0.070 | 0.125 | -0.294 | -0.728 | 0.140 | 0.248 |

**Supplementary Table 8: Loadings of each behaviour in Principal Component (PC) composite variables for males in 14 minutes before and after calls in experiment 1**

| Behaviour | PC1 | PC2 | PC3 |
|---|---|---|---|
| Rest | 0.465 | 0.402 | 0.242 |
| Fly | 0.276 | 0.407 | 0.350 |
| Bask | 0.334 | 0.575 | 0.147 |
| Flutter | 0.443 | 0.378 | 0.235 |
| Court | 0.117 | 0.282 | 0.599 |
| Copulate | 0.006 | 0.093 | 0.039 |
| Walk | 0.489 | 0.275 | 0.321 |
| Antenna wiggle | 0.377 | 0.182 | 0.240 |
| Sitting near each other | 0.047 | 0.016 | 0.467 |
| % Variance explained | 29.23 | 19.82 | 15.88 |
| % Total variance explained | 29.23 | 49.06 | 64.94 |

**Supplementary Table 9: Loadings of each behaviour in Principal Component (PC) composite variables for females in 14 minutes before and after calls in experiment 1**

| Behaviour | PC1 | PC2 | PC3 |
|---|---|---|---|
| Rest | 0.548 | 0.285 | 0.204 |
| Fly | 0.320 | 0.272 | 0.104 |
| Bask | 0.469 | 0.416 | 0.153 |
| Flutter | 0.388 | 0.537 | 0.026 |
| Copulate | 0.045 | 0.136 | 0.842 |
| Walk | 0.376 | 0.446 | 0.051 |
| Antenna wiggle | 0.262 | 0.332 | 0.381 |
| Sit near each other | 0.041 | 0.087 | 0.246 |
| Lifting abdomen | 0.100 | 0.215 | 0.068 |
| % Variance explained | 28.99 | 17.63 | 12.24 |
| % Total variance explained | 28.99 | 46.63 | 58.87 |

**Supplementary Table 10: Effect of treatment (Rufous-tailed Jacamar, Eastern Kingbird, Toco Toucan and Greenhouse background noise calls), state (before call and after call) and their interaction on 14 minute behaviours before and after call, and male PC1, PC2, PC3, *courtship, copulation, sit near* and female PC1, PC2, PC3, *abdomen lifting* behaviours in experiment 1.**

| ♂ | AIC | Df | F value | Pr (>F) | ♀ | AIC | Df | F value | Pr (>F) |
|---|---|---|---|---|---|---|---|---|---|
| PC1 | 644 | | | | PC1 | 635 | | | |
| Treatment | | 3 | 0.734 | 0.533 | **Treatment** | | **3** | **3.615** | **0.014** |
| State | | 1 | 0.014 | 0.907 | State | | 1 | 1.488 | 0.224 |
| Treatment*State | | 3 | 1.068 | 0.364 | Treatment*State | | 3 | 0.236 | 0.871 |
| PC2 | 580 | | | | PC2 | 560 | | | |
| Treatment | | 3 | 1.482 | 0.222 | Treatment | | 3 | 0.731 | 0.535 |
| State | | 1 | 0.380 | 0.538 | State | | 1 | 0.155 | 0.284 |
| Treatment*State | | 3 | 0.133 | 0.952 | Treatment*State | | 3 | 0.770 | 0.512 |
| PC3 | 542 | | | | PC3 | 500 | | | |
| Treatment | | 3 | 1.639 | 0.183 | Treatment | | 3 | 1.142 | 0.334 |
| State | | 1 | 0.664 | 0.416 | State | | 1 | 0.498 | 0.481 |
| Treatment*State | | 3 | 0.062 | 0.980 | Treatment*State | | 3 | 0.433 | 0.730 |
| Courtship | 1606 | | | | Copulation | 1983 | | | |
| Treatment | | 3 | 1.313 | 0.272 | Treatment | | 3 | 1.967 | 0.121 |
| State | | 1 | 1.933 | 0.166 | State | | 1 | 0 | 1 |
| Treatment*State | | 3 | 0.243 | 0.866 | Treatment*State | | 3 | 0 | 1 |
| Sitting near other | 2027 | | | | Abdomen lift | 1676 | | | |
| Treatment | | 3 | 0.953 | 0.417 | Treatment | | 3 | 0.613 | 0.608 |
| State | | 1 | 0.264 | 0.608 | State | | 1 | 0.621 | 0.432 |
| Treatment*State | | 3 | 0.221 | 0.882 | Treatment*State | | 3 | 0.979 | 0.404 |

**Supplementary Table 11: ANOVA post-hoc test results with PC as dependent variables and the treatments (calls), state (before and after) and their interaction as response variables for males and females in 14 minutes before and after calls in experiment 1. EK= Eastern kingbird; RJ=Rufous-tailed jacamar; TT=Toco toucan; GN=Greenhouse background noise; Difference= pairwise difference; lwr=lower range; upr=upper range; padj= adjusted p-value.**

| | | ♂ | | | | ♀ | | |
|---|---|---|---|---|---|---|---|---|
| | | PC1 | | | | PC1 | | |
| Treatment | Difference | lwr | upr | padj | Difference | lwr | upr | padj |
| EK-RJ | 0.160 | -0.755 | 1.076 | 0.968 | 0.064 | -0.956 | 0.826 | 0.997 |
| TT-RJ | 0.305 | -0.610 | 1.221 | 0.821 | -0.043 | -0.934 | 0.848 | 0.999 |
| GN-RJ | 0.528 | -0.435 | 1.492 | 0.486 | **0.948** | **0.010** | **1.887** | **0.046** |
| TT-EK | 0.145 | -0.759 | 1.050 | 0.975 | 0.021 | -0.859 | 0.902 | 0.999 |
| GN-EK | 0.367 | -0.586 | 1.321 | 0.748 | **1.013** | **0.084** | **1.942** | **0.026** |
| GN-TT | 0.222 | -0.731 | 1.176 | 0.929 | **0.992** | **0.063** | **1.920** | **0.031** |
| State | | | | | | | | |
| After-Before | -0.029 | -0.530 | 0.471 | 0.907 | -0.301 | -0.789 | 0.186 | 0.224 |
| | | PC2 | | | | PC2 | | |
| Treatment | Difference | lwr | upr | padj | Difference | lwr | upr | padj |
| EK-RJ | 0.202 | -0.552 | 0.956 | 0.898 | 0.177 | -0.533 | 0.887 | 0.916 |
| TT-RJ | -0.250 | -1.005 | 0.503 | 0.823 | **0.382** | **-0.328** | **1.093** | **0.502** |
| GN-RJ | -0.370 | -1.165 | 0.423 | 0.620 | 0.305 | -0.443 | 1.053 | 0.714 |
| TT-EK | -0.452 | -1.198 | 0.292 | 0.394 | 0.205 | -0.497 | 0.907 | 0.872 |
| GN-EK | -0.572 | -1.358 | 0.213 | 0.235 | 0.128 | -0.612 | 0.868 | 0.969 |
| GN-TT | -0.119 | -0.905 | 0.666 | 0.978 | -0.077 | -0.817 | 0.663 | 0.993 |
| State | | | | | | | | |
| After-Before | -0.128 | -0.542 | 0.284 | 0.538 | 0.211 | -0.177 | 0.600 | 0.284 |

**Supplementary Table 12: GLMM results on the effect of treatment (calls) and sex on proportion of butterflies changing their behaviour at the start of calls in experiment 2. p<0.05 are bolded**

| Fixed effect | Estimate | SE | z value | Pr (>\|z\|) |
|---|---|---|---|---|
| **Intercept** | **-2.03** | **0.48** | **-4.18** | **<0.0001** |
| Treatment (Eastern Kingbird) | 0.57 | 0.57 | 1.01 | 0.312 |
| **Treatment (Tropical Kingbird)** | **1.16** | **0.54** | **2.15** | **<0.05** |
| Sex (male) | 0.08 | 0.42 | 0.21 | 0.832 |
| Random effect | | | | |
| Order (Intercept) | 6.9e-15 | 8.3e-8 | | |

**Supplementary Table 13: Pairwise differences in the proportion of butterflies changing their behavioural state in response to the start of calls in experiment 2**

**ANOVA Type II Wald Chisquare test**

Treatment: $\chi2 = 4.807$; df = 2; p-value = 0.09

Sex: $\chi2 = 0.044$; df = 1; p-value = 0.832

Pairwise comparisons between treatment

| Group 1 | Group 2 | p-value | Adj. p-value |
|---|---|---|---|
| Tropical Kingbird | Eastern Kingbird | 0.336 | 1 |
| Tropical Kingbird | Greenhouse noise | 0.042 | 0.128 |
| Eastern Kingbird | Greenhouse noise | 0.402 | 1 |

**Supplementary Table 14: GLMM results of the effect of treatment (calls) and sex on proportion of butterflies changing their behaviour at the end of calls in experiment 2. p<0.05 are bolded**

| Fixed effect | Estimate | SE | z value | Pr (>\|z\|) |
|---|---|---|---|---|
| **Intercept** | **-1.55** | **0.41** | **-3.70** | **<0.001** |
| Treatment (Eastern Kingbird) | 0.47 | 0.49 | 0.95 | 0.340 |
| Treatment (Tropical Kingbird) | 0.68 | 0.48 | 1.41 | 0.157 |
| Sex (male) | 0.07 | 0.39 | 0.19 | 0.844 |
| Random effect | | | | |
| Order (Intercept) | 0 | 0 | | |

**Supplementary table 15: Pairwise differences in the proportion of males and females changing their behavioural state in response to the end of calls in experiment 2**

**ANOVA Type II Wald Chisquare test**

Treatment: $\chi 2 = 2.037$; df = 2; p-value = 0.361

Sex: $\chi 2 = 0.038$; df = 1; p-value = 0.844

Pairwise comparisons between treatment

| Group 1 | Group 2 | p-value | Adj. p-value |
|---|---|---|---|
| Tropical Kingbird | Eastern Kingbird | 0.817 | 1 |
| Tropical Kingbird | Greenhouse noise | 0.231 | 0.693 |
| Eastern Kingbird | Greenhouse noise | 0.459 | 1 |

**Supplementary Table 16: GLMM results of the effect of treatment (calls) and sex on proportion of butterflies changing their behaviour in response to calls in experiment 2. p<0.05 are bolded**

| Fixed effect | Estimate | SE | z value | Pr (>\|z\|) |
|---|---|---|---|---|
| **Intercept** | **-1.31** | **0.38** | **-3.43** | **<0.001** |
| **Treatment (Eastern Kingbird)** | **0.89** | **0.44** | **2.00** | **0.044** |
| **Treatment (Tropical Kingbird)** | **0.98** | **0.44** | **2.20** | **0.027** |
| Sex (male) | 0.31 | 0.35 | 0.88 | 0.376 |
| Random effect | | | | |
| Order (Intercept) | 0 | 0 | | |

**Supplementary table 17: Pairwise differences in the proportion of individuals changing their behavioural state in response to the calls in experiment 2**

**ANOVA Type II Wald Chisquare test**

Treatment: $\chi 2 = 5.756$; df = 2; p-value = 0.056

Sex: $\chi 2 = 0.783$; df = 1; p-value = 0.376

Pairwise comparisons between treatment

| Group 1 | Group 2 | p-value | Adj. p-value |
|---|---|---|---|
| Tropical Kingbird | Eastern Kingbird | 1 | 1 |
| Tropical Kingbird | Greenhouse noise | 0.032 | 0.096 |
| Eastern Kingbird | Greenhouse noise | 0.052 | 0.158 |

**Supplementary Table 18: Loadings of each behaviour in Principal Component (PC) composite variables for males in a minute before, during and after calls in experiment 2**

| Behaviour | PC1 | PC2 | PC3 |
|---|---|---|---|
| Rest | 0.537 | 0.514 | 0.139 |
| Fly | 0.292 | 0.017 | 0.413 |
| Bask | 0.274 | 0.401 | 0.400 |
| Flutter | 0.485 | 0.438 | 0.224 |
| Court | 0.258 | 0.026 | 0.332 |
| Copulate | 0.033 | 0.463 | 0.622 |
| Walk | 0.493 | 0.400 | 0.249 |
| Antenna wiggle | 0.056 | 0.074 | 0.194 |
| % Variance explained | 23.84 | 19.50 | 16.99 |
| % Total variance explained | 23.84 | 43.35 | 60.34 |

**Supplementary Table 19: Loadings of each behaviour in Principal Component (PC) composite variables for females in a minute before, during and after calls in experiment 2**

| Behaviour | PC1 | PC2 | PC3 |
|---|---|---|---|
| Rest | 0.427 | 0.609 | 0.087 |
| Fly | 0.277 | 0.017 | 0.141 |
| Bask | 0.365 | 0.288 | 0.556 |
| Flutter | 0.506 | 0.314 | 0.319 |
| Copulate | 0.121 | 0.598 | 0.550 |
| Walk | 0.478 | 0.275 | 0.420 |
| Antenna wiggle | 0.307 | 0.045 | 0.278 |
| Lifting abdomen | 0.106 | 0.099 | 0.045 |
| % Variance explained | 27.03 | 19.39 | 15.16 |
| % Total variance explained | 27.03 | 46.42 | 61.58 |

**Supplementary Table 20: Effect of treatment (Tropical Kingbird, Eastern Kingbird, and Greenhouse background noise calls), state (before, during, and after call) and their interaction on male PC1, PC2, PC3, *courtship, sit near* and female PC1, PC2, PC3, *copulation*, *abdomen lifting* behaviours in experiment 2. p<0.05 bolded. For male *sit near* behaviour, there were zero occurrences.**

| ♂ | AIC | Df | F value | Pr (>F) | ♀ | AIC | Df | F value | Pr (>F) |
|---|---|---|---|---|---|---|---|---|---|
| PC1 | 760 | | | | PC1 | 784 | | | |
| Treatment | | 2 | 0.062 | 0.940 | Treatment | | 2 | 0.599 | 0.550 |
| State | | 2 | 0.440 | 0.645 | State | | 2 | 1.249 | 0.289 |
| Treatment*State | | 4 | 0.249 | 0.910 | Treatment*State | | 4 | 0.172 | 0.952 |
| PC2 | 713 | | | | PC2 | 712 | | | |
| Treatment | | 2 | 2.531 | 0.082 | Treatment | | 2 | 2.361 | 0.096 |
| State | | 2 | 0.667 | 0.514 | State | | 2 | 0.207 | 0.813 |
| Treatment*State | | 4 | 0.048 | 0.995 | Treatment*State | | 4 | 0.167 | 0.954 |
| PC3 | 683 | | | | PC3 | 661 | | | |
| **Treatment** | | **2** | **3.157** | **0.044** | Treatment | | 2 | 1.075 | 0.343 |
| State | | 2 | 0.015 | 0.985 | State | | 2 | 0.385 | 0.681 |
| Treatment*State | | 4 | 0.205 | 0.935 | Treatment*State | | 4 | 0.453 | 0.770 |
| Courtship | 1021 | | | | Copulation | 1929 | | | |
| Treatment | | 2 | 2.064 | 0.130 | **Treatment** | | **2** | **3.413** | **0.034** |
| State | | 2 | 0.292 | 0.747 | State | | 2 | 0 | 1 |
| Treatment*State | | 4 | 0.731 | 0.572 | Treatment*State | | 4 | 0 | 1 |
| Sitting near other | NA | | | | Abdomen lift | 1034 | | | |
| Treatment | | 2 | 0 | 0 | Treatment | | 2 | 1.279 | 0.280 |
| State | | 2 | 0 | 0 | State | | 2 | 1.588 | 0.207 |
| Treatment*State | | 4 | 0 | 0 | Treatment*State | | 4 | 1.292 | 0.274 |

**Supplementary Table 21: ANOVA post-hoc test results with PC as dependent variables and the treatments (calls), state (before, during, and after) and their interaction as response variables for males and females in a minute before, during, and after calls in experiment 2. EK= Eastern kingbird; TK=Tropical kingbird; GN=Greenhouse background noise; Difference= pairwise difference; lwr=lower range; upr=upper range; padj= adjusted p-value.**

| | ♂ | | | | ♀ | | | |
|---|---|---|---|---|---|---|---|---|
| | PC1 | | | | PC1 | | | |
| Treatment | Difference | lwr | upr | padj | Difference | lwr | upr | padj |
| EK-TK | -0.055 | -0.620 | 0.508 | 0.970 | -0.120 | -0.717 | 0.477 | 0.883 |
| GN-TK | -0.080 | -0.633 | 0.472 | 0.936 | 0.150 | -0.435 | 0.735 | 0.817 |
| GN-EK | -0.025 | -0.578 | 0.527 | 0.993 | 0.270 | -0.315 | 0.856 | 0.520 |
| State | | | | | | | | |
| During-Before | -0.213 | -0.773 | 0.339 | 0.627 | -0.078 | -0.667 | 0.511 | 0.947 |
| After-Before | -0.144 | -0.701 | 0.411 | 0.812 | -0.373 | -0.962 | 0.215 | 0.294 |
| After-During | 0.072 | -0.483 | 0.629 | 0.949 | -0.295 | -0.884 | 0.293 | 0.463 |
| | PC2 | | | | PC2 | | | |
| Treatment | Difference | lwr | upr | padj | Difference | lwr | upr | padj |
| EK-TK | -0.409 | -0.914 | 0.095 | 0.136 | 0.028 | -0.475 | 0.533 | 0.990 |
| GN-TK | -0.417 | -0.912 | 0.077 | 0.116 | -0.376 | -0.870 | 0.118 | 0.173 |
| GN-EK | -0.007 | -0.502 | 0.486 | 0.999 | -0.404 | -0.899 | 0.089 | 0.131 |
| State | | | | | | | | |
| During-Before | 0.175 | -0.322 | 0.673 | 0.684 | -0.002 | -0.499 | 0.495 | 0.999 |
| After-Before | 0.234 | -0.263 | 0.731 | 0.508 | 0.116 | -0.381 | 0.613 | 0.845 |
| After-During | 0.058 | -0.438 | 0.556 | 0.957 | 0.118 | -0.379 | 0.615 | 0.840 |

**Supplementary Table 22: Loadings of each behaviour in Principal Component (PC) composite variables for males in 14 minutes before and after calls in experiment 2**

| Behaviour | PC1 | PC2 | PC3 |
|---|---|---|---|
| Rest | 0.013 | 0.435 | 0.374 |
| Fly | 0.325 | 0.102 | 0.225 |
| Bask | 0.047 | 0.233 | 0.631 |
| Flutter | 0.480 | 0.052 | 0.378 |
| Court | 0.538 | 0.207 | 0.190 |
| Copulate | 0.244 | 0.636 | 0.039 |
| Walk | 0.262 | 0.203 | 0.401 |
| Antenna wiggle | 0.014 | 0.455 | 0.217 |
| Sit near each other | 0.491 | 0.212 | 0.141 |
| % Variance explained | 24.60 | 20.21 | 17.82 |
| % Total variance explained | 24.60 | 44.81 | 62.64 |

**Supplementary Table 23: Loadings of each behaviour in Principal Component (PC) composite variables for females in 14 minutes before and after calls in experiment 2**

| Behaviour | PC1 | PC2 | PC3 |
|---|---|---|---|
| Rest | 0.210 | 0.708 | 0.151 |
| Fly | 0.350 | 0.029 | 0.221 |
| Bask | 0.430 | 0.234 | 0.262 |
| Flutter | 0.424 | 0.180 | 0.462 |
| Copulate | 0.278 | 0.603 | 0.319 |
| Walk | 0.452 | 0.090 | 0.429 |
| Antenna wiggle | 0.355 | 0.062 | 0.360 |
| Sit near each other | 0.078 | 0.175 | 0.468 |
| Lifting abdomen | 0.227 | 0.047 | 0.097 |
| % Variance explained | 29.96 | 18.61 | 13.99 |
| % Total variance explained | 29.96 | 48.58 | 62.58 |

**Supplementary Table 24: Effect of treatment (Rufous-tailed Jacamar, Eastern Kingbird, Toco Toucan and Greenhouse background noise calls), state (before call and after call) and their interaction on 14 minute behaviors before and after call, and male PC1, PC2, PC3, *courtship, copulation, sit near* and female PC1, PC2, PC3, *abdomen lifting* behaviours in experiment 2.**

| ♂ | AIC | Df | F value | Pr (>F) | ♀ | AIC | Df | F value | Pr (>F) |
|---|---|---|---|---|---|---|---|---|---|
| PC1 | 527 | | | | PC1 | 555 | | | |
| Treatment | | 2 | 0.558 | 0.574 | Treatment | | 2 | 1.184 | 0.309 |
| State | | 1 | 0.288 | 0.592 | State | | 1 | 0.002 | 0.961 |
| Treatment*State | | 2 | 0.840 | 0.434 | Treatment*State | | 2 | 0.010 | 0.990 |
| PC2 | 497 | | | | PC2 | 486 | | | |
| Treatment | | 2 | 1.979 | 0.142 | Treatment | | 2 | 1.363 | 0.259 |
| State | | 1 | 1.212 | 0.273 | State | | 1 | 0.463 | 0.497 |
| Treatment*State | | 2 | 0.025 | 0.975 | Treatment*State | | 2 | 0.280 | 0.756 |
| PC3 | 483 | | | | PC3 | 443 | | | |
| Treatment | | 2 | 0.009 | 0.991 | Treatment | | 2 | 2.640 | 0.075 |
| State | | 1 | 0.060 | 0.808 | State | | 1 | 0.645 | 0.423 |
| Treatment*State | | 2 | 0.391 | 0.677 | Treatment*State | | 2 | 0.573 | 0.565 |
| Courtship | 1436 | | | | Copulation | 2028 | | | |
| Treatment | | 2 | 0.492 | 0.612 | Treatment | | 2 | 2.821 | 0.063 |
| State | | 1 | 0.332 | 0.565 | State | | 1 | 0.838 | 0.361 |
| Treatment*State | | 2 | 0.829 | 0.439 | Treatment*State | | 2 | 0.112 | 0.894 |
| Sitting near other | 1148 | | | | Abdomen lift | 1456 | | | |
| Treatment | | 2 | 0.115 | 0.891 | Treatment | | 2 | 0.101 | 0.904 |
| State | | 1 | 1.041 | 0.309 | State | | 1 | 0.002 | 0.966 |
| Treatment*State | | 2 | 0.268 | 0.765 | Treatment*State | | 2 | 0.554 | 0.576 |

**Supplementary Table 25: ANOVA post-hoc test results with PC as dependent variables and the treatments (calls), state (before and after) and their interaction as response variables for males and females in 14 minutes before, during, and after calls in experiment 2. EK= Eastern kingbird; TK=Tropical kingbird; GN=Greenhouse background noise; Difference= pairwise difference; lwr=lower range; upr=upper range; padj= adjusted p-value.**

| | | ♂ | | | | ♀ | | |
|---|---|---|---|---|---|---|---|---|
| | | PC1 | | | | PC1 | | |
| Treatment | Difference | lwr | upr | padj | Difference | lwr | upr | padj |
| EK-TK | 0.059 | -0.683 | 0.802 | 0.980 | -0.057 | -0.879 | 0.763 | 0.984 |
| GN-TK | 0.304 | -0.423 | 1.032 | 0.584 | 0.418 | -0.387 | 1.223 | 0.437 |
| GN-EK | 0.244 | -0.483 | 0.972 | 0.705 | 0.476 | -0.329 | 1.281 | 0.343 |
| State | | | | | | | | |
| After-Before | -0.135 | -0.634 | 0.363 | 0.592 | 0.013 | -0.583 | 0.565 | 0.961 |
| | | PC2 | | | | PC2 | | |
| Treatment | Difference | lwr | upr | padj | Difference | lwr | upr | padj |
| EK-TK | -0.290 | -0.958 | 0.378 | 0.560 | -0.379 | -1.024 | 0.264 | 0.345 |
| GN-TK | -0.549 | -1.204 | 0.105 | 0.118 | -0.391 | -1.023 | 0.239 | 0.308 |
| GN-EK | -0.259 | -0.914 | 0.395 | 0.615 | -0.012 | -0.643 | 0.619 | 0.998 |
| State | | | | | | | | |
| After-Before | -0.249 | -0.698 | 0.199 | 0.272 | -0.149 | -0.582 | 0.284 | 0.497 |
