## supplementary material 2 for "Behavioural changes in aposematic *Heliconius melpomene* butterflies in response to their predatory bird calls"

#Code to extract number of seconds spent performing each behavior and add them using a .csv file extracted from SpectatorGO
 
#! /usr/bin/env python3
import sys      ##imports system files 
import csv      ## imports syntax to read/write csv files 
from datetime import datetime  ## importing datetime syntax from datetime library. datetime allows to convert text date into datetime format 
from datetime import timedelta ## importing timedelta 
start=()    ## creating a variable called start
stop=()     ## creating a variable called stop 
behav=()    ## creating a variable called behav
total_behav={}  ##creating a dictionary called total_behav

with open('file_name') as csvfile:             ## opens the csv file to be read. The file name needs to be manually changed for each file
    readCSV=csv.reader(csvfile, delimiter=',')              ## reads the csv file wherein deliminter is ","
    next(readCSV)                                           ## skips header/ first row 
    next(readCSV)                                           ## skips the next row 
    count=1                                                 ## sets counter to 1
    for row in readCSV:                                     ## defines for loop to loop through each row and starts reading each row  
        #print (row)                                        ## can be used to verify the correct row is being used to pick up data 
        start=row[1]                                        ## assigns value in second column for the row being read to variable "start'
        stop=row[2]                                         ## assigns value in third column for the row being read to variable "stop"
        behav=row[4]                                        ## assigns value in fifth column for the row being read to variable "behav"
        #print (start,stop,behav)                           ## this can be used to confirm the correct values were identified by the syntax
        if behav=="Flutter" or behav=="Ant Wiggle":         ## if statement to filter out "ant wiggle" and "flutter" behaviours
            continue
        start_str=str(start)                                ## assigns the value of "start" as a string to "start_str"
        stop_str=str(stop)                                  ## assigns the value of "stop" as a string to "stop_str"
        start_time=datetime.strptime(start_str, '%H:%M:%S') ## converts value in "start_str" from string format to datetime and assigns it to "start_time"
        print (start_time)                                  ## prints "start_time" variable which should be in datetime formate
        stop_time=datetime.strptime(stop_str, '%H:%M:%S')   ## converts value in "stop_str" from string format to datetime and assigns it to "stop_time"
        print (stop_time)                                   ## prints "stop_time" variable which should be in datetime formate
        tdelta=stop_time-start_time                         ## calculates the time difference "stop_time" and "start_time"
        #print (start_time,stop_time,tdelta,behav)          ## can be used to print all variables needed - start time, stop time, time difference and behaviour 
        if behav in total_behav:                            ## if statement that says "if the contents of the variable "behav is existing in the dictionary "total_behv""
            new_value=total_behav.get(behav)+ tdelta        ## computes a new value for the existing key. The exisitng key would be behaviour
            total_behav.update({behav:new_value})           ## updates the dictionary with newly computed value 
        else:
            total_behav.update({behav:tdelta})              ## if the behaviour didn't already exist in the dictionary, an item is added with the behaviour and respective time

for key,value in total_behav.items():                       ## looping through the final dictionary 
    value_seconds=total_behav.get(key).total_seconds()      ## creating a variable "value_seconds" and assigns the time associated with a behaviour in seconds to the variable
    total_behav.update({key:value_seconds})                 ## updates the dictionary behaviours with the time in seconds 

#with open('file_name_Output.csv', 'w') as csv_file: #Change for the file
 #   writer = csv.writer(csv_file)
  #  writer.writerow(["Behaviour","s"])
   # for key, value in total_behav.items():
    #   writer.writerow([key, value])

print (total_behav)                                         ## prints the final dictionary containing each behaviour apart from ant wiggle and flutter along with their respective total time in seconds
